# Gene-level heritability analysis explains the polygenic architecture of cancer

**DOI:** 10.1101/599753

**Authors:** Viola Fanfani, Luca Citi, Adrian L. Harris, Francesco Pezzella, Giovanni Stracquadanio

## Abstract

Genome-wide association studies (GWAS) have found hundreds of single nucleotide polymorphisms (SNPs) associated with increased risk of cancer. However, the amount of heritable risk explained by these variants is limited, thus leaving most of cancer heritability unexplained.

Recent studies have shown that genomic regions associated with specific biological functions explain a large proportion of the heritability of many traits. Since cancer is mostly triggered by aberrant genes function, we hypothesised that SNPs located in protein-coding genes could explain a significant proportion of cancer heritability.

To perform this analysis, we developed a new method, called Bayesian Gene HERitability Analysis (BAGHERA), to estimate the heritability explained by all the genotyped SNPs and by those located in protein coding genes directly from GWAS summary statistics.

By applying BAGHERA to the 38 cancers reported in the UK Biobank, we identified 1, 146 genes explaining a significant amount of cancer heritability. We found these genes to be tumour suppressors directly involved in the hallmark processes controlling the transformation from normal to cancer cell; moreover, these genes also harbour somatic driver mutation for many tumours, suggesting a two-hit model underpinning tumorigenesis.

Our study provides new evidence for a functional role of SNPs in cancer and identifies new targets for risk assessment and patients’ stratification.

## 1 Introduction

Decades of research have shown that inherited genomic mutations affect the risk of individuals of developing cancer [1, 49]. In cancer syndromes, mutations in susceptibility genes, such as the *tumour protein 53* (*TP53*) [30], and the BRCA1/2 DNA Repair Associated (*BRCA1, BRCA2*) genes[34, 58], confer up to an 8-fold increase in cancer risk in first degree relatives [49]. However, these inherited mutations are rare and highly penetrant and explain only a small fraction of the relative risk for all cancers [32].

It has been hypothesized that part of cancer risk could be apportioned to high-frequency low-penetrant variants, such as single nucleotide polymorphism (SNPs). Genome-Wide Association Studies (GWAS) have been instrumental in identifying SNPs associated with increased risk of cancer in the broader population [55], including breast [13, 52, 19], prostate [51, 14], testicular [27, 57, 29], chronic lymphocytic leukaemia [12, 46, 26], acute lymphocytic leukaemia [53, 36] and several lymphomas [11, 15]. However, the vast majority of SNPs account only for a limited increase in cancer risk [49, 47] and are usually filtered out by multiple hypotheses correction procedures applied in GWAS analysis [37].

Although most SNPs have only subtle effects, there is mounting evidence suggesting that they still contribute to the risk of developing cancer [55, 3]. Recently, we have shown that low-penetrant germline mutations in p53 pathway genes can directly control cancer related processes, including p53 activity and response to chemotherapies [60]. Moreover, the Pan-Cancer Analysis of Whole Genomes (PCAWG) study found that 17% of all patients have rare germline variants associated with cancer [7]. It is now becoming apparent that quantifying the contribution of low-penetrance inherited mutations can improve our understanding of cancer risk and the aetiology of the disease.

Heritability analysis provides the statistical framework to estimate the contribution of all common SNPs to cancer risk regardless of their statistical significance [54]. The study of heritability is now becoming a crucial step in recent cancer GWAS studies and has already provided insights on the risk of developing many malignancies [41], including prostate [31], cervical [9], testicular germ cell tumour [28] and breast cancer [42, 16].

However, since the functional impact of the SNPs is context-dependent [43], it is important to quantify the amount of heritability explained by genomic regions associated with well-characterised biological functions [17, 18, 45]. For cancer, in particular, which is mostly driven by mutations in genes rather than regulatory regions, estimating the heritability of SNPs in protein-coding genes could provide novel insights into the aetiology of this disease. However, developing analytical methods for estimating heritability at the gene-level has been challenging, and current methods allow only the estimation of heritability for large functional regions or SNP categories, such as histone marks or eQTL [18, 45].

Here we developed a new method, called BAyesian Gene HERitability Analysis (BAGHERA), which implements a hierarchical Bayesian model to obtain simultaneous estimates of the heritability explained by all genotyped SNPs (genome-wide heritability) and by those in protein coding genes (gene-level heritability). BAGHERA is specifically designed to analyse traits with putative low heritability, such as cancer, and to use GWAS summary statistics rather than genotype data; this facilitates cancer heritability analysis across different studies and cancer types. We performed extensive simulations to assess whether BAGHERA was suitable to study cancer GWAS and found that our method provides robust, unbiased genome-wide heritability estimates, and simultaneously identifies genes explaining a higher proportion of heritability, while controlling the false discovery rate. Comparison with other state-of-the-art methods clearly showed that BAGHERA provides significantly more accurate heritability estimates for diseases with heritability lower than 10%.

We then used BAGHERA to analyse the 38 histologically different malignancies reported in the UK Biobank cohort [6]. Here we provide new genome-wide estimates for all cancers and a map of 1, 146 genes that have a significant contribution to the heritability of at least 1 cancer. We then showed that the vast majority of these genes are tumour suppressors and are directly involved in the hallmark processes controlling the transformation from normal to cancer cells. While we observed pleiotropy across cancers at the functional level, we did not observe pleiotropy at the gene level; this result suggests that while the functional mechanisms mediating risk are common to all cancers, the genes affecting these processes are cancer specific. Our study provides new methods to analyse GWAS data and genetic evidence of a causal role for high-frequency inherited mutations in cancer.

## 2 Results

### 2.1 Simulation results

We performed extensive testing of our method on simulated data to assess i) the robustness of genome-wide estimates for low heritability traits and ii) the false discovery rate (FDR) associated with gene-level predictions. Finally, we compared our method with state-of-the-art approaches for genome-wide and local heritability analysis. All our datasets were calibrated to simulate low heritability traits 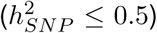, which is a reasonable assumption for cancer. Our analyses show that BAGHERA provides robust and unbiased genome-wide and gene-level heritability.

#### 2.1.1 Simulations assessing robustness of genome-wide and gene-level estimates for low heritability traits

We generated genotype data for *M* = 100, 000 SNPs of *N* = 50, 000 subjects using haplotypes of chromosome 1 from European populations (See Supplementary Methods). Here we studied two different heritability effects, denoted as dense and gene-level effects; while the former defines constant per-SNP heritability 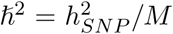, the latter condition the amount of explained heritability as a function of the gene harbouring the SNP.

Our analyses shows that BAGHERA provides robust unbiased genome-wide estimates under both dense (Fig. 1A) and gene-level heritability (Fig. 1B) models. Interestingly, while extreme values of gene-level heritability might affect genome-wide estimates, we found that BAGHERA returns robust estimates both as the median of the posterior genome-wide heritability distribution (Fig. 1A-B, green diamond marker) and the sum of single gene-level heritability contributions (Fig. Fig. 1A-B, red square marker).

**Figure 1:**
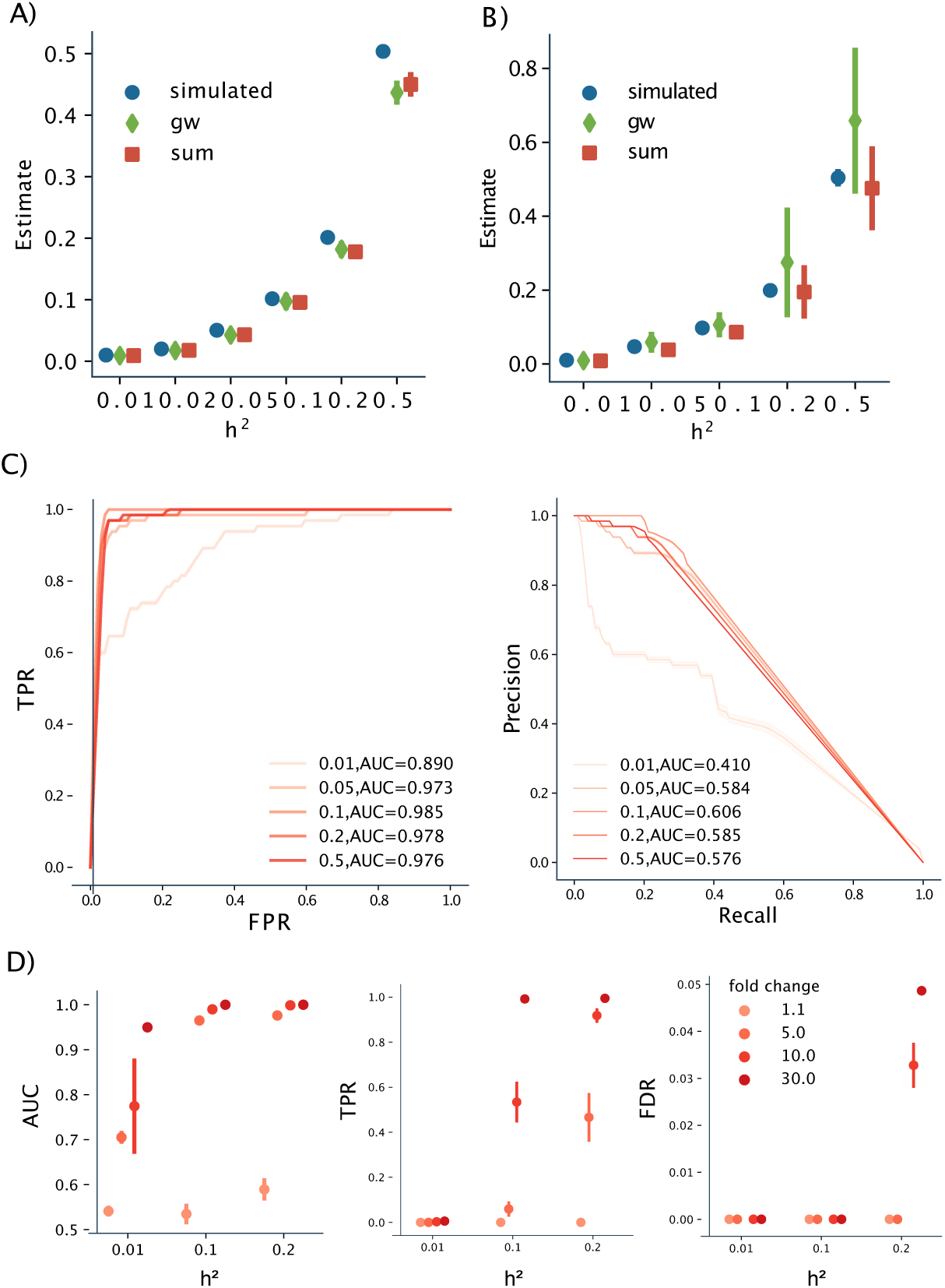
Performance on simulated data. A) Genome-wide heritability estimates for dense effects. For each value of *h*^2^, we plot the simulated value, the target, the genome-wide (gw) estimate, which is the median of the posterior of genome-wide heritability term, and the gene-level estimate which is the sum of all median gene heritability estimate (sum). B) Genome-wide heritability estimates for datasets with varying gene-level heritability. For each value of *h*^2^ we plot the simulated value, the target, the gw estimate which is the median of the prior heritability term, and the gene-level estimate which is the sum of all median gene heritability estimate C) Receiver Operator Characteristic curves and Precision Recall curves, for the performance of BAGHERA at retrieving positive genes for different values of genome-wide *h*^2^. D) Performance of BAGHERA for different values of *h*^2^ and gene level enrichment. We show the AUCs of the ROC curves, the True Positive Rate and False Discovery Rate (FDR) for *η* > 0.99. A-B-C show the performance on simulated genotype data on chromosome 1. D is showing the performance on the data simulated from summary statistics.

We then assessed whether BAGHERA was able to identify heritability genes, that is genes harbouring SNPs with a contribution to heritability higher than expected under a constant per-SNP heritability contribution. To do that, we selected 1% of the genes on chromosome 1 (≈ 13) as heritability genes and computed Receiver Operator Characteristic (ROC) and Precision Recall (PR) curves at varying levels of genome-wide heritability. Here we found that BAGHERA correctly identified heritability genes (Fig. 1C), although precision and recall were significantly higher for higher genome-wide heritability levels.

However, our simulated datasets have two limitations. First, they take into account *M* ≈ 100, 000 SNPs from a single chromosome, whereas more than 1M are routinely genotyped in modern studies. This was a necessary restriction to reduce the time and memory required to simulate genotypes, which is a computationally taxing task. Moreover, fine tuning gene-level heritability is not trivial with genotype data; low and high gene-level heritability enrichments produce either undetectable signal or extremely skewed statistics, while LD patterns might produce spurious signal difficult to control.

We addressed these limitations by developing new model for simulating summary statistics using only linkage disequilibrium information (see Supplementary Materials). This approach provides a tractable framework to test varying levels of heritability enrichment, reported in terms of fold-change with respect to the genome-wide estimate, and to simulate SNPs across the entire genome, rather than a single chromosome.

Here we found that BAGHERA correctly identifies heritability genes, even with fold-changes in heritability as low as *f*_*c*_ = 5 (see Fig. 1D and supplementary material), although the true positive rate was significantly lower for low heritability levels. Nonetheless, we found BAGHERA to be conservative with a low false discovery rate in all scenarios; this result suggests that our method is suitable for exploratory analyses, and that significant results are due to true biological signal.

#### 2.1.2 Comparison with state-of-the-art methods for genome-wide and local heritability estimation

To the best of our knowledge BAGHERA is the first method specifically designed to analyse low heritability traits and to provide heritability estimates at the gene level. Nonetheless, a number of methods have been proposed to estimate genome-wide and local heritability, thus we proceeded to compare our approach to state-of-the-art methods to perform these analyses.

For the genome-wide analysis, we compared our results with LD score regression (LDsc) [5]. Since LDsc is routinely applied to estimate heritability for the traits in the UK Biobank data, we directly retrieved the results for all 38 cancers. Unsurprisingly, since BAGHERA uses a similar genome-wide estimator, we found strong consensus between the estimates of the two methods (Supplementary Materials). Nonetheless, BAGHERA is more robust for low heritability traits, since our Bayesian formulation always provides correct genome-wide heritability estimates, whereas LDsc usually provides negative values.

For local heritability analysis, we compared our performances with the Heritability Estimation from Summary Statistics (HESS) method [45], by using BAGHERA to estimate the heritability of 1703 regions of the original article (See Supplementary Materials). Here we focused on breast and prostate cancer data, since they are those with higher 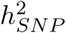 estimates and would not favour either method. We found a statistically significant correlation between the local estimates of BAGHERA and HESS, with all loci reported as significant by HESS also reported by BAGHERA. However, BAGHERA is able to identify more loci with increased heritability, while providing more robust heritability estimates for regions with low 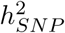 (see Supplementary Materials).

Taken together, we have shown that BAGHERA is more robust than existing methods on low heritability traits and can provide useful insights into the disease risk, being able to scale up to 15, 000 different loci across the genome.

### 2.2 Genome-wide estimates of cancer heritability in the UK Biobank

We used BAGHERA to analyse 38 cancers in the UK Biobank [6], a large-scale prospective study aiming at systematically screening and phenotyping more than 500, 000 individuals, with age ranging between 37 and 73 years.

We obtained summary statistics for *N* = 361, 194 individuals ([35], see Table 1), including subjects whose tumours were histologically characterised according to the ICD10 classification, where malignant neoplasms are identified with codes ranging from C00 to C97 (see Supplementary Material). The number of cases varies significantly across cancers, ranging from 102 individuals, for malignant neoplasm of base of tongue and other, to 9086 individual, for malignant neoplasms of the skin. In this cohort, cancer prevalence ranges between 0.29% and 2.51%, with higher estimates for common malignancies in European populations, such as breast and prostate cancer [4].

**Table 1:** **Genome-wide heritability of the** 38 **cancers in the UK BioBank**. For each cancer, we report the number of cases, the prevalence in the cohort, the average *χ*^2^ of the GWAS analysis 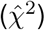, the genome-wide estimates of heritability, both on the observed 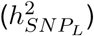 and the liability 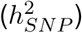 scale, and the number of heritability genes (HG) reported by BAGHERA as significant for *η* > 0.99. In bold, we denote the cancers used to build the cancer heritability genes (CHGs) panel.

| ICD10 | Malignancy | Cases | Prevalence | $\hat{\chi}^2$ | $h_{SNP}^2$ | $h_{SNP_L}^2$ | HG |
| --- | --- | --- | --- | --- | --- | --- | --- |
| C44 | Other malignant neoplasms of skin | 9086 | 0.0252 | 1.1408 | 0.0341 | 0.2422 | 422 |
| C50 | **Malignant neoplasm of breast** | 8304 | 0.0230 | 1.0869 | 0.0170 | 0.1285 | 267 |
| C61 | **Malignant neoplasm of prostate** | 4342 | 0.0120 | 1.0765 | 0.0191 | 0.2320 | 271 |
| C18 | **Malignant neoplasm of colon** | 2226 | 0.0062 | 1.0399 | 0.0070 | 0.1416 | 33 |
| C43 | **Malignant melanoma of skin** | 1672 | 0.0046 | 1.0288 | 0.0051 | 0.1293 | 52 |
| C15 | **Malignant neoplasm of oesophagus** | 519 | 0.0014 | 1.0236 | 0.0035 | 0.2296 | 24 |
| C67 | **Malignant neoplasm of bladder** | 1554 | 0.0043 | 1.0222 | 0.0047 | 0.1254 | 39 |
| C34 | **Malignant neoplasm of bronchus and lung** | 1427 | 0.0040 | 1.0208 | 0.0035 | 0.1010 | 17 |
| C20 | **Malignant neoplasm of rectum** | 1118 | 0.0031 | 1.0130 | 0.0031 | 0.1091 | 15 |
| C62 | **Malignant neoplasm of testis** | 221 | 0.0006 | 1.0120 | 0.0024 | 0.3158 | 29 |
| C71 | **Malignant neoplasm of brain** | 368 | 0.0010 | 1.0116 | 0.0030 | 0.2578 | 19 |
| C45 | **Mesothelioma** | 150 | 0.0004 | 1.0110 | 0.0012 | 0.2213 | 5 |
| C91 | **Lymphoid leukaemia** | 349 | 0.0010 | 1.0109 | 0.0018 | 0.1646 | 11 |
| C02 | **Malignant neoplasm of other and unspecified parts of tongue** | 152 | 0.0004 | 1.0106 | 0.0013 | 0.2475 | 23 |
| C16 | **Malignant neoplasm of stomach** | 388 | 0.0011 | 1.0106 | 0.0010 | 0.0868 | 12 |
| C83 | **Diffuse non-Hodgkin's lymphoma** | 587 | 0.0016 | 1.0104 | 0.0014 | 0.0824 | 14 |
| C82 | **Follicular (nodular) non-Hodgkin's lymphoma** | 320 | 0.0009 | 1.0101 | 0.0031 | 0.3059 | 21 |
| C90 | Multiple myeloma and malignant plasma cell neoplasms | 401 | 0.0011 | 1.0092 | 0.0013 | 0.1020 | 15 |
| C56 | Malignant neoplasm of ovary | 693 | 0.0019 | 1.0063 | 0.0012 | 0.0616 | 13 |
| C54 | Malignant neoplasm of corpus uteri | 988 | 0.0027 | 1.0063 | 0.0008 | 0.0295 | 14 |
| C48 | Malignant neoplasm of retroperitoneum and peritoneum | 122 | 0.0003 | 1.0053 | 0.0009 | 0.2064 | 5 |
| C64 | Malignant neoplasm of kidney except renal pelvis | 701 | 0.0019 | 1.0043 | 0.0009 | 0.0455 | 10 |
| C01 | Malignant neoplasm of base of tongue | 102 | 0.0003 | 1.0043 | 0.0014 | 0.3596 | 10 |
| C73 | Malignant neoplasm of thyroid gland | 278 | 0.0008 | 1.0042 | 0.0011 | 0.1254 | 13 |
| C49 | Malignant neoplasm of other connective and soft tissue | 222 | 0.0006 | 1.0040 | 0.0017 | 0.2229 | 28 |
| C80 | Malignant neoplasm without specification of site | 398 | 0.0011 | 1.0040 | 0.0016 | 0.1300 | 14 |
| C53 | Malignant neoplasm of cervix uteri | 192 | 0.0005 | 1.0039 | 0.0005 | 0.0709 | 14 |
| C22 | Malignant neoplasm of liver and intrahepatic bile ducts | 189 | 0.0005 | 1.0031 | 0.0009 | 0.1353 | 7 |
| C21 | Malignant neoplasm of anus and anal canal | 139 | 0.0004 | 1.0027 | 0.0007 | 0.1436 | 23 |
| C85 | Other and unspecified types of non-Hodgkin's lymphoma | 762 | 0.0021 | 1.0023 | 0.0013 | 0.0600 | 9 |
| C09 | Malignant neoplasm of tonsil | 162 | 0.0004 | 1.0022 | 0.0006 | 0.1009 | 5 |
| C92 | Myeloid leukaemia | 328 | 0.0009 | 1.0011 | 0.0008 | 0.0764 | 9 |
| C17 | Malignant neoplasm of small intestine | 114 | 0.0003 | 1.0007 | 0.0015 | 0.3596 | 12 |
| C19 | Malignant neoplasm of rectosigmoid junction | 498 | 0.0014 | 0.9992 | 0.0006 | 0.0390 | 10 |
| C25 | Malignant neoplasm of pancreas | 403 | 0.0011 | 0.9991 | 0.0005 | 0.0402 | 12 |
| C81 | Hodgkin's disease | 150 | 0.0004 | 0.9989 | 0.0003 | 0.0597 | 5 |
| C69 | Malignant neoplasm of eye and adnexa | 137 | 0.0004 | 0.9970 | 0.0004 | 0.0705 | 14 |
| C32 | Malignant neoplasm of larynx | 159 | 0.0004 | 0.9914 | 0.0003 | 0.0450 | 7 |

Estimating heritability from non-targeted cohorts can be challenging, due to the small prevalence of the disease. To test whether we had sufficient signal for each cancer, we reasoned that if the SNP test statistic follows a *χ*^2^ distribution with 1 degree of freedom, under the null hypothesis of no association, its expected value is *E*[*χ*^2^] = 1; thus, similar to other studies, we expected to have sufficient polygenic signal for our analysis if the average *χ*^2^ was greater than 1 [18]. Here we found the vast majority of cancers to have an average *χ*_2_ ≈ 1, with only 17 having a deviation greater than 1% from the expected value of the test statistic. We also did not consider cancers assigned to other malignant neoplasm of the skin (C44), as these usually comprise tumours of basal and squamous cells, which are mostly caused by sun exposure. Thus, we restricted our analysis to 16 cancers for which we had enough power to perform our analysis, although we also report results for the other cancers in the Supplementary Materials.

We then estimated genome-wide heritability of each cancer by computing the median of the posterior distribution of 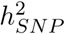 and transforming this value on to the liability scale, 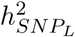, to obtain estimates independent from prevalence and comparable across malignancies. We found cancer heritability to be 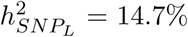 on average, ranging from 8% for non-Hodgkin’s lymphoma and up to 31% for testis (see Table 1) consistent with other available estimates for this cohort (see Supplementary Materials). While comparison between cancer heritability estimates are usually difficult across studies, due to differences in histological classification and genetic confounders, we found our heritability estimates on the liability scale to be consistent with those reported for other cohorts, in particular for breast, prostate, testes and bladder [22, 41, 31, 28]. *The heritability of testicular cancer* 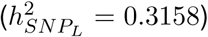 is the highest among all malignancies, consistent with the hypothesis that germline variants have stronger effects in early onset cancers. However, early onset cancers are underrepresented in the UK Biobank, since children and young adults were not enrolled in the study, and thus an accurate estimation of the correlation between age of onset and heritability is not possible. Nonetheless, it is interesting to note that many malignancies with onset in late adulthood, such as prostate or bladder, still display a significant heritable component, ranging from 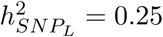 for brain tumours (age of onset: 59) to 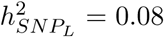 for diffuse non-Hodgkin’s lymphoma (age of onset: 60). Overall, 14 out of 16 cancers (87%) show heritability higher than 10% suggesting a consistent contribution of SNPs to the heritable risk of cancer.

### 2.3 Cancer heritability genes across 16 malignancies

We identified 783 heritability genes (*η* > 0.99), harbouring 1, 146 protein-coding genes, across 16 cancers (Fig. 2), with 53 heritability genes per malignancy on average, ranging from 5 genes in mesothelioma, to 271 genes for prostate (see Table 1, Figure 3A). It is worth noting that we are here using the term heritability genes when referring to the genomic, non-overlapping, regions tested by BAGHERA. Gene-level heritability across the selected 16 cancers has a long-tail distribution (Figure 3B), with a median 16-fold increase compared to the genome-wide estimate, ranging from 4.4-fold for the *Phosphodiesterase 4D (PDE4D)* gene to 276-fold for the *fibroblast growth factor receptor 2 (FGFR2)* gene in breast cancer. Interestingly, 87% of heritability genes show per-SNP heritability 10-fold higher than the genome-wide estimate. Only 3 genes have fold changes below 5 and more than 99% of genes with fold-changes below 10 are found in the breast and prostate datasets, which have 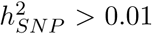. Importantly, based on our simulations for datasets with similar heritability enrichment, our set of heritability genes are expected to have a limited number of false positives.

**Figure 2:**
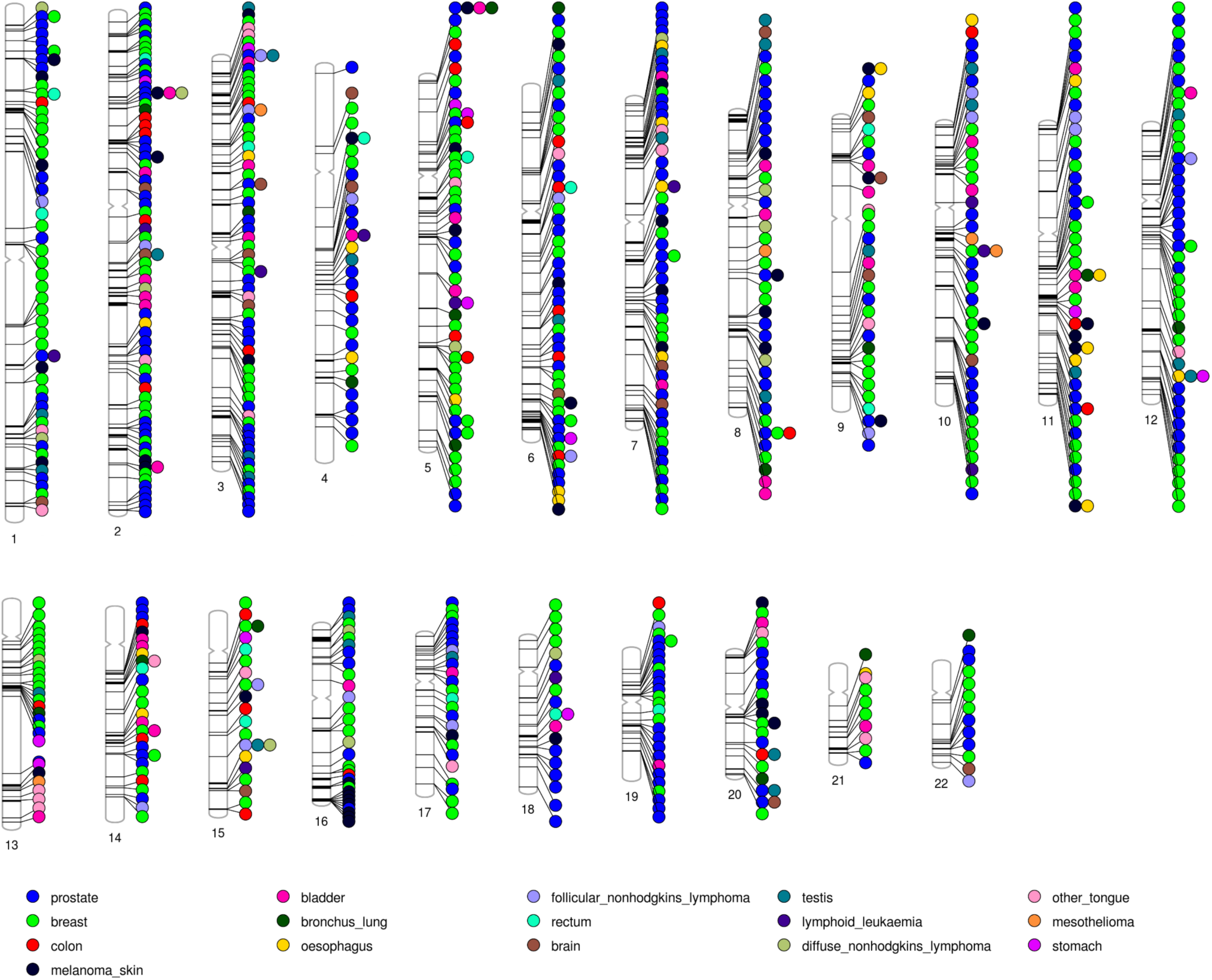
Cancer heritability genes across the human genome. For each cancer heritability gene, we report its locus and the associated cancer.

**Figure 3:**
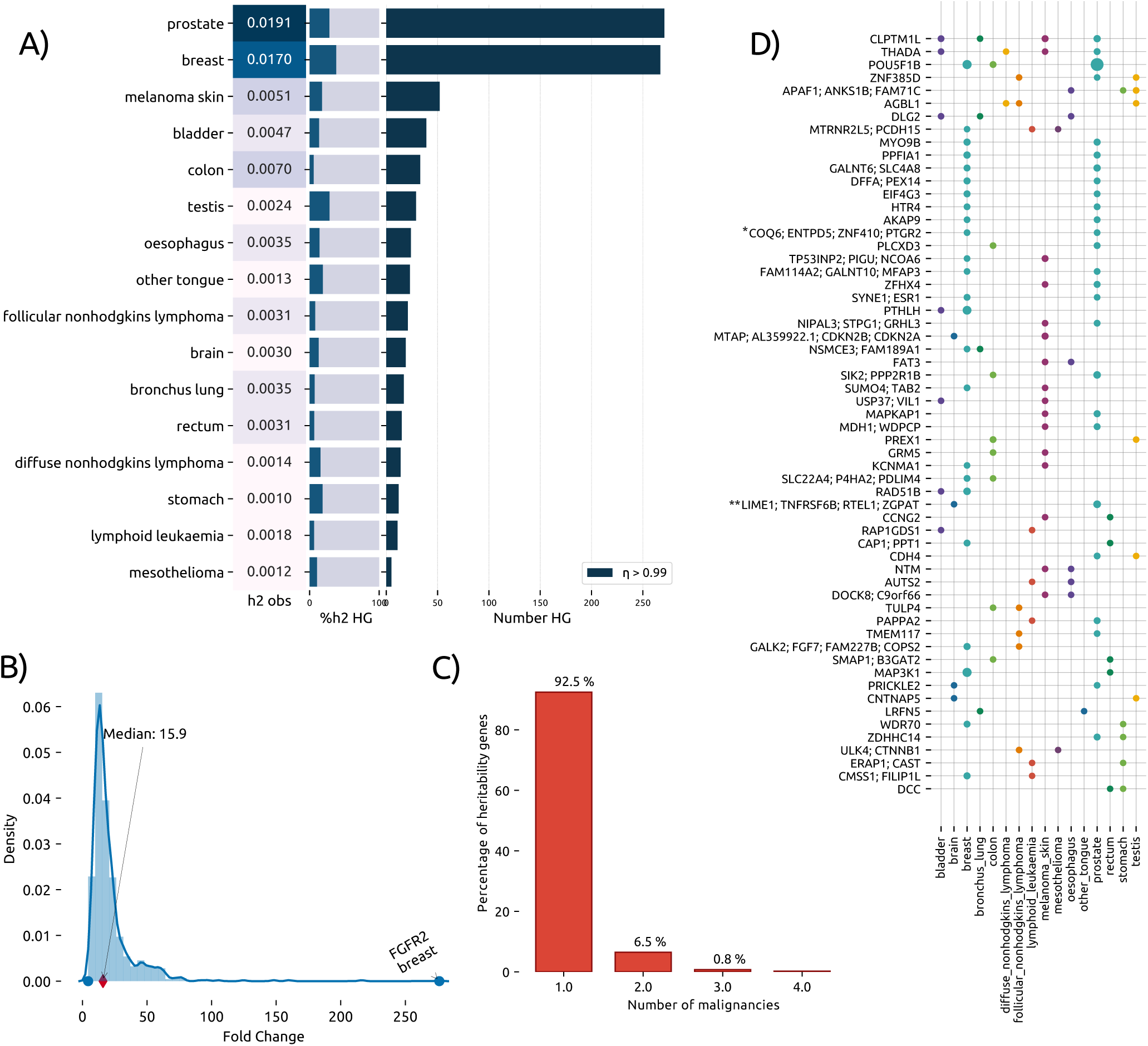
**Heritability genes across** 16 **cancers in the UK Biobank**. A) For each malignancy, we report the observed heritability (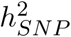, left box), the percentage of 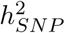 explained by heritability genes (central barplot, dark blue is the percentage explained by HGs) and the number of heritability genes (right barplot). B) Gene-level heritability density distribution across heritability genes, expressed as fold-change with respect to the genome-wide estimate. Highlighted are the top genes and the median fold-change across all cancers. C) Percentage of cancer heritability genes associated with multiple cancers. Approximately 8% of HGs are common to multiple malignancies. D) Cancer heritability genes associated with multiple cancers. We report the 59 HGs common to at least 2 cancers; here the size of the dot is proportional to the heritability enrichment of the gene in the specific cancer.

Heritability genes represent less than 1% of all the genes in the genome, but they are significantly more than those harbouring genome-wide significant SNPs (see Supplementary Materials), consistent with cancer being polygenic. Although we identified a polygenic signal, heritability genes account for up to 38% of all the heritable risk (breast cancer), suggesting that a significant amount of heritability could be explained by only few loci (Figure 3A). Consistent with our hypotheses, when we looked at the contribution of SNPs outside our protein-coding regions, we did not observe any difference compared to the genome-wide estimate.

We then tested whether heritability genes were shared among multiple cancers to identify any potential genomic hotspot for pan cancer heritability. We found that only 59 (≈ 8%) of the 783 heritability genes show a significant heritability enrichment in at least 2 cancers, and 8 (*<* 1%)in 3 or more (Figure 3C-D). This observation is consistent with results from tumour sequencing studies, which have shown that pleiotropic effects are limited to few master regulators, such as *TP53* [2]. Nonetheless, after performing literature curation, we found evidence for a cancer mediating role for 7 of the 11 unique protein coding genes found in at least 3 cancers, including 4 genes (CLPTM1L, APAF1, THADA, AGBL1) involved in apoptosis and 3 genes (PCDH15, DLG2, POU5F1B) involved in cell division, migration and tumorigenesis [44, 21]. It is important to note that the *cisplatin resistance-related protein 9* (CLPTM1L) is the heritability gene found in most cancers (4) and is one of the gene in the 5p15.33 locus (the other being TERT), which has been consistently associated with 17 different cancer types [39].

Taken together, our analysis found 784, harbouring 1, 146 protein-coding genes, having a significant contribution to the heritable risk of at least 1 cancer. We denoted these 1, 146 loci as cancer heritability genes (CHGs).s

### 2.4 Cancer heritability genes are recurrently mutated in tumours

Tumour sequencing projects, including the The Cancer Genome Atlas (TCGA) program and the Pan-Cancer Analysis of Whole Genomes (PCAWG) project, have identified a number of driver genes, which promote tumorigenesis when acquiring a somatic mutation.

There is also increasing evidence that genes harbouring germline and somatic mutations can mediate cancer phenotypes [38, 60, 47], thus we tested whether cancer heritability genes are significantly enriched among known cancer driver genes. To do that, we built a curated list of driver genes using the COSMIC Cancer Gene Census (Supplementary Table 2). We found that 60 of the 1, 146 CHGs (≈ 5%) are significantly enriched among known cancer driver genes (*OR* = 1.75, *P* : 1.3 × 10^−4^). These genes include members of the p53 pathway, such as *CDKN2A*, the *Tumour Protein 63* (*TP63*) and *MDM4 regulator of p53* (*MDM4*), as well as genes mutated across multiple types of cancer, including *FGFR2* and the *anaplastic lymphoma kinase (Ki-1)* (*ALK*) gene (Figure 4A and B).

**Figure 4:**
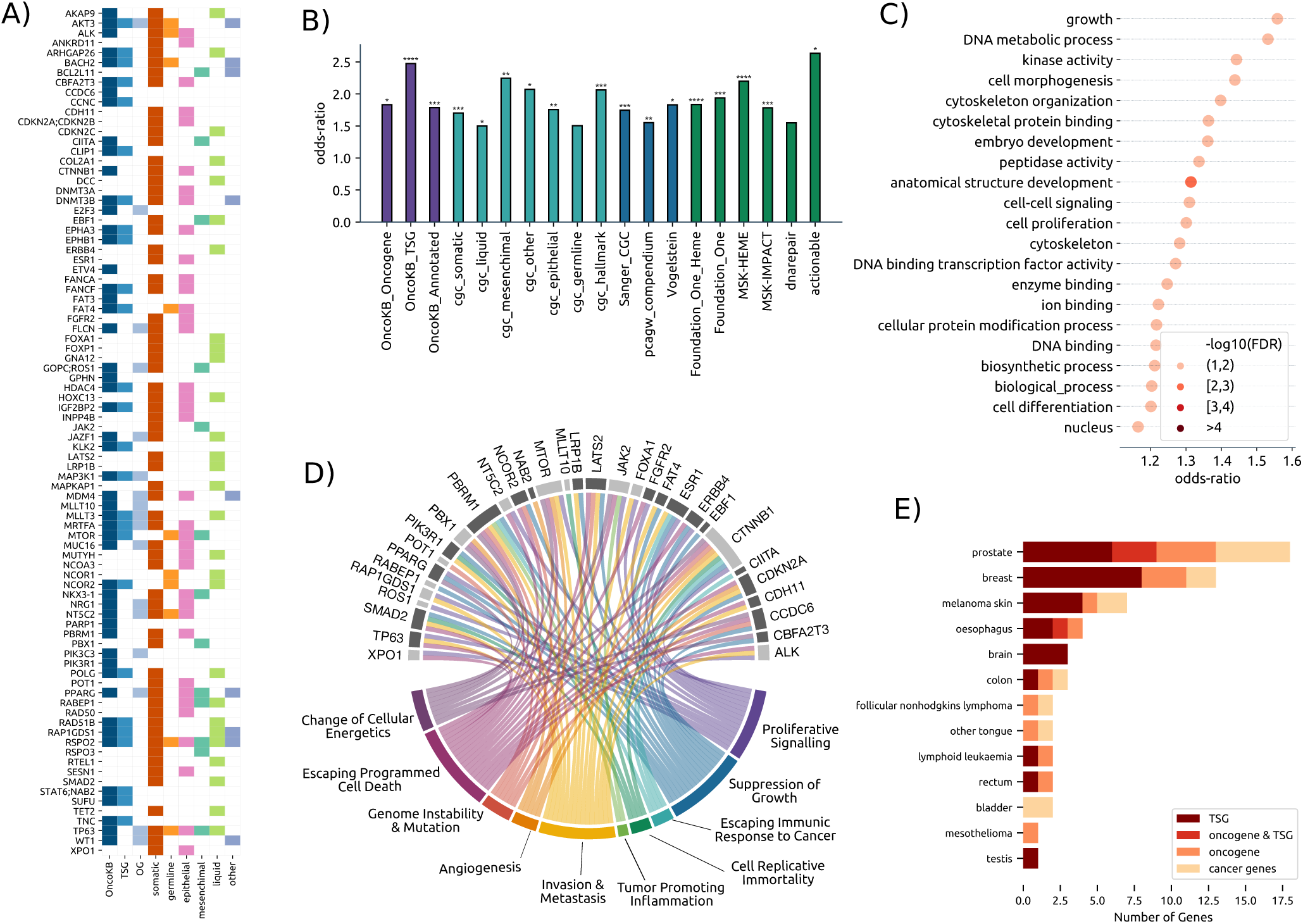
Functional characterisation of cancer heritability genes. A) List of CHGs reported as cancer driver genes across multiple annotations. With the blue hue, first three columns, we report the genes annotated by OncoKB, specifying whether they are tumour suppressors (TSG) or oncogenes (OG). With red and orange, 4-th and 5-th columns, we report the genes that are included in the COSMIC annotation as drivers and whether the reported mutation is somatic and germline. In the last four columns, we annotate each gene to the cancer type for which is denoted as driver in COSMIC. B) Enrichment of CHGs across cancer driver genes annotations; here we report OncoKB (purple), COSMIC database (light blue), different cancer driver sets (dark blue) and other sets (green) like DNA repair genes and known actionable targets. Stars indicate statistical significance, with multiple terms having *P <* 10^−4^. C) Gene Ontology enrichment analysis using Fisher’s exact test. For each significant term, we report the odds-ratio (x-axis) and − *log*_10_(FDR) (color gradients). D) CHGs associated with the hallmark of cancers; genes in darker grey are tumour suppressors. Each gene is connected to the hallmarks that it mediates. E) tumour suppressor and oncogene CHGs across cancers. For each cancer type (y-axis), we report the number of genes (x-axis) reported as tumour suppressors (TSGs) and/or oncogenes in OncoKB (colour codes).

However, the number of cancer driver genes is extremely variable across malignancies and studies, thus we tested whether the enrichment of CHGs in cancer driver genes was independent from the cancer driver gene annotation used. To do that, we collected lists of cancer driver genes from multiple studies, including the PCAWG project ([7]), the Precision Oncology Knowledge Base (OncoKB, [8]), Memorial Sloan Kettering Impact and Heme gene panels [10], and the curated list of cancer genes by Vogelstein et al. [56]. Here we found that CHGs are significantly enriched in each cancer driver gene annotation analysed, with an enrichment ranging from *OR* = 1.55 for the PCAWG annotation to 2.47 for OncoKB tumour suppressors (Supplementary Table 2). Interestingly, we did not find any enrichment of CHGs in genes carrying germline driver mutations; this is consistent with the fact that most germline driver mutations are rare, and thus are unlikely to be genotyped in GWAS studies.

Taken together, we found 60 cancer heritability genes that are also recurrently mutated in multiple tumours; this result suggests that SNPs in cancer heritability genes might affect the same biological programs altered by somatic mutations in tumours.

### 2.5 Cancer heritability genes underpin biological processes affecting tumorigenesis

Our gene-level heritability analysis identified 1, 146 loci explaining a significant proportion of the heritable risk of at least 1 cancer. We then showed that cancer heritability genes are enriched in known cancer driver genes, suggesting that loci recurrently mutated in tumours also harbour high-frequency inherited mutations that could mediate cancer risk. Thus, we hypothesised that cancer heritability genes could be involved in molecular functions and biological processes affecting tumorigenesis.

To do that, we characterized CHGs by gene ontology enrichment analysis, using the slim gene ontology for human (Supplementary Table 1). We found a statistically significant enrichment for 21 terms (Fisher’s exact test; False Discovery Rate, FDR *<* 10%, Figure 4C and Supplementary Table 1), with an average odds ratio of 1.31 and up to 1.55 for growth. CHGs are genes predominantly involved in biological processes driving cell morphogenesis (*OR* : 1.43, *P* : 2.40 × 10^−3^), differentiation (*OR* : 1.20, *P* : 7.7 × 10^−3^), which includes the *mammalian target of rapamycin (mTOR)* (*MTOR*) gene, growth (*OR* : 1.55, *P* : 2.62 × 10^−4^) and cell proliferation (*OR* : 1.3, *P* : 3.4 × 10^−3^), which includes the *Poly [ADP-ribose] polymerase 1* (*PARP1*) gene. We also observed a significant enrichment of genes associated with cytoskeleton organization (*OR* : 1.40, *P* : 2.39 × 10^−2^) and anatomical structure development (*OR* : 1.32, *P* : 6.13 × 10^−3^), which includes members of the SWI/SNF complex, such as the *AT-Rich Interaction Domain 2* (*ARID2*).

While these molecular processes drive normal cell fate, survival and proliferation, they are recurrently hijacked by cancer cells to gain growth advantage and spread through the body through metastases [48], a process that is considered an hallmark of cancer. We then tested whether cancer heritability genes are associated with any other hallmark of cancer, which are processes, common to all malignancies, controlling the transformation of normal into cancer cells [20]. These lists of biological processes include proliferative signalling, suppression of growth, escaping immunic response, cell replicative immortality, promoting inflammation, invasion and metastasis, angiogenesis, genome instability and mutation, and escaping cell death. Interestingly, we found 33 CHGs associated with at least one hallmark (*OR* : 2.062, *P* : 3 × 10^−4^). Consistent with our previous analysis, cancer heritability genes are involved in escaping cell death, mediating proliferative signalling, invasion and metastasis (Figure 4D and Supplementary Table 3). We then went further to study whether CHGs mediate these cancer processes by acting either as tumour suppressor genes (TSGs) or oncogenes (see Fig. 4E). To do that, we used the Precision Oncology Knowledge Base (OncoKB, [8]), a curated list of 519 cancer genes, including 197 tumour suppressor genes (TSGs), 148 oncogenes and other cancer genes of unknown function. We found that 27 CHGs are tumour suppressors (OR: 2.47, *P* : 7.9 × 10^−6^), whereas 17 are reported as oncogene (OR: 1.83, *P* : 0.0198) of which 4 can function both as TSG and as oncogene (Figure 4A, D and E and Supplementary tables 2 and 3). Tumour suppressor CHGs include well-known cancer driver genes, such as *CDKN2A* and *MTOR* which regulate cell growth, and DNA repair genes, such as *MUTYH* and *FANCA* [25].

Taken together, we found evidence that cancer heritability genes directly mediate processes underpinning tumorigenesis; interestingly, while we did not observe pleiotropic effects at genomic level, we found that cancer heritability genes are involved in biological processes common to all cancers. It is then conceivable that inherited mutations in genes controlling these biological programs could provide a selective advantage to cancer cells, once they acquire a driver somatic mutation. Our results suggest a functional role for cancer heritability genes consistent with a two-hit model [24]; while inherited mutations associated with oncogene activation are likely to be under purifying selection, mutations in tumour suppressor genes can be observed at higher frequency because deleterious effects are only observed upon complete loss of function.

## 3 Discussion

Our study provides new fundamental evidence demonstrating a strong contribution of high-frequency inherited mutations to the heritable risk of cancer. Here we provide a high resolution map of the heritable cancer genome consisting of 1, 146 genes showing a significant contribution to the heritable risk of 16 malignancies. We showed that these loci harbour tumour suppressors controlling growth, cell morphogenesis and proliferation, which are fundamental processes required for tumorigenesis.

Ultimately, our results support a two-hit model, where inherited mutations in tumour suppressor genes could create a favourable genetic background for tumorigenesis. It is conceivable that SNPs make normal cells more likely to evade the cell-cell contact inhibition of proliferation, to elude the anatomical constrains of their tissue and to achieve more easily independent motility in presence of other early oncogenic events. Preliminary support for this model has been provided by studies in hereditary diffuse gastric cancer (HDGC) [33] and, more recently, by germline variant burden analyses [38].

Obtaining a genomic map with gene-level resolution required the development of a new method, we called Bayesian Gene Heritability Analysis (BAGHERA), for estimating heritability of low heritability traits at the gene-level. We performed extensive simulations to show that our method provides robust genome-wide and gene-level heritability estimates across different genetic architectures, and outperforms existing methods when used to analyse low heritability traits, such as cancer.

We also recognize the limitations of our work. While our method provides accurate estimates of genome-wide heritability, extremely low heritability diseases could lead to negative gene-level heritability estimates; this was a trade-off to ensure reasonable computational efficiency, although a rigorous model is provided as part of our software. Our analysis does not incorporate functional information, such as gene expression, which limits our power of detecting tissue-specific contributions. On this point, as the genes may be expressed in different cellular compartments, they may contribute to the stromal niches in which cancers develop and their role in tissue specificity of mutations will be of interest to analyse experimentally.

Taken together, our study provides a new view of the genetic architecture of cancer with gene-level resolution. We anticipate that the availability of genome editing techniques will enable testing of the functional mechanisms mediated by cancer heritability genes. We also expect that integrating our results with tumour sequencing data will provide new venues for personalized treatment and patients’ stratification.

## Notes

G.S. and V.F. conceived the study. V.F., G.S. and L.C. designed the model. V.F. wrote the software and performed all analyses, supervised by G.S.. G.S., A.L.H. and F.P. analysed the functional roles of cancer heritability genes. G.S. and V.F. wrote the manuscript with contributions from all authors. The authors also declare no competing interests. The results of our analyses have been deposited in CSV format on Zenodo at: https://doi.org/10.5281/zenodo.3968269.

## Acknowledgments

The authors would like to thank Dr Yongjin Park at the MIT for the useful discussions about GWAS data simulation.

## 4 Methods

### Estimation of heritability at the gene level

Narrow sense heritability, *h*^2^, is defined as the amount of phenotype variance explained by additive genetic effects. Genome-wide association studies (GWAS) provide unique opportunities to study heritability of many diseases; in particular, with the advent of high-density arrays, where more than 500, 000 single nucleotide polymorphisms (SNPs) are genotyped, the heritability explained by these variants, 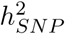, represents a reasonable estimate for *h*^2^.

Our goal is to identify the portion of 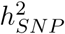 explained by each protein-coding gene, which requires a unique assignment of SNPs to genes to avoid biased estimates.

We denote as genome-wide the amount of heritability explained by all genotyped SNPs, *M*, whereas we refer to the amount of heritability explained by the SNPs in a gene as gene-level heritability. In a model where each SNP has equal contribution to the genome-wide heritability, the per-SNP heritability is simply 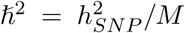. Conversely, if variants can have varying contribution to the genome-wide heritability, we can model the per-SNP heritability as a random variable, 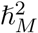, whose expectation is 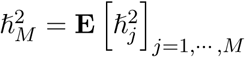, where *M* denotes the number of SNPs used to average the per-SNP contribution to heritability.

We hereby demonstrate that the genome-wide heritability can be expressed as the sum of the gene-level contribution and that the per-SNP genome-wide heritability is the expectation of the per-SNP gene-level heritability. Let *K* be the number of non-overlapping genes in the human genome, each of them with *M*_*k*_ SNPs, the genome-wide heritability can be expressed as 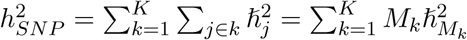 where 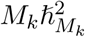 is the amount of heritability explained by all the SNPs in the *k*-th gene. Thus, let the number of SNPs in each gene and the gene-level per-SNP heritability be independent random variables, it is straightforward to prove that the expectation of the gene-level per-SNP heritability is the per-SNP genome-wide estimate 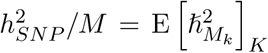. However, estimating 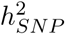 only from SNPs assigned to genes would lead to biased estimates, since the contribution of the SNPs in intergenic regions would be neglected; thus, SNPs outside genic regions are assigned to a nuisance gene, such that the heritability is correctly estimated from all genotyped SNPs.

### A hierarchical Bayesian model for heritability estimation

The estimation of heritability can be modelled as a hierarchical Bayesian regression problem, which provides a robust approach to simultaneously estimate the genome-wide heritability, 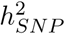, and the gene-level heritability, 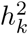, from the observed data *Y*. Our base Bayesian regression model can be defined as follows:

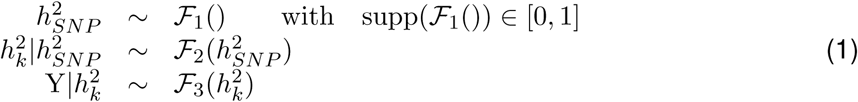

where *ℱ*_1_, *ℱ*_2_, *ℱ*_3_ are suitable distributions.

SNP heritability, 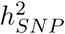, is the ratio of the variance of the additive genetic effects, 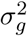, and the phenotypic variance, 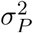. Let 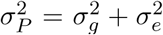, where 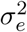 are the non-additive and environmental effects, these quantities can be modelled as random variables with 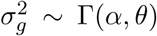 and 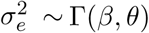, respectively. Since Γ(*α, θ*)*/*(Γ(*α, θ*) + Γ(*β, θ*)) ∼ Beta(*α, β*), a suitable distribution for *F*_1_,in Eq. 1, would be an uninformative Beta distribution, e.g. Beta(1, 1). In practice, the use of a Beta distribution as prior for 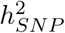 allow us to obtain accurate estimates of heritability in the unit range even for low-heritability diseases, where classical methods are usually unreliable [5].

The gene-level heritability, 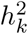, can be modelled as a random variable following a Gamma distribution with shape 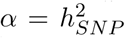 and rate *β* = 1. Therefore, for *ℱ*_2_ = Gamma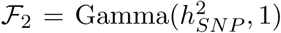, the expectation would be 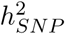, which is an unbiased estimator of the genome-wide heritability.

Finally, our model requires a suitable estimator to regress 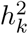 from the observed data. Recently, many methods have been proposed to estimate heritability from GWAS data [59]; however, the vast majority requires genotype data, which are both difficult to obtain, due to privacy concerns, and computationally taxing to analyse, because of high-dimensionality. Thus, we adopted the LD-score (LDsc) regression model [5], which allows estimation of heritability from GWAS summary statistics, such as regression coefficients and standard errors, which are readily available [37].

Thus, for *ℱ*_3_, we rewrote the LDsc model to estimate gene-level heritability, from summary statistics of *M* SNPs in a GWAS with *N* subjects, as follows:

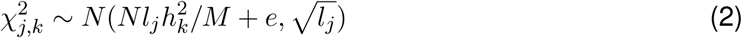

where 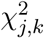 and *l*_*j*_ are the *χ*^2^ statistic and LD score associated with SNP *j* in gene *k*, respectively. The LD score is a quantity defined as 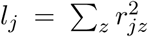, where 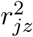 is the linkage disequilibrium between variant *j* and variant *z* in a given population [50]. Moreover, setting the standard deviation to the LD score of the *j*-th SNP allow us to control for heteroskedasticity of the test statistics due to linkage disequilibrium, somehow similar to the weighting scheme used in LDsc. The *e* term accounts for confounding biases and it is modelled using an uninformative normal prior.

### The Bayesian Gene HERitability Analysis (BAGHERA) software

We implemented our hierarchical model (see Eq. 3) as part of the BAGHERA software, which allows simultaneous estimation of genome-wide and gene-level heritability, and predicts heritability genes, that are genes with a per-SNP heritability higher than the genome-wide estimate (see Supplementary Figure 1). Since fitting t he B eta-Gamma m odel i s c omputationally taxing, we relaxed our requirements by modelling 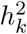 as a random variable following a Normal distribution whose mean is the genome-wide heritability, 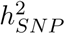, and the standard deviation is controlled by an uninformative Inverse-Gamma prior. While this formulation might provide gene-level heritability estimates outside the unit domain, we found this problem to be well controlled in practice.

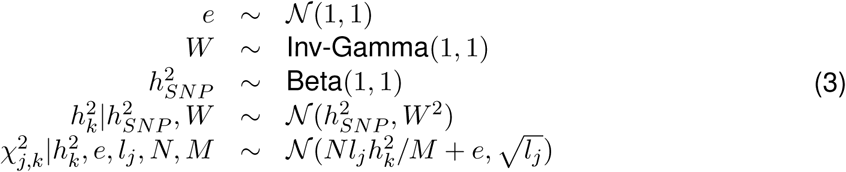

BAGHERA predicts heritability genes by computing the posterior distribution of 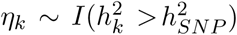, where *I* is a function that returns 1 if the evaluated condition is true, and 0 otherwise. The expectation of the posterior distribution of *η*_*k*_, **E**[*η*_*k*_], is the probability of the heritability of gene *k* of being higher than the genome-wide estimate; specifically, we report as heritability genes, those with **E**[*η*_*k*_] ≥ 0.99. For each gene, we also report effect sizes in terms of fold-change with respect to the genome-wide heritability estimate, 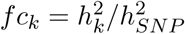.

We use the No-U-Turn Sampler as implemented in PyMC 3.4 [40], using 4 chains with 10^4^ sweeps each and a burnin step consisting of 2, 000 samples. Convergence of the sampling process was assessed based on the Gelman-Rubin convergence criterion.

BAGHERA is released as a Python software package under MIT license, and it is available on GitHub (https://github.com/stracquadaniolab/baghera), as installable package on Anaconda, and as a Docker image. BAGHERA also implements the Beta-Gamma model described in the previous section, called BAGHERA-Γ. Alongside the source code, we also provide a Snake-make workflow (https://github.com/stracquadaniolab/workflow-baghera).

### UK BioBank summary statistics processing and curation

We used summary statistics of the UK BioBank GWAS for ICD10 classified cancer types [35]. We developed a custom pipeline to assign LD scores to SNPs, and SNPs to human genes (Supplementary Material). For each dataset, we mapped each SNP to a precomputed LD score. We used pre-computed LD scores for SNPs on autosomal chromosomes with minor allele frequency MAF > 0.01 in the European population (EUR) of the 1000 Genomes project. We removed the SNPs on chr6:26,000,000-34,000,000, since this region contains the Major Histocompatibility Complex (MHC) that have unusual genetic patterns and is known to affect GWAS result interpretation [5, 23]. Overall, our analysis is conducted on 1285620 SNPs over 22 chromosomes.

We then used Gencode v31 to determine the genomic coordinates of protein coding genes in the GRCh37 human genome. We then merged overlapping genes by creating a new pseudogene, whose name reports the merged gene names and whose boundaries are defined as the first and last base-pair of the overlapping genes. We assigned a SNP to a gene if it is within ± 50kb from the gene boundaries, which allow us to account for cis-regulatory elements, overall 55% of SNPs were mapped to a gene. For BAGHERA we considered only those genes harboring at least 10 variants. Our dataset consists of 15025 genes, 12042 of them are harboring more than 10 SNPs, then they are used for the UKBB analysis.

#### 4.1 Enrichment analyses

We used a one-tailed Fisher’s exact test for all enrichment analyses, with p-value adjusted using the Benjamini-Hochberg procedure. Since genes in our analysis might represent overlapping protein-coding regions, we post-processed our gene lists by converting each composed region into the set of its genes for functional characterization and annotation. The overlap with cancer datasets has instead been tested with individual one-tailed Fisher’s exact tests.

