## Supplementary Materials for "Gene-level heritability analysis explains the polygenic architecture of cancer"

August 3, 2020

**5 Supplementary Tables** **48**

[*]Institute of Quantitative Biology, Biochemistry, and Biotechnology, SynthSys, School of Biological Sciences, University of Edinburgh, Edinburgh, EH9 3BF, UK

[†]School of Computer Science and Electronic Engineering, University of Essex, Colchester CO4 3SQ, United Kingdom

[‡]Molecular Oncology Laboratories, Department of Oncology, The Weatherall Institute of Molecular Medicine, University of Oxford, Oxford, UK.

[§]Nuffield Department of Clinical Laboratory Sciences, University of Oxford, John Radcliffe Hospital, Oxford, UK

[¶]Institute of Quantitative Biology, Biochemistry, and Biotechnology, SynthSys, School of Biological Sciences, University of Edinburgh, Edinburgh, EH9 3BF, UK. Phone: +44 (0) 131 6507193,. Corresponding author.

### 1 Simulated datasets

We performed extensive simulations to assess the performance of our hierarchical Bayesian model, as implemented in BAGHERA.

First, we generated datasets with a realistic genetic architecture and linkage disequilibrium patterns by using the data from the 1000 Genomes Project, see section 1.1. Since these simulations are computationally taxing and existing tools do not scale for genome-wide simulations, we restricted our analyses to SNPs located on chromosome 1. First, we used these datasets to test the performance of BAGHERA for regressing genome-wide heritability estimates. We then tested the performance for gene-level heritability testing.

Nonetheless, we also wanted to explore the behaviour of BAGHERA in presence of whole genome datasets, not only one chromosome, which is the common usage of the method. Henceforth, we decided to simulate whole genome summary statistics with a varying number of heritability genes and enrichments, see section 1.2.

When assessing the performance of BAGHERA in detecting heritability genesets, we remind that our model estimates the posterior distribution of $\eta_k$, whose expected value is the probability of the per-SNP gene-level heritability to be higher than the per-SNP genome-wide estimate. Thus, we can find how many heritability genes are recovered as a function of the expected value for $\eta_k$. Since heritability genes are known a-priori in our experiments, we derived Receiver Operating Characteristic (ROC) curves and computed the corresponding Area Under the Curve (AUC) for the various scenarios. While ROC curves allow straightforward comparison of different experimental conditions, they can be problematic for interpreting genomic data, since the number of positives samples is significantly smaller than the negatives. For this reason, we also derived Precision and Recall (PR) curves as a more accurate approach to control Type 1 errors.

Below we are going to describe in detail the generation of the datasets and the main results of the simulation analysis.

#### 1.1 Simulated datasets with realistic architecture

With HAPGEN2 [4] we simulated N = $50,000$ subjects and M = $100,000$ SNPs on chromosome 1, from 1000 Genome reference data from $503$ European ancestry subjects; here, we used haplotype data downloaded from the IMPUTE website (`https://mathgen.stats.ox.ac.uk/impute/impute_v2.html#download`). Finally, we filtered out SNPs with minor allele frequency (MAF) larger than 0.01, leading to a final dataset consisting of 99,586 SNPs.

We then controlled whether the simulated genetic architecture was coherent with the one observed in Europeans. To do that, we estimated the correlation between the observed minor allele frequency (MAF) between the 1000 Genomes dataset and our simulation; our analysis found a statistically significant correlation (Pearson correlation coefficient $\rho = 0.9929$, $P \leq 10^{-5}$), suggesting that our strategy was appropriate to generate realistic genotype data.

Summary statistics were then simulated following a dense and gene-level effect size model. We used the dense effect model to test the robustness of the genome-wide heritability estimates returned by BAGHERA. To do that, we explicitly set the variance of the SNPs to be $\hbar^2 = h^2/M$, with $h^2 = [0.01, 0.1, 0.2, 0.5]$; for each parameters' setting, we generated $5$ different datasets (Figure 1A in the main manuscript).

We then assessed BAGHERA as a method for discovering heritability genes. To do that, we considered causal only those SNPs that are in a predefined set of genes Hence, we tested whether BAGHERA was able to identify them under different genome-wide heritability levels and number of heritability genes. Out of all genes $G$ , we select a portion of them, $sg$, as significant, $G_{sig} = G \times sg$. We then assign $90\%$ of the variance to the $M_{sig}$ SNPs falling into the

$G_{sig}$ genes. The remaining $10\%$ variance is equally distributed to the others. We simulated data with $h^2 = \{0.01, 0.05, 0.1, 0.2\}$ and $sg = 0.01$; taken together, we obtained $G_{sig} = 13$ heritability genes out of $1322$ genes with more than $10$ SNPs on chromosome 1. For each combination of parameters, we simulated $5$ datasets. Results are in Figure 1B and 1C in the main manuscript.

##### 1.2 Whole genome simulated datasets

Restricting the analysis to just the chromosome 1 would not provide conclusive evidence on the good performances of our method, which was designed to run on high-density genotype data. We then used a simpler model, which does not require genotype data, to generate simulated summary statistics for 22 chromosomes with a varying number of heritability genes and heritability enrichment.

We assigned random effect sizes to SNPs with MAF $> 0.01$ in the European populations of the 1000 Genomes Phase 3 project by sampling from a normal distribution and weighting the random variate by $w_j = \sqrt{(1 + \frac{N}{M} h_k^2 l_j)}$, where $h_k^2$ is the gene-level heritability and $l_j$ is the LD score of the $j$-th SNP in the dataset [1]. Using LD scores allow us to account for positional constraints and LD patterns without using genotype data. We then randomly selected a fraction of genes as heritability genes and set their heritability $h_k^2 = fc_k \times h_{SNP}^2$ where $h_{SNP}^2$ is the genome-wide heritability, $fc_k$ is the fold-change in heritability in gene $k$ compared to the genome-wide estimate.

In our experiments, we set the genome-wide heritability to $h_{SNP}^2 = [0.01, 0.1, 0.2]$, to mimic a disease with a reasonably low heritability, such as cancer. We then considered $p = 1\%$ of the genes in the genome as the heritability genes, and set the heritability fold-change as $fc_k = [1.1, 5, 10, 30]$. For each possible parameter setting, we generated $3$ independent datasets, which resulted in a testbed consisting of $36$ datasets in total. We are expecting BAGHERA to be unable to identify any difference between the set heritability genes and the rest. Fold change value $fc = 1.1$ is used as control. Indeed, in all results we find that the for the control simulations the ROC and PR curves report expected random performances with AUCs respectively $AUC_{ROC} \sim 0.5$ and $AUC_{PR} \sim p$, where $p$ is the proportion of heritability genes.

While the AUC for both PR and ROC curves are in the main text (Figure 1C) we show the ROC and PR curves in Supplementary Figures 2 and 3.

#### 2 Comparison with other methods.

##### 2.1 Comparison of genome-wide heritability estimates between BAGHERA and LDsc

We compared BAGHERA genome-wide estimates with the observed $h_{SNP}^2$ estimates of LD score regression (LDsc) [2]. In Supplementary FIgure 4, it is straightforward to note that BAGHERA is more robust on low heritability malignancies, including $9$ cases where LDsc reported negative estimates.

##### 2.2 Comparison between BAGHERA and HESS

We compared the estimates of local heritability with HESS [3], which, to date, is the only method for the estimation of local heritability that can be readily applied on regions smaller than a chromosome.

First, we identify the main differences between the two methods, which could confuse the interpretation of the results. HESS has been shown to provide robust heritability estimates for genomic regions defined as LD independent. BAGHERA, instead, provides heritability estimates for any non overlapping set of genomic regions, including $\approx 15,000$ protein-coding genes in the human genome. Thus, BAGHERA can provide heritability estimates at a much higher genomic resolution.

It is also important to note the different output returned by BAGHERA and HESS. We remind the reader that each region explains a portion of heritability $h^2_{(k)} = \sum_{j=1}^{M_k} \hbar^2_j$, where $h^2_{(k)}$ is the output of HESS. With the notation we introduced in the methods, we can notice that $h^2_{(k)}/M_k = h^2_k/M$, where $h^2_k$ is the gene-level heritability estimated by BAGHERA. Both methods, however, test whether the local single SNP heritability, either $h^2_k/M$ or $h_{(k)}/M_k$ is larger than expected genomewide $\hbar^2_M$.

It is also worth mentioning that the two methods employ very different testing strategies; after the estimation of local heritability, HESS converts the estimates to z-scores to obtain a p-value for each region, and then uses Bonferroni correction to control the family-wise error rate. BAGHERA instead uses a Bayesian hierarchical model to return the posterior distribution of the genome-wide and gene-level heritability, along with the one of an indicator function ($\eta$), which can be used to select regions with heritability enrichment.

###### 2.2.1 Comparison of local estimates for HESS and BAGHERA

We then applied both HESS and BAGHERA on the two datasets from the UKBB with the strongest signal: C50 breast , C61 prostate. In order to compare the local heritability estimates of the two methods, we used the same set of SNPs and the $1703$ regions originally used in HESS, although we filtered out $10$ of them having less than $10$ SNPs.

Here are the overall results (H for HESS and B for BAGHERA), listing the genome-wide estimates $h^2$, the number of significant genomic regions (sig) and the correlation between the two local heritability estimates (Pearson's $\rho$ and pvalues $p$).

| Dataset | $\mathbf{h^2(se)}$ H | $\mathbf{h^2(sd)}$ B | sig H | sig B (common) | $\rho$ | p |
|---|---|---|---|---|---|---|
| C50 | $0.0111(0.00316)$ | $0.0149(0.0018)$ | 2 | 119(2) | $0.78$ | $\leq 1e^6$ |
| C61 | $0.00896(0.00316)$ | $0.0098(0.0017)$ | 1 | 116(1) | $0.76$ | $\leq 1e^6$ |

In Supplementary Figure 5 and 6, we show the results of the analysis; for each panel, the first figure shows HESS and BAGHERA values of $h^2_{(k)}$, the second one is limited to the significant regions defined by BAGHERA and overlaps HESS estimates. The last figure instead, rescales HESS $h^2_{(k)}$ estimates to BAGHERA's $h^2_k$, as $h^2_{(k)}/M_k \times SNPs$.

It is straightforward to note that BAGHERA provides more robust local heritability estimates, since the number of negative estimates is significantly lower than HESS, as more clearly shown when rescaling the results. While BAGHERA is not immune from negative local heritability estimates, in practice this phenomenon seems to be controlled better.

### 3 Analysis of 38 UKBB Cancer datasets

#### 3.1 UKBB analysis Data

From `http://www.nealelab.is/uk-biobank` we downloaded on $30/07/2019$ the descriptive tables. From the list of all phenotypes we filter out those of interest using only the ICD10 terms, corresponding to malignant neoplasms. Full explanation of the ICD10 terms in the UKBB can be found here `http://biobank.ndph.ox.ac.uk/showcase/field.cgi?id=41202`. Malignancies have CXX codes (C00-C97). We removed the benign neoplasms and in situ carcinoma/melanoma and the secondary neoplasms (C77,C78,C79) The final table has 38 terms. Mapping the SNPs to the genes we obtained a wide spectrum of genes with different sizes, see Supplementary Figure 7

**LD-score** data was downloaded from https://data.broadinstitute.org/alkesgroup/LDSCORE/ on 15/03/2018-15/04/2018. We use **Gencode** version 31 available at https://www.gencodegenes.org/). The **Gene Ontology (GO) slim** dataset was generated using the map2slim utility of the OWL tools on 16/10/2019. We also report enrichment results for the entire Gene Ontology dataset (downloaded from MSigDB. http://software.broadinstitute.org/gsea/msigdb).The **Precision Oncology Knowledge Base (OncoKB)** dataset, alongside the MSK and Vogelstein data were downloaded on 01/10/2018. The **Cancer Gene Census** data was downloaded from https://cancer.sanger.ac.uk/census on 17/07/2019. PCAGW data, **the compendium of mutational driver elements** was downloaded on 24/04/2020 from https://dcc.icgc.org/pcawg/. All dates are formatted as dd/mm/yyy.

#### 3.2 Relationship between GWAS raw results and BAGHERA

Heritability analysis from summary statistics is built on top of GWAS results. We then tested whether higher heritability could be explained by the presence of genome-wide significant SNPs ($P < 5 \times 10^{-8}$) nearby protein-coding regions.

For each cancer, we identified genes harbouring at least $1$ genome-wide significant SNP, and denoted this set as minSNPs. We found $119$ minSNPs in total, with at least $1$ minSNP in $18$ of the $38$ cancers (Supplementary Table 4). This is a striking difference compared to the $1523$ heritability genes found in total for all $38$ malignancies; interestingly, our method was able to recover $98$ $(82\%)$ of the minSNSP suggesting that it can detect heritability genes regardless of the association strength of their SNPs.

We then proceeded to analyse whether there is a correlation between minSNP p-values and heritability estimates. Interestingly, while many minSNPs are also heritability genes, we do not observed a linear relationship between the $\eta$ and p-values, see Supplementary Figure 50 and 51. However, as expected, there is a correlation between each gene average statistics and local heritability, Supplementary Figure 50.

#### 3.3 Relationship with self-reported tumours

In this paper we are analysing the GWAS results of histologically characterised tumours. However, the UK Biobank also provides a self-reported classification of the patients on which GWAS have been conducted. Here we report the results for the round 1 of GWAS analysis from the Neale lab for self-reported cancer types. Indeed, in the second release of the data, which is the one we used in the main paper, cancer types such as breast and testicular cancer were analysed with an inconsistent number of controls.

In supplementary table 5, we report an overview of our results. In particular, we found only 11 datasets with $\hat{\chi}^2 > 1.01$ compared to the 17 of the ICD10 classification. Prevalence is also higher for the ICD10 classification datasets $(0.0029)$ than the self-reported on average $(0.0023)$. Overall, we find that directly comparing the two datasets might be difficult. In Supplementary Figure 54, we show the Jaccard similarity for each pair of datasets, when we consider the significant genes. For this data, we used the Gencode v27 annotation, which might have resulted in a slightly different mapping of the genes. As expected, in some cases there is a good similarity between the same cancers, however the great differences in signal and the different mapping might be affecting the discovery power, especially for tumours with fewer heritability genes.

We then proceeded with the analysis of the self-reported dataset, similarly to what shown for the histologically characterised tumours. Unsurprisingly, breast and prostate cancer still show high values of heritability, and both breast and testicular cancer have more than $30\%$ of their heritability explained by HG Supplementary Figure 55A . As expected, these datasets, whose signal is lower compared to the ICD10 classified malignancies, have a higher heritability enrichment, consistent with results on simulated data (Supplementary Figure 55B). The percentage of CHGs occurring in multiple malignancies is almost identical to the one for the 38 histologically characterised tumours (Supplementary Figure 55C) and with similar pan-cancer CHGs (Supplementary Figure 55D and Figure 52D).

Interestingly, when characterising the CHGs, we find the overall results to be highly consistent with those of the histologically characterised dataset (see Supplementary Figure 56). We would like to highlight that $90\%$ of GO terms significant in this analysis are shared with the results for the 38 cancers above and, also in this case, there is also a significant enrichment for tumour suppressors over oncogenes.

### 4 Figures

#### 4.1 BAGHERA outline

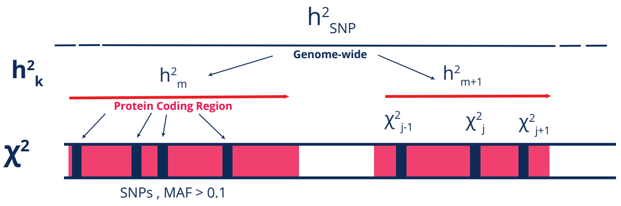

$$
\begin{aligned}
e &\sim \mathcal{N}(1,1) \\
W &\sim \text{Inv-Gamma}(1,1) \\
h^2_{SNP} &\sim \text{Beta}(1,1) \\
h^2_k | h^2_{SNP}, W &\sim \mathcal{N}(h^2_{SNP}, W^2) \\
\chi^2_{j,k} | h^2_k, e, l_j, N, M &\sim \mathcal{N}(N l_j h^2_k / M + e, \sqrt{l_j})
\end{aligned}
$$

Supplementary Figure 1: **Graphical representation of BAGHERA's model.** On the left, from bottom to top we represent how single SNP's $\chi^2$ are grouped into a single gene-level heritability term and then into the genome-wide $h^2$ estimate. On the right we report the actual Bayesian model with the Normal prior on $h^2_k$.

#### 4.2 Performance on simulated data

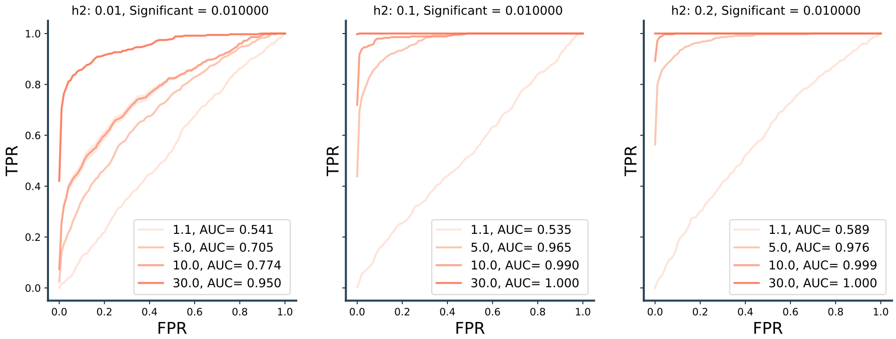

Supplementary Figure 2: **ROC Curves for summary statistics simulations.** Receiver Operating characteristic curve for the data simulated from summary statistics. Different colours mark different fold changes (1,1, 5,10,30) while each column corresponds to different values of $h^2$.

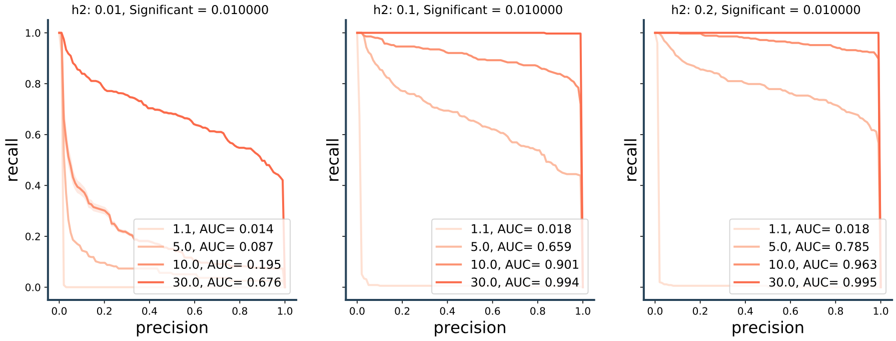

Supplementary Figure 3: **PR Curves for summary statistics simulations.** Precision Recall curves for the data simulated from summary statistics. Different colours mark different fold changes (1,1, 5,10,30) while each column corresponds to different values of $h^2$.

#### 4.3 Comparison with other methods

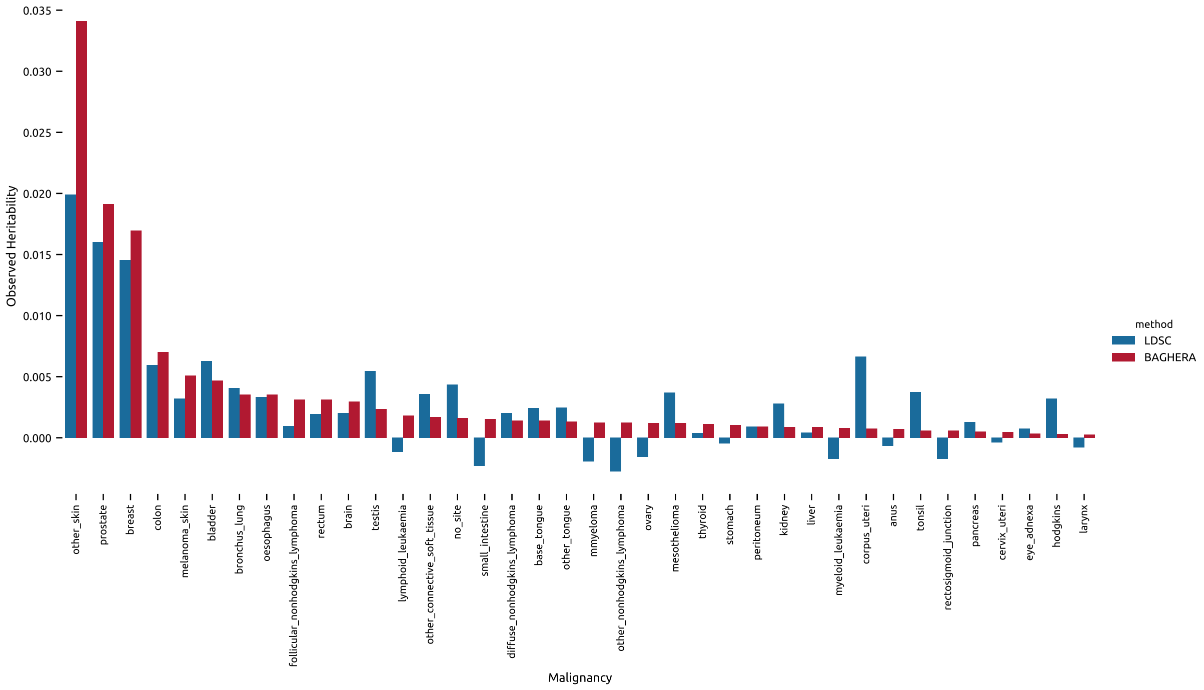

Supplementary Figure 4: **Comparison between LDSC and BAGHERA.** For each of the 38 malignancies (x-axis) we show the observed $h^2$ estimate (y-axis) for LDSC and BAGHERA.

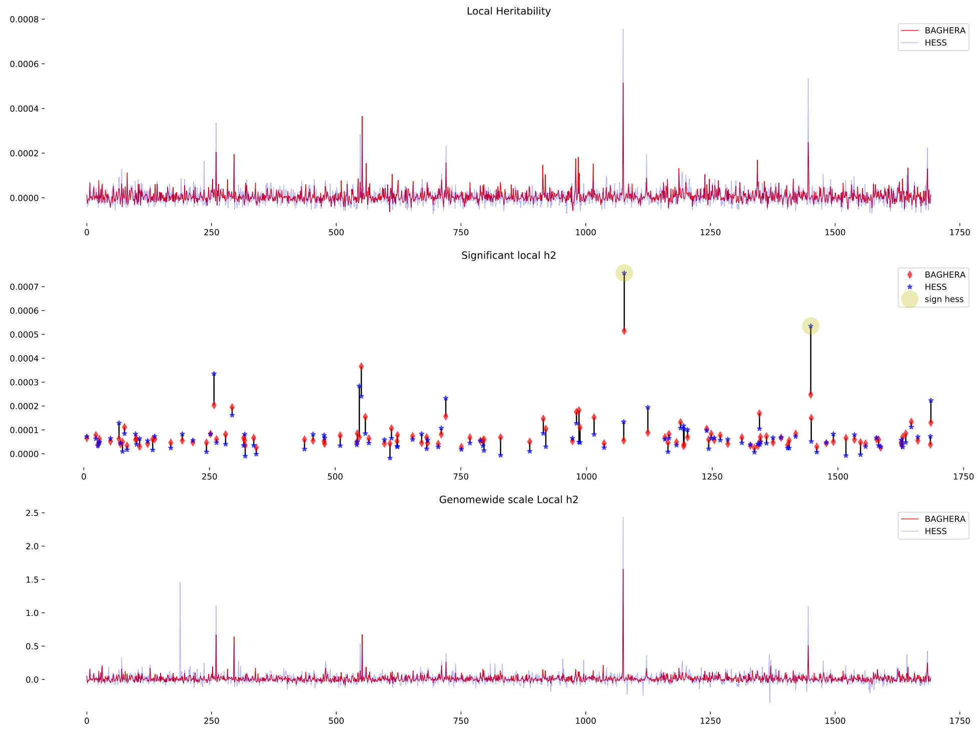

Supplementary Figure 5: **Comparison C50, breast cancer** For each panel, the first figure shows HESS and BAGHERA values of local heritability $h^2_{(k)}$. The second one reports the values of $h^2_{(k)}$, but it is limited the regions that are deemed as significant by BAGHERA and HESS. The last figure instead, rescales HESS estimates to those native of BAGHERA as $h^2_{(k)}/M_k \times SNPs$.

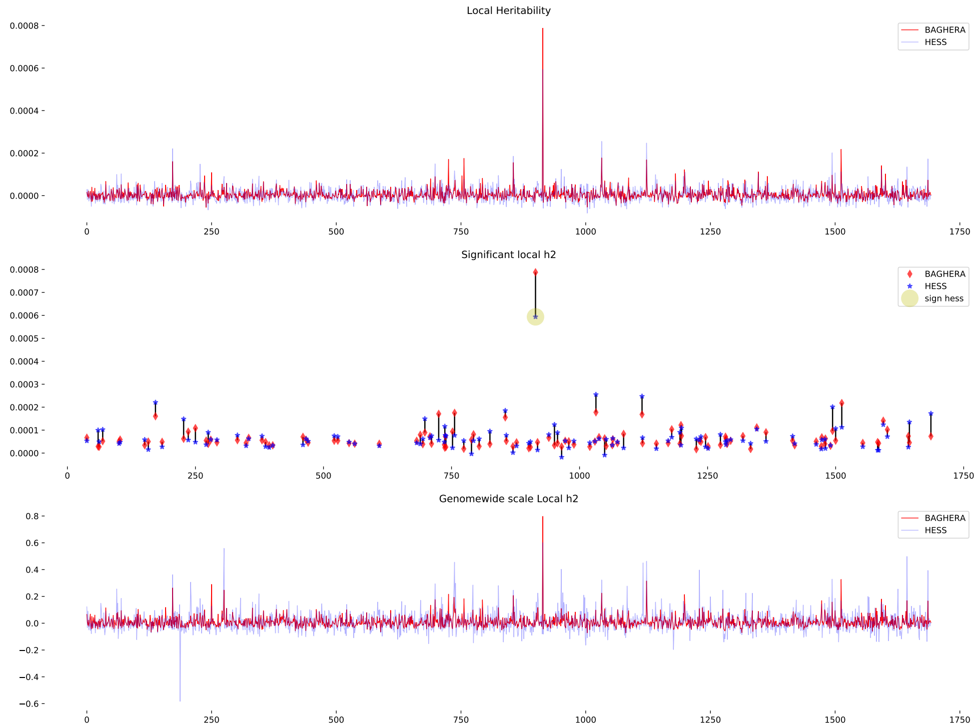

Supplementary Figure 6: **Comparison C50, prostate**. For each panel, the first figure shows HESS and BAGHERA values of local heritability $h^2_{(k)}$. The second one reports the values of $h^2_{(k)}$, but it is limited the regions that are deemed as significant by BAGHERA and HESS. The last figure instead, rescales HESS estimates to those native of BAGHERA as $h^2_{(k)}/M_k \times SNPs$.

#### 4.4 GWAS results for the UKBB

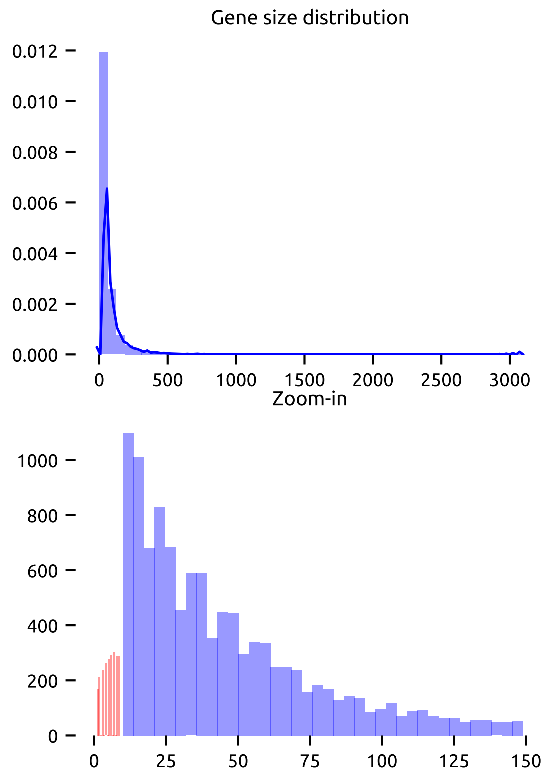

Supplementary Figure 7: **SNPs in genes.** Here we show the distribution (density on y-axis) of the number of SNPs in the UKBB datasets for each gene used by BAGHERA (x-axis). The second figure is a zoomed in version of the first one and it reports the actual frequency (y-axis) of genes harboring between 0 and 150 SNPs of the UKBB. We denote in red the genes that are filtered out, whose SNPs are added to the nuisance gene

##### 4.5 QQ-plots

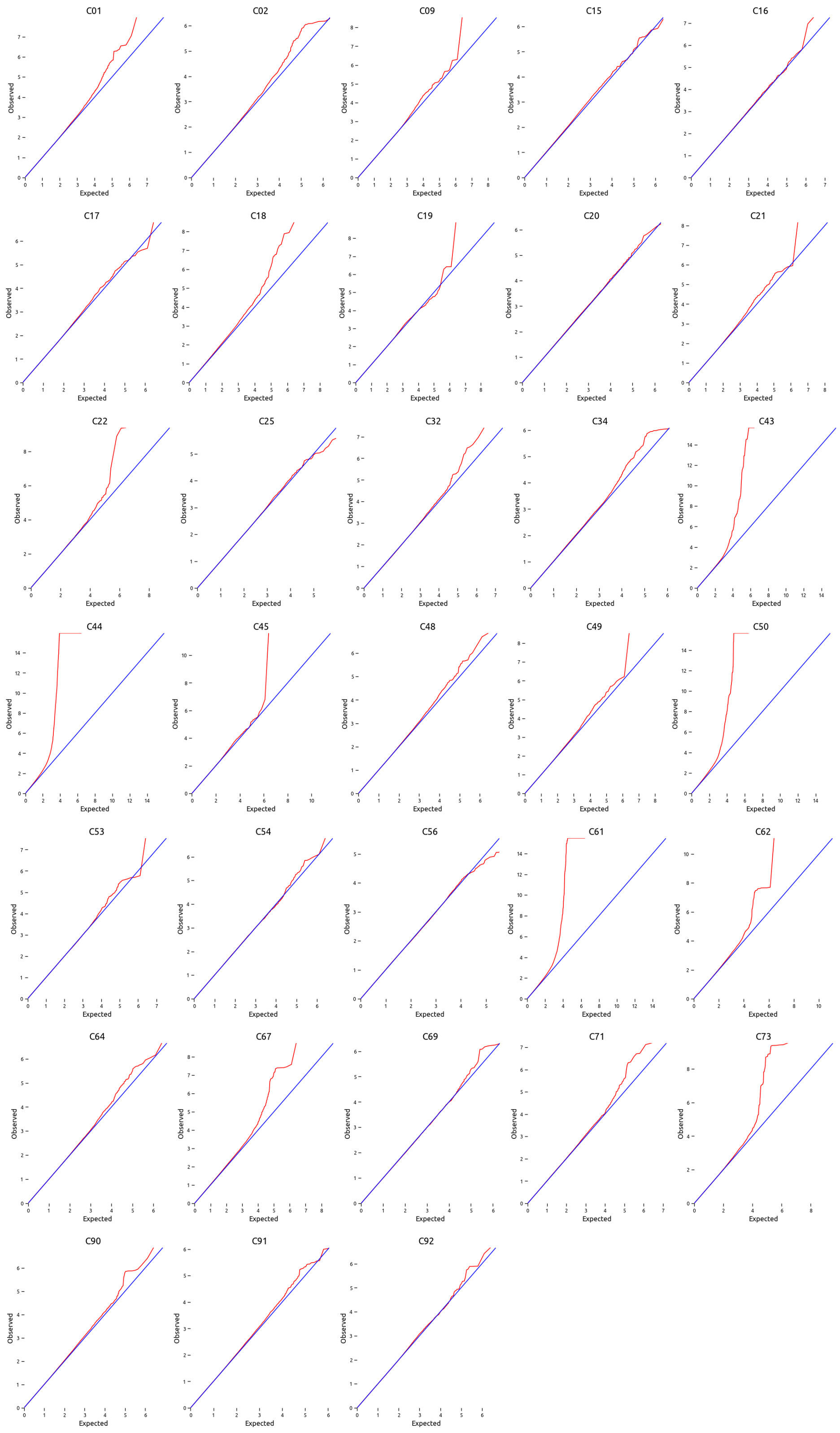

Supplementary Figure 8: **QQ-plots of all datasets.** QQ plots of the GWAS results for all UK Bio Bank datasets. These are the summary statistics on which we run BAGHERA. FOr each SNP we report the expected (x-axis) and observed (y-axis). Titles report the ICD10 code of the dataset. We can that for C44 (Other malignant neoplasm of skin) there is a considerable inflation in the statistics, which reflects in a high average $\chi^2$. Conversely, C56, Malignant Neoplasm of ovary, and C25, Malignant neoplasm of pancreas show smaller statistics than expected.

#### 4.6 BAGHERA UKBB results

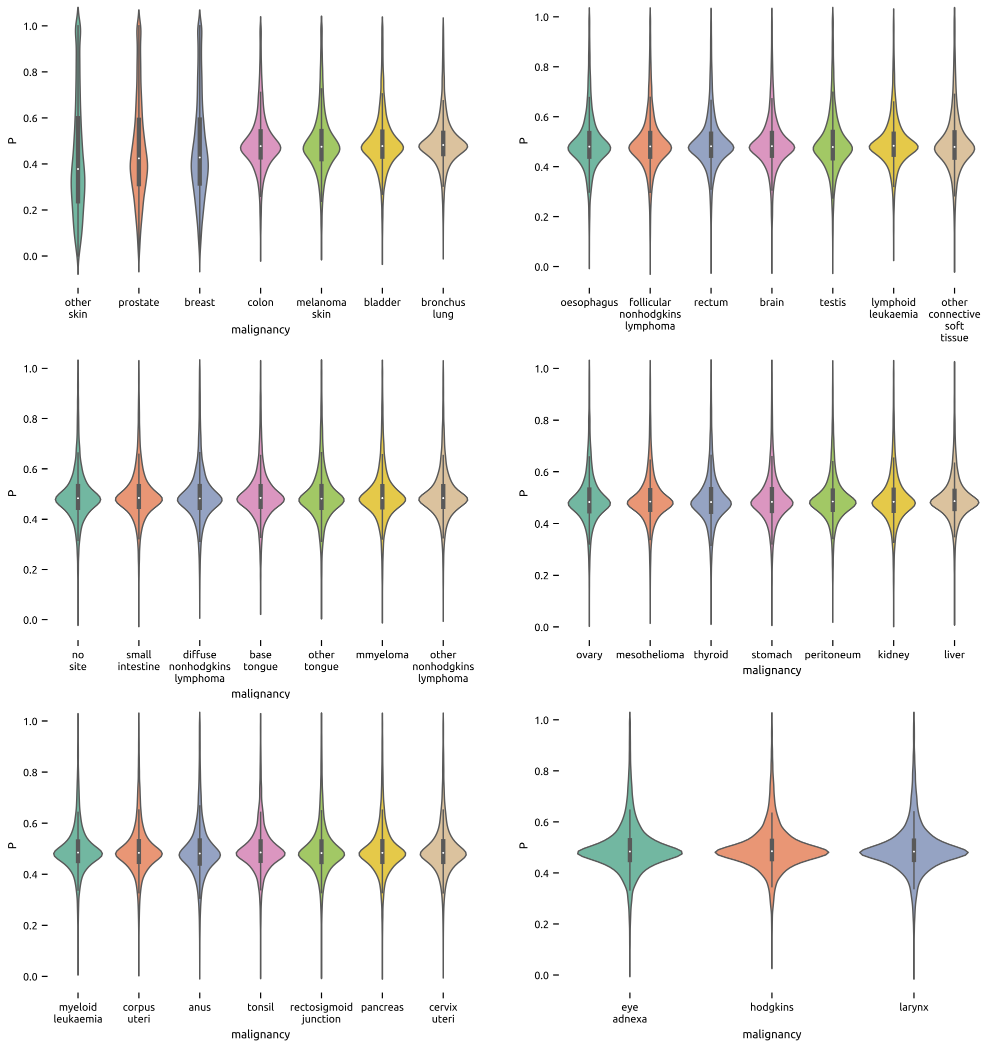

Supplementary Figure 9: $\eta$ **distributions.** For each analysed dataset (x-axes) we report the distribution of the indicator function $\eta$, in the implementation this term is named P (y-axes). Each violin-plot shows the mass distribution of the $\eta$ values.

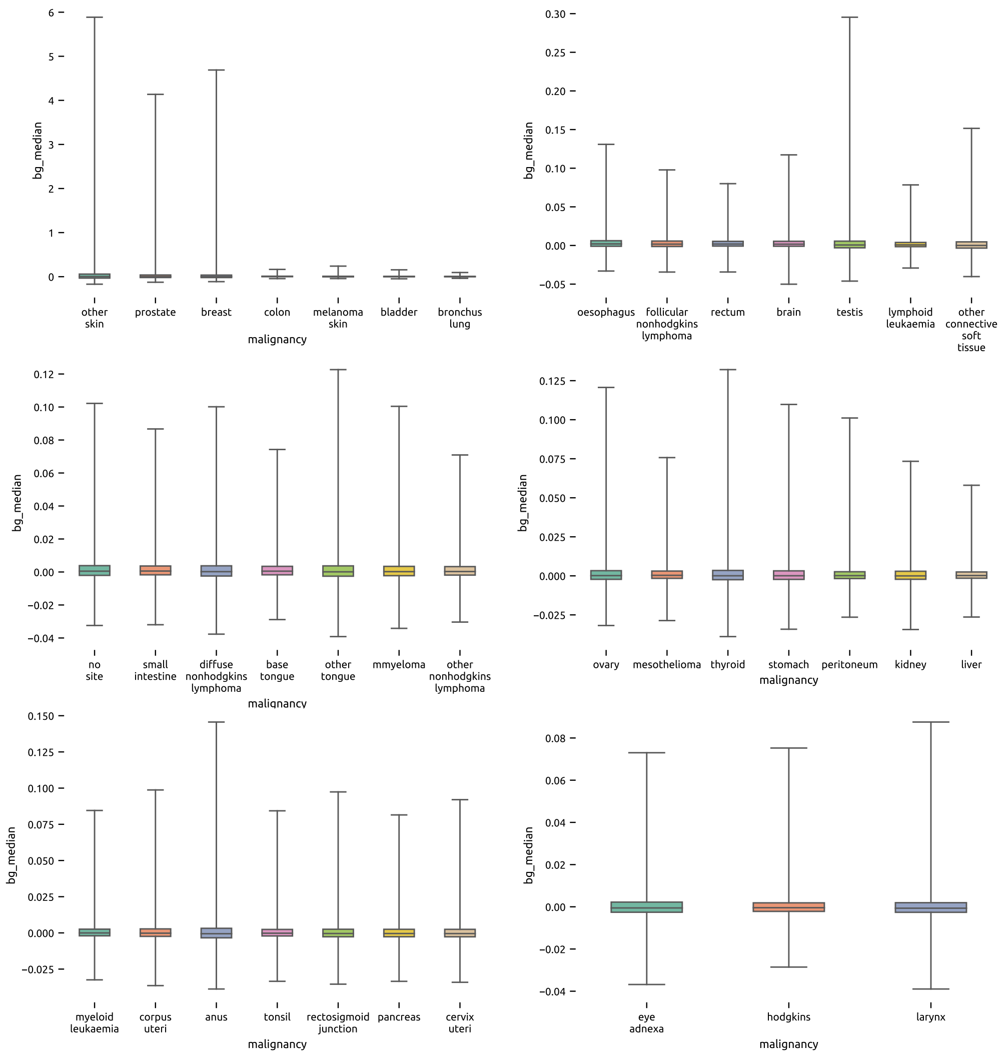

Supplementary Figure 10: **Local heritability distributions** For each analysed dataset (x-axes) we show the boxplot of the median $h_k^2$ for each gene, in the implementation this term is named bg median (y-axes).

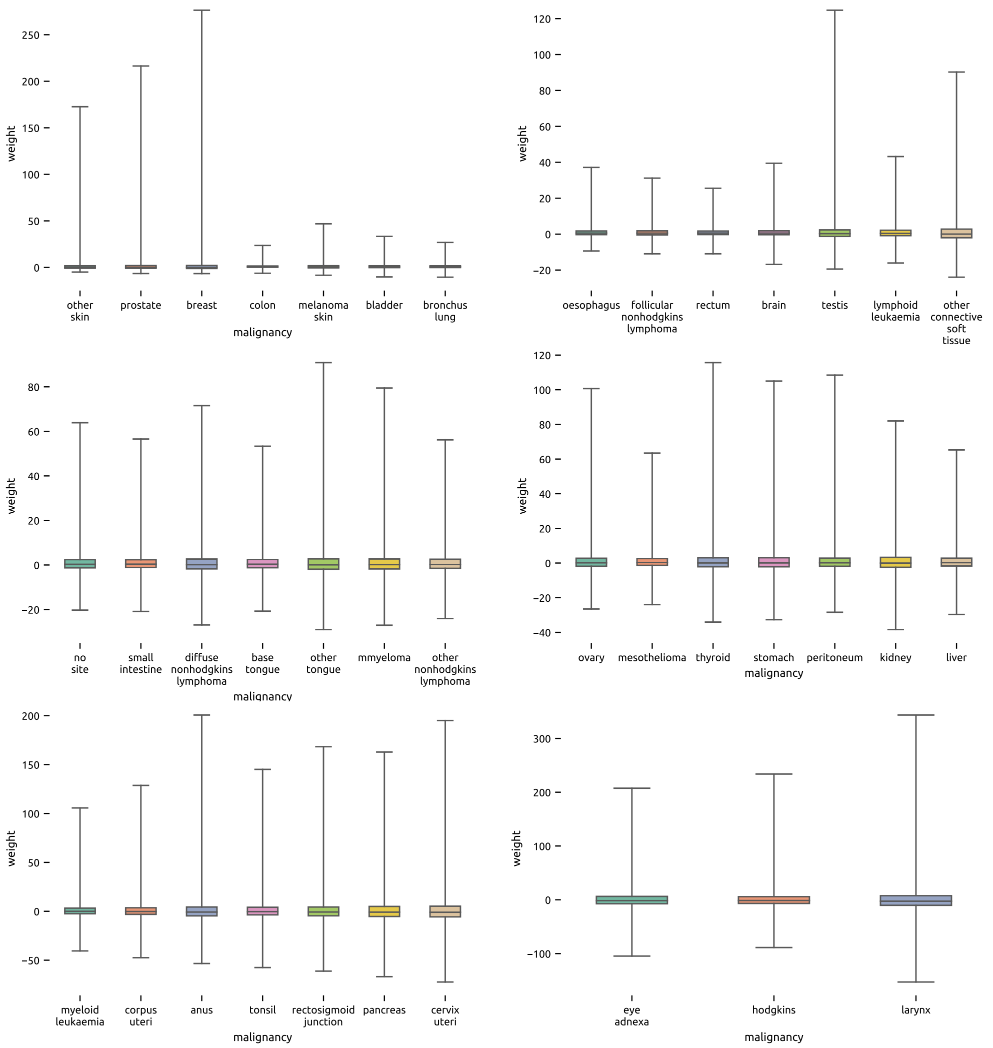

Supplementary Figure 11: **Local heritability Weights.** For each analysed dataset (x-axes) we show the boxplot of the local heritability weights $w_k = (h_k^2 - h^2)/h^2$ for each gene. Please note that what we denote as fold change has the following relationship with the weights $fc = w_k + 1$.

##### 4.6.1 Single cancer results

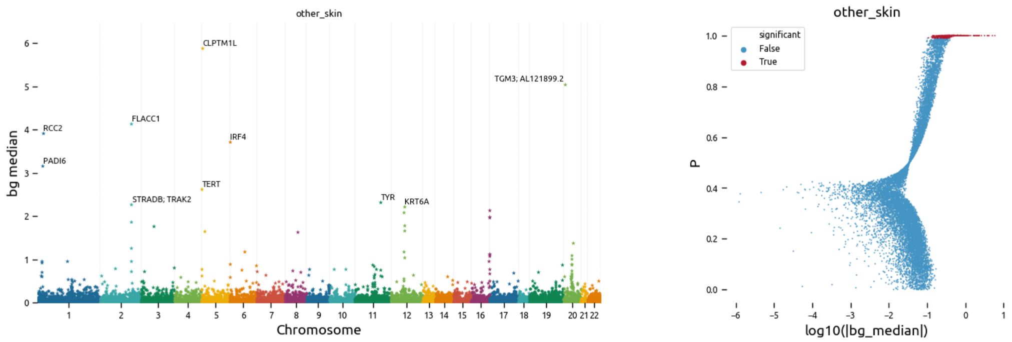

Supplementary Figure 12: **BAGHERA results for C44: Other Malignant neoplasms of skin.** On the left we show a Manhattan plot of the local heritability $h_k^2$ (y-axis bg median in the implementation) for each gene in the analsis (x-axis, genes are sorted by position). Star markers denote significant genes, we report the names of the top scoring genes. On the right, we show the relationship between the local heritability estimates and the testing results. For each gene, on the x-axis we report the $log_{10}|h_k^2|$ and on the y-axis we report the indicator function value $\eta$, respectively bg median and P in the implementation. Significant genes are represented in red. The discontinuity at P$\sim 0.5$ denotes the change between positive and negative values of local heritability.

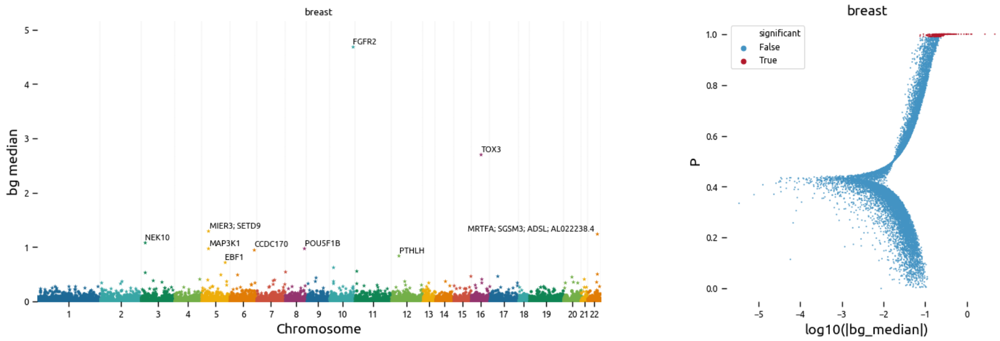

Supplementary Figure 13: **BAGHERA results for C50: Malignant neoplasms of breast**. On the left we show a Manhattan plot of the local heritability $h_k^2$ (y-axis bg median in the implementation) for each gene in the analsis (x-axis, genes are sorted by position). Star markers denote significant genes, we report the names of the top scoring genes. On the right, we show the relationship between the local heritability estimates and the testing results. For each gene, on the x-axis we report the $log_{10}|h_k^2|$ and on the y-axis we report the indicator function value $\eta$, respectively bg median and P in the implementation. Significant genes are represented in red. The discontinuity at P$\sim 0.5$ denotes the change between positive and negative values of local heritability.

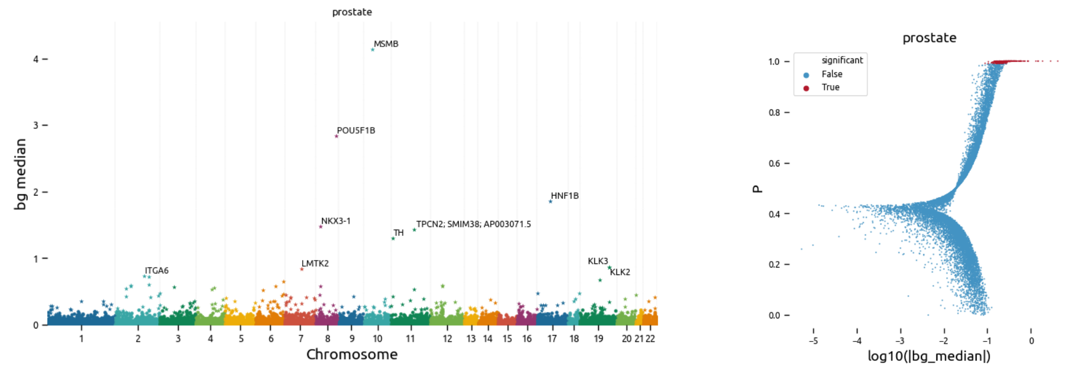

Supplementary Figure 14: **BAGHERA results for C61: Malignant neoplasms of prostate**. On the left we show a Manhattan plot of the local heritability $h_k^2$ (y-axis bg median in the implementation) for each gene in the analsis (x-axis, genes are sorted by position). Star markers denote significant genes, we report the names of the top scoring genes. On the right, we show the relationship between the local heritability estimates and the testing results. For each gene, on the x-axis we report the $log_{10}|h_k^2|$ and on the y-axis we report the indicator function value $\eta$, respectively bg median and P in the implementation. Significant genes are represented in red. The discontinuity at P$\sim 0.5$ denotes the change between positive and negative values of local heritability.

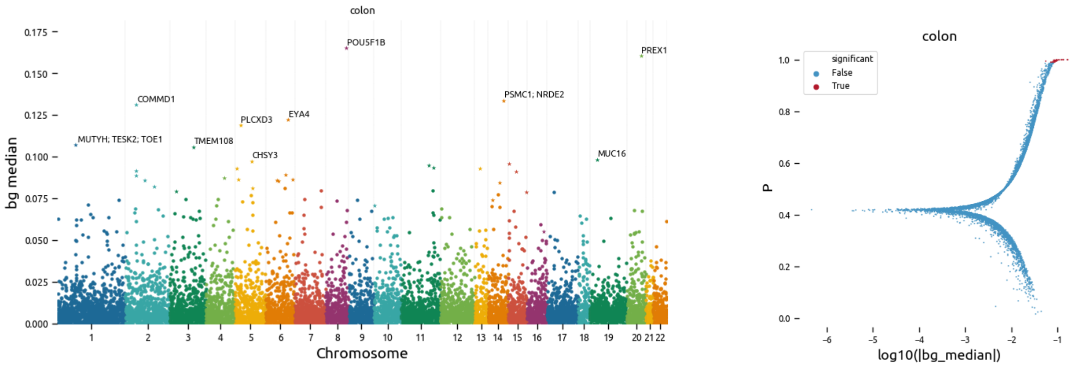

Supplementary Figure 15: **BAGHERA results for C18: Malignant neoplasms of colon**. On the left we show a Manhattan plot of the local heritability $h_k^2$ (y-axis bg median in the implementation) for each gene in the analsis (x-axis, genes are sorted by position). Star markers denote significant genes, we report the names of the top scoring genes. On the right, we show the relationship between the local heritability estimates and the testing results. For each gene, on the x-axis we report the $log_{10}|h_k^2|$ and on the y-axis we report the indicator function value $\eta$, respectively bg median and P in the implementation. Significant genes are represented in red. The discontinuity at P$\sim 0.5$ denotes the change between positive and negative values of local heritability.

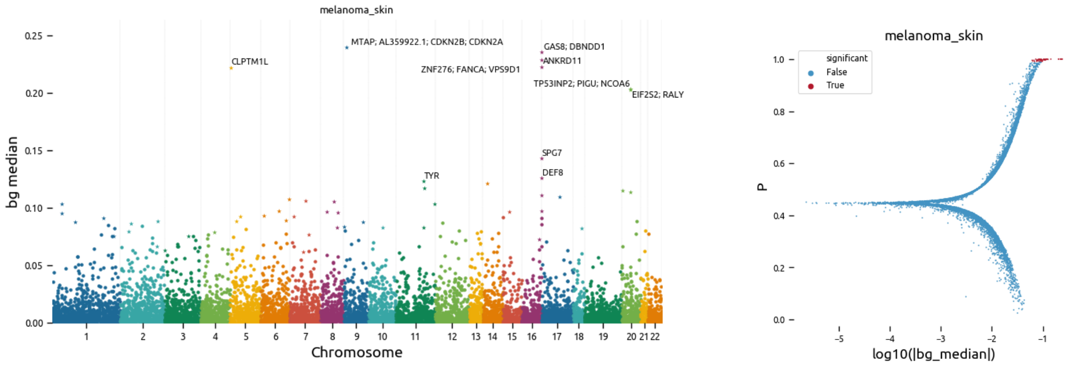

Supplementary Figure 16: **BAGHERA results for C43: Malignant melanoma of skin**. On the left we show a Manhattan plot of the local heritability $h_k^2$ (y-axis bg median in the implementation) for each gene in the analisis (x-axis, genes are sorted by position). Star markers denote significant genes, we report the names of the top scoring genes. On the right, we show the relationship between the local heritability estimates and the testing results. For each gene, on the x-axis we report the $log_{10}|h_k^2|$ and on the y-axis we report the indicator function value $\eta$, respectively bg median and P in the implementation. Significant genes are represented in red. The discontinuity at P$\sim 0.5$ denotes the change between positive and negative values of local heritability.

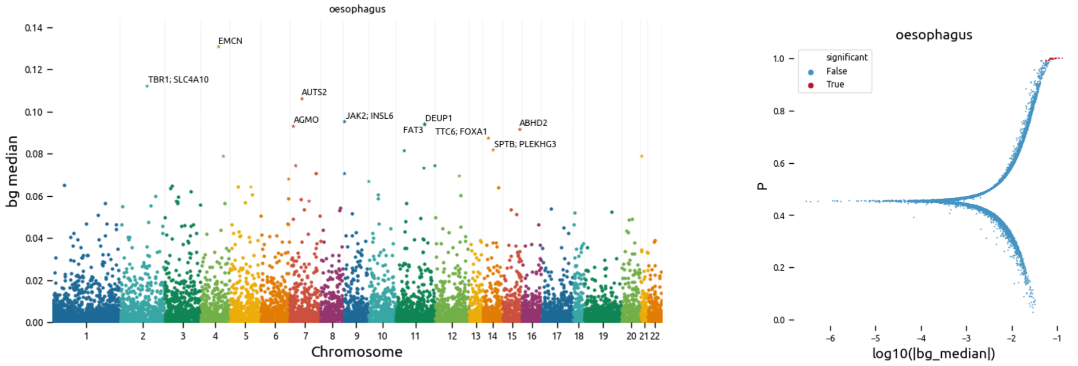

Supplementary Figure 17: **BAGHERA results for C15: Malignant neoplasms of oesophagus**. On the left we show a Manhattan plot of the local heritability $h_k^2$ (y-axis bg median in the implementation) for each gene in the analisis (x-axis, genes are sorted by position). Star markers denote significant genes, we report the names of the top scoring genes. On the right, we show the relationship between the local heritability estimates and the testing results. For each gene, on the x-axis we report the $log_{10}|h_k^2|$ and on the y-axis we report the indicator function value $\eta$, respectively bg median and P in the implementation. Significant genes are represented in red. The discontinuity at P$\sim 0.5$ denotes the change between positive and negative values of local heritability.

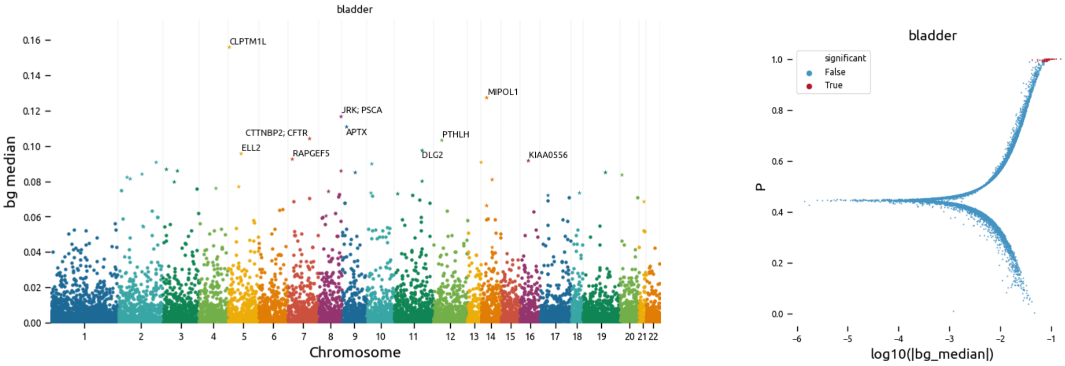

Supplementary Figure 18: **BAGHERA results for C67: Malignant neoplasms of bladder**. On the left we show a Manhattan plot of the local heritability $h_k^2$ (y-axis bg median in the implementation) for each gene in the analsis (x-axis, genes are sorted by position). Star markers denote significant genes, we report the names of the top scoring genes. On the right, we show the relationship between the local heritability estimates and the testing results. For each gene, on the x-axis we report the $log_{10}|h_k^2|$ and on the y-axis we report the indicator function value $\eta$, respectively bg median and P in the implementation. Significant genes are represented in red. The discontinuity at P$\sim$ $0.5$ denotes the change between positive and negative values of local heritability.

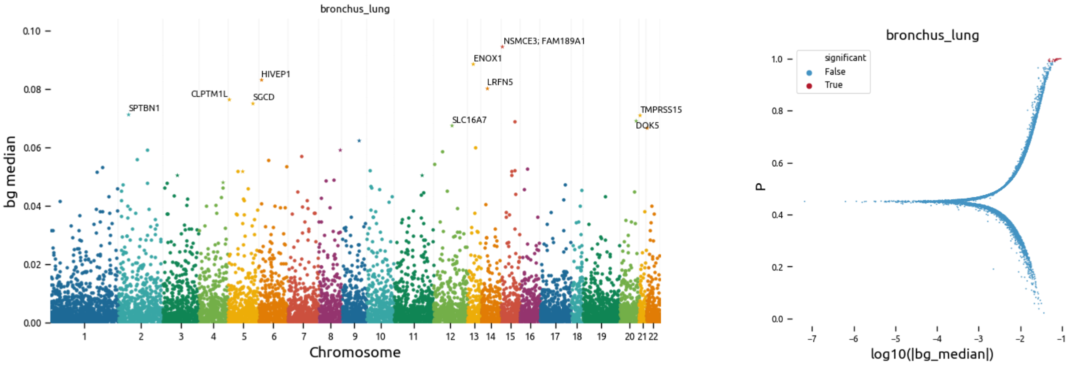

Supplementary Figure 19: **BAGHERA results for C34: Malignant neoplasms of brunchus and lung**. On the left we show a Manhattan plot of the local heritability $h_k^2$ (y-axis bg median in the implementation) for each gene in the analsis (x-axis, genes are sorted by position). Star markers denote significant genes, we report the names of the top scoring genes. On the right, we show the relationship between the local heritability estimates and the testing results. For each gene, on the x-axis we report the $log_{10}|h_k^2|$ and on the y-axis we report the indicator function value $\eta$, respectively bg median and P in the implementation. Significant genes are represented in red. The discontinuity at P$\sim$ $0.5$ denotes the change between positive and negative values of local heritability.

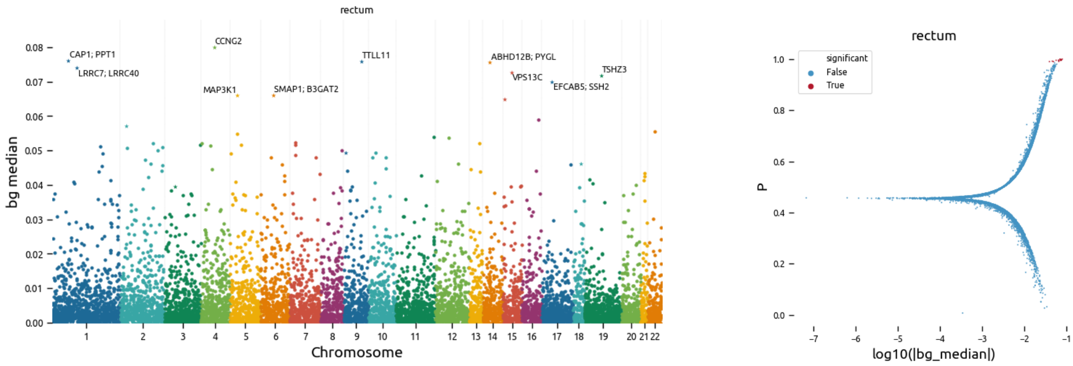

Supplementary Figure 20: **BAGHERA results for C20: Malignant neoplasms of rectum**. On the left we show a Manhattan plot of the local heritability $h_k^2$ (y-axis bg median in the implementation) for each gene in the analsis (x-axis, genes are sorted by position). Star markers denote significant genes, we report the names of the top scoring genes. On the right, we show the relationship between the local heritability estimates and the testing results. For each gene, on the x-axis we report the $log_{10}|h_k^2|$ and on the y-axis we report the indicator function value $\eta$, respectively bg median and P in the implementation. Significant genes are represented in red. The discontinuity at P$\sim 0.5$ denotes the change between positive and negative values of local heritability.

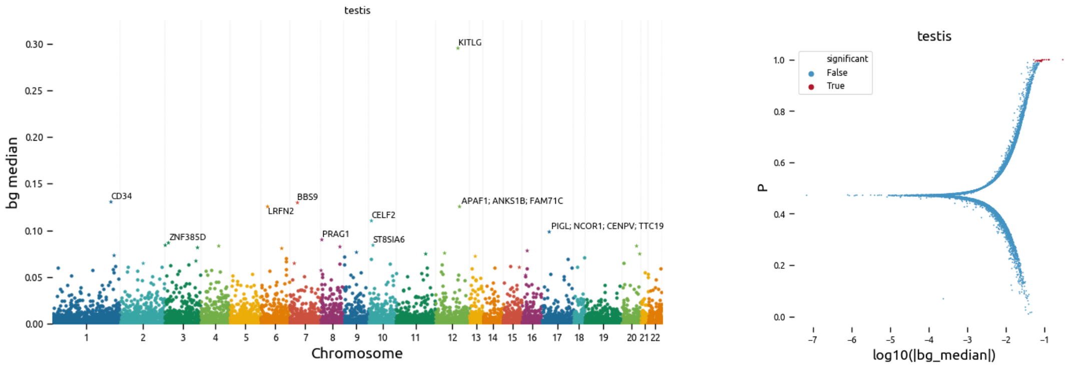

Supplementary Figure 21: **BAGHERA results for C62: Malignant neoplasms of testis**. On the left we show a Manhattan plot of the local heritability $h_k^2$ (y-axis bg median in the implementation) for each gene in the analsis (x-axis, genes are sorted by position). Star markers denote significant genes, we report the names of the top scoring genes. On the right, we show the relationship between the local heritability estimates and the testing results. For each gene, on the x-axis we report the $log_{10}|h_k^2|$ and on the y-axis we report the indicator function value $\eta$, respectively bg median and P in the implementation. Significant genes are represented in red. The discontinuity at P$\sim 0.5$ denotes the change between positive and negative values of local heritability.

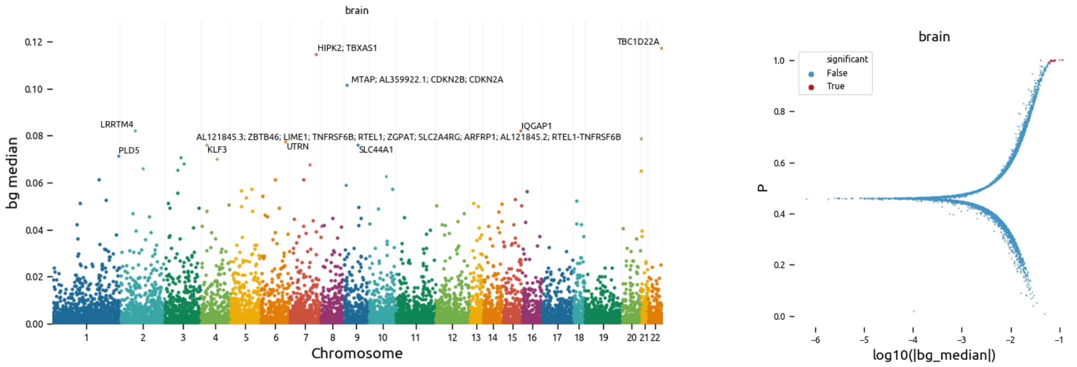

Supplementary Figure 22: **BAGHERA results for C71: Malignant neoplasms of brain**. On the left we show a Manhattan plot of the local heritability $h_k^2$ (y-axis bg median in the implementation) for each gene in the analsis (x-axis, genes are sorted by position). Star markers denote significant genes, we report the names of the top scoring genes. On the right, we show the relationship between the local heritability estimates and the testing results. For each gene, on the x-axis we report the $log_{10}|h_k^2|$ and on the y-axis we report the indicator function value $\eta$, respectively bg median and P in the implementation. Significant genes are represented in red. The discontinuity at P$\sim 0.5$ denotes the change between positive and negative values of local heritability.

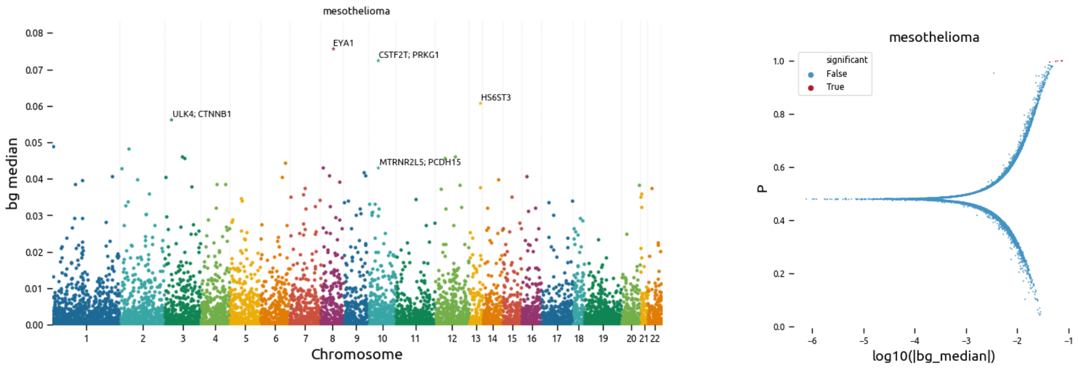

Supplementary Figure 23: **BAGHERA results for C45: Mesothelioma**. On the left we show a Manhattan plot of the local heritability $h_k^2$ (y-axis bg median in the implementation) for each gene in the analsis (x-axis, genes are sorted by position). Star markers denote significant genes, we report the names of the top scoring genes. On the right, we show the relationship between the local heritability estimates and the testing results. For each gene, on the x-axis we report the $log_{10}|h_k^2|$ and on the y-axis we report the indicator function value $\eta$, respectively bg median and P in the implementation. Significant genes are represented in red. The discontinuity at P$\sim 0.5$ denotes the change between positive and negative values of local heritability.

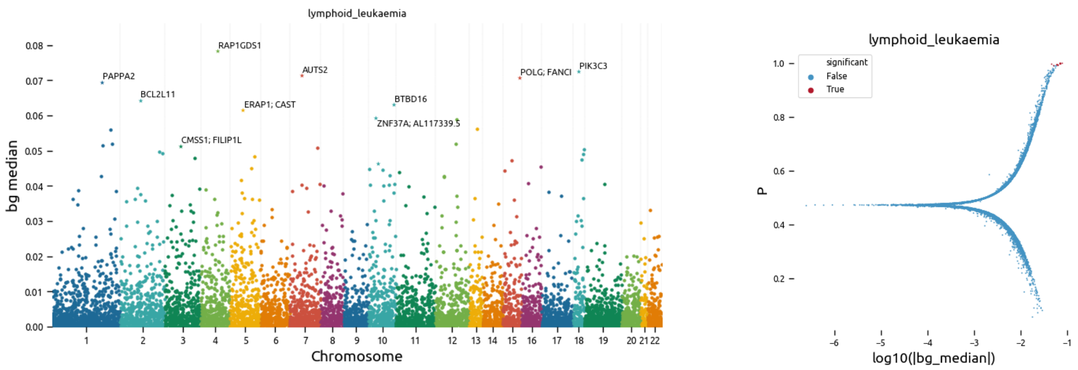

Supplementary Figure 24: **BAGHERA results for C91: Lymphiod Leukaemia**. On the left we show a Manhattan plot of the local heritability $h_k^2$ (y-axis bg median in the implementation) for each gene in the analsis (x-axis, genes are sorted by position). Star markers denote significant genes, we report the names of the top scoring genes. On the right, we show the relationship between the local heritability estimates and the testing results. For each gene, on the x-axis we report the $log_{10}|h_k^2|$ and on the y-axis we report the indicator function value $\eta$, respectively bg median and P in the implementation. Significant genes are represented in red. The discontinuity at P$\sim 0.5$ denotes the change between positive and negative values of local heritability.

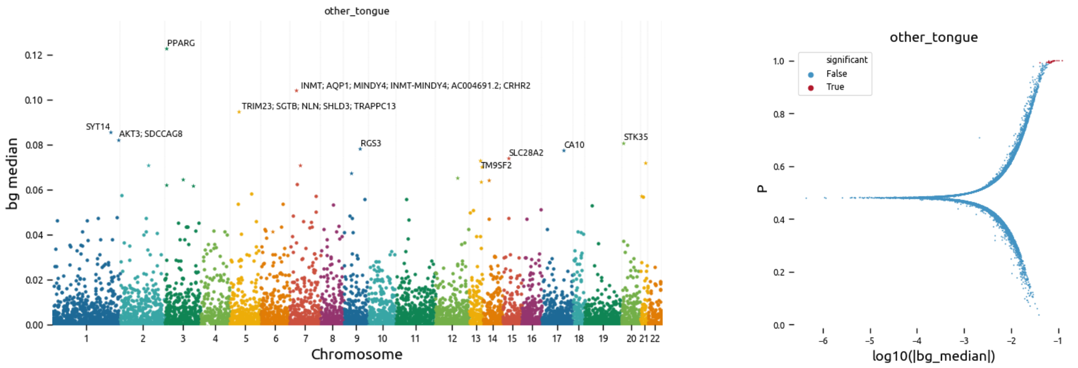

Supplementary Figure 25: **BAGHERA results for C02: Malignant neoplasms of other and unspecified parts of tongue**. On the left we show a Manhattan plot of the local heritability $h_k^2$ (y-axis bg median in the implementation) for each gene in the analsis (x-axis, genes are sorted by position). Star markers denote significant genes, we report the names of the top scoring genes. On the right, we show the relationship between the local heritability estimates and the testing results. For each gene, on the x-axis we report the $log_{10}|h_k^2|$ and on the y-axis we report the indicator function value $\eta$, respectively bg median and P in the implementation. Significant genes are represented in red. The discontinuity at P$\sim 0.5$ denotes the change between positive and negative values of local heritability.

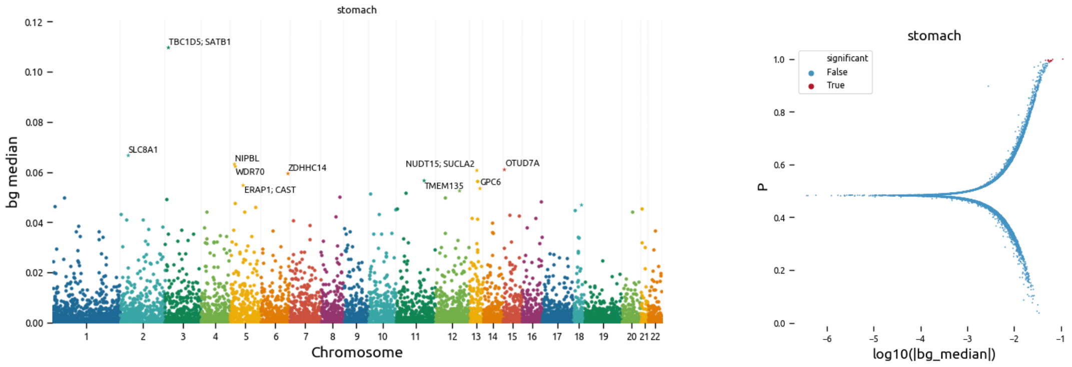

Supplementary Figure 26: **BAGHERA results for C61: Malignant neoplasms of stomach**. On the left we show a Manhattan plot of the local heritability $h_k^2$ (y-axis bg median in the implementation) for each gene in the analisis (x-axis, genes are sorted by position). Star markers denote significant genes, we report the names of the top scoring genes. On the right, we show the relationship between the local heritability estimates and the testing results. For each gene, on the x-axis we report the $log_{10}|h_k^2|$ and on the y-axis we report the indicator function value $\eta$, respectively bg median and P in the implementation. Significant genes are represented in red. The discontinuity at P$\sim 0.5$ denotes the change between positive and negative values of local heritability.

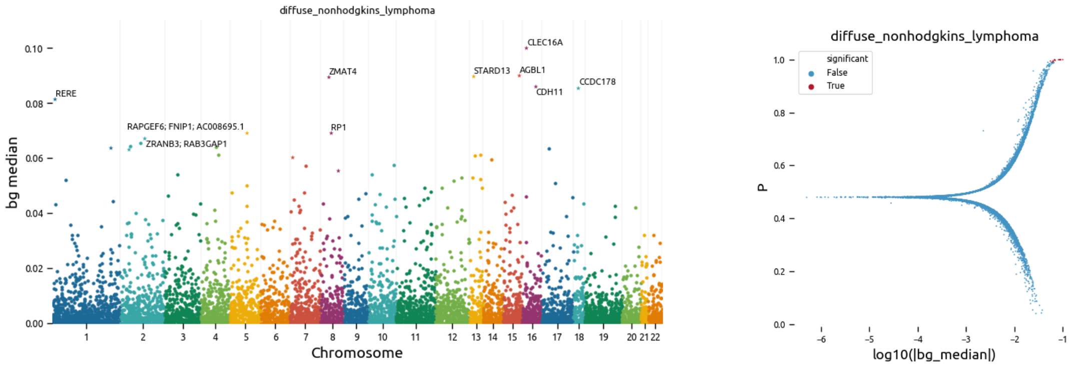

Supplementary Figure 27: **BAGHERA results for C83: Diffuse non-Hodgkin's lymphoma**. On the left we show a Manhattan plot of the local heritability $h_k^2$ (y-axis bg median in the implementation) for each gene in the analisis (x-axis, genes are sorted by position). Star markers denote significant genes, we report the names of the top scoring genes. On the right, we show the relationship between the local heritability estimates and the testing results. For each gene, on the x-axis we report the $log_{10}|h_k^2|$ and on the y-axis we report the indicator function value $\eta$, respectively bg median and P in the implementation. Significant genes are represented in red. The discontinuity at P$\sim 0.5$ denotes the change between positive and negative values of local heritability.

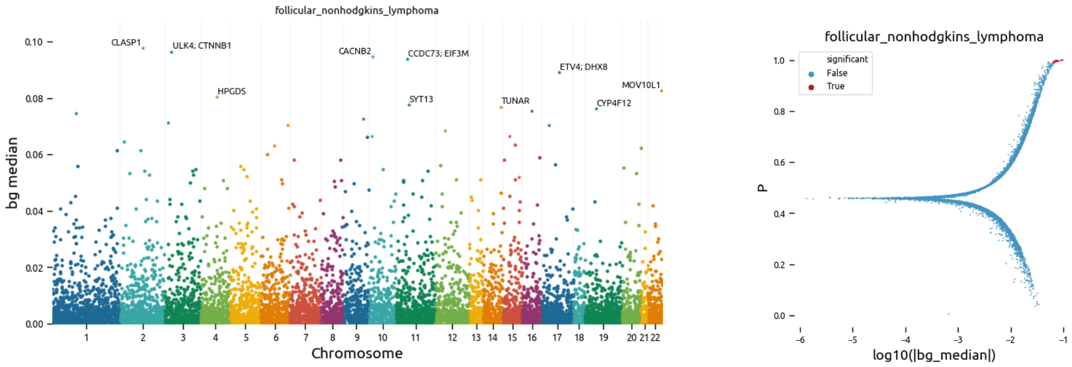

Supplementary Figure 28: **BAGHERA results for C82: Follicular non-Hodgkin's lymphoma**. On the left we show a Manhattan plot of the local heritability $h_k^2$ (y-axis bg median in the implementation) for each gene in the analsis (x-axis, genes are sorted by position). Star markers denote significant genes, we report the names of the top scoring genes. On the right, we show the relationship between the local heritability estimates and the testing results. For each gene, on the x-axis we report the $log_{10}|h_k^2|$ and on the y-axis we report the indicator function value $\eta$, respectively bg median and P in the implementation. Significant genes are represented in red. The discontinuity at P$\sim 0.5$ denotes the change between positive and negative values of local heritability.

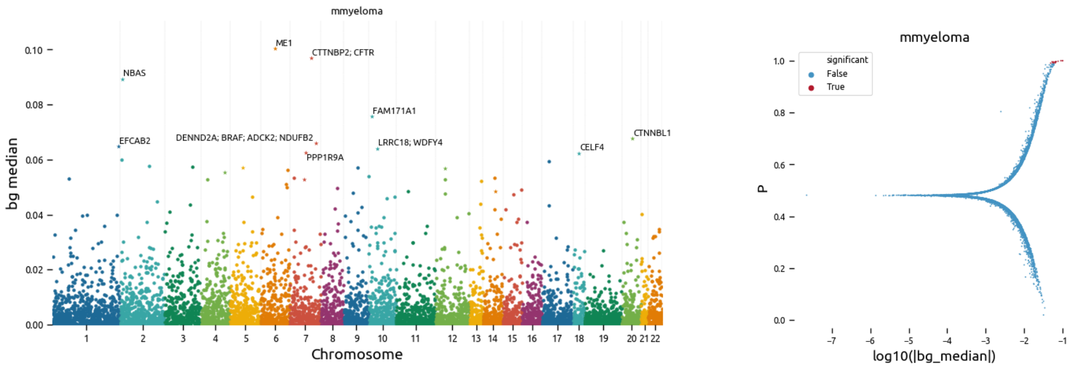

Supplementary Figure 29: **BAGHERA results for C90: Multiple Myeloma**. On the left we show a Manhattan plot of the local heritability $h_k^2$ (y-axis bg median in the implementation) for each gene in the analsis (x-axis, genes are sorted by position). Star markers denote significant genes, we report the names of the top scoring genes. On the right, we show the relationship between the local heritability estimates and the testing results. For each gene, on the x-axis we report the $log_{10}|h_k^2|$ and on the y-axis we report the indicator function value $\eta$, respectively bg median and P in the implementation. Significant genes are represented in red. The discontinuity at P$\sim 0.5$ denotes the change between positive and negative values of local heritability.

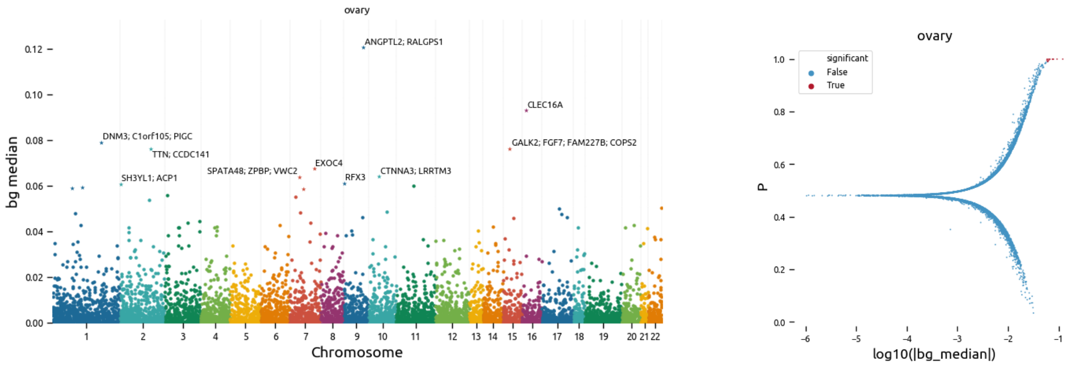

Supplementary Figure 30: **BAGHERA results for C56: Malignant neoplasm of ovary**. On the left we show a Manhattan plot of the local heritability $h_k^2$ (y-axis bg median in the implementation) for each gene in the analsis (x-axis, genes are sorted by position). Star markers denote significant genes, we report the names of the top scoring genes. On the right, we show the relationship between the local heritability estimates and the testing results. For each gene, on the x-axis we report the $log_{10}|h_k^2|$ and on the y-axis we report the indicator function value $\eta$, respectively bg median and P in the implementation. Significant genes are represented in red. The discontinuity at P$\sim 0.5$ denotes the change between positive and negative values of local heritability.

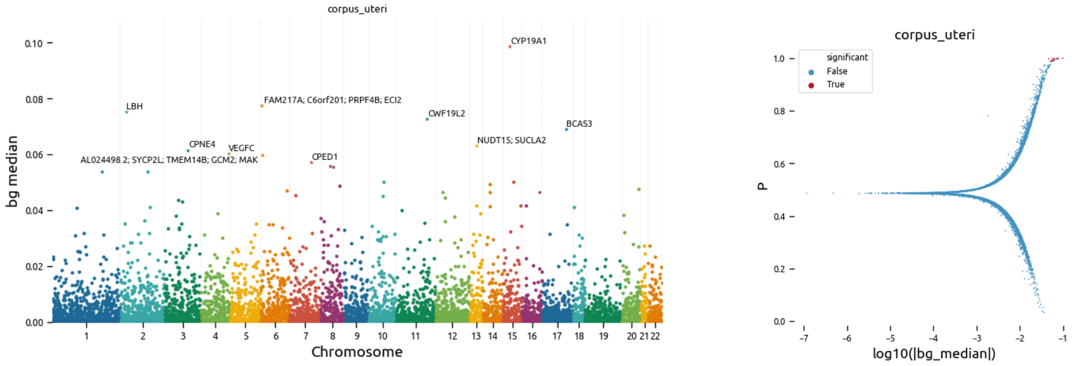

Supplementary Figure 31: **BAGHERA results for C54: Malignant neoplasm of corpus uteri.** On the left we show a Manhattan plot of the local heritability $h_k^2$ (y-axis bg median in the implementation) for each gene in the analsis (x-axis, genes are sorted by position). Star markers denote significant genes, we report the names of the top scoring genes. On the right, we show the relationship between the local heritability estimates and the testing results. For each gene, on the x-axis we report the $log_{10}|h_k^2|$ and on the y-axis we report the indicator function value $\eta$, respectively bg median and P in the implementation. Significant genes are represented in red. The discontinuity at P$\sim 0.5$ denotes the change between positive and negative values of local heritability.

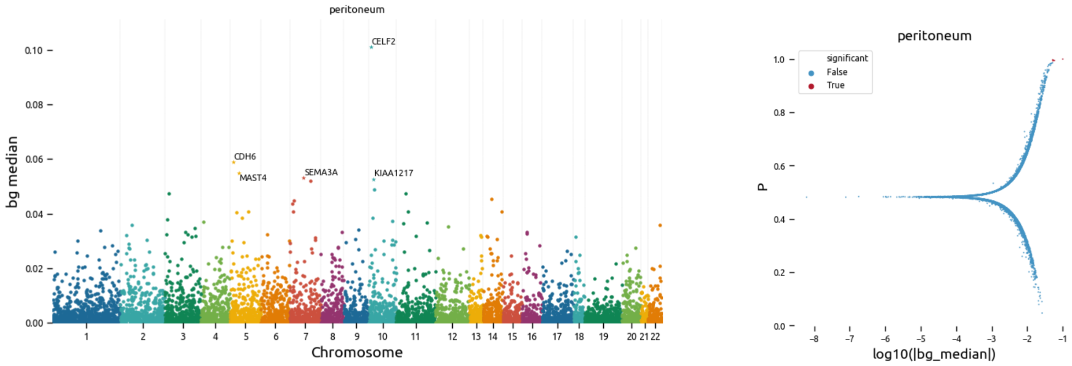

Supplementary Figure 32: **BAGHERA results for C48: Malignant neoplasm of retroperitoneum and peritoneum.** On the left we show a Manhattan plot of the local heritability $h_k^2$ (y-axis bg median in the implementation) for each gene in the analsis (x-axis, genes are sorted by position). Star markers denote significant genes, we report the names of the top scoring genes. On the right, we show the relationship between the local heritability estimates and the testing results. For each gene, on the x-axis we report the $log_{10}|h_k^2|$ and on the y-axis we report the indicator function value $\eta$, respectively bg median and P in the implementation. Significant genes are represented in red. The discontinuity at P$\sim 0.5$ denotes the change between positive and negative values of local heritability.

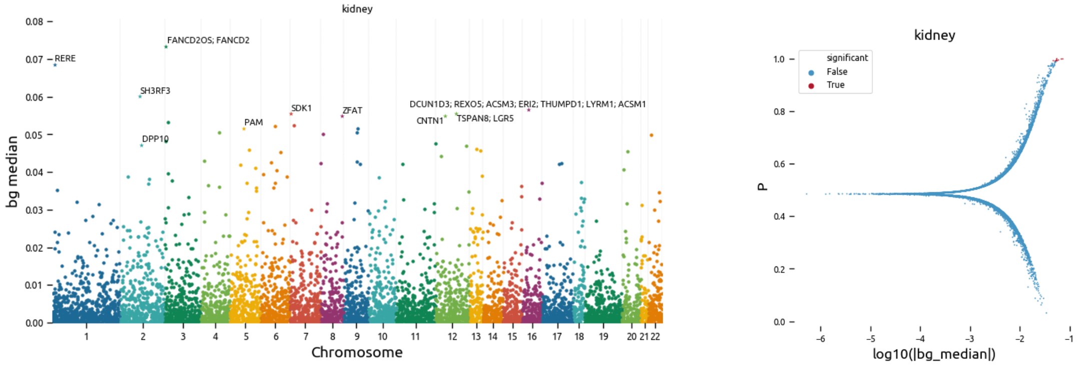

Supplementary Figure 33: **BAGHERA results for C64: Malignant neoplasm of kidney except renal pelvis.** On the left we show a Manhattan plot of the local heritability $h_k^2$ (y-axis bg median in the implementation) for each gene in the analsis (x-axis, genes are sorted by position). Star markers denote significant genes, we report the names of the top scoring genes. On the right, we show the relationship between the local heritability estimates and the testing results. For each gene, on the x-axis we report the $log_{10}|h_k^2|$ and on the y-axis we report the indicator function value $\eta$, respectively bg median and P in the implementation. Significant genes are represented in red. The discontinuity at P$\sim 0.5$ denotes the change between positive and negative values of local heritability.

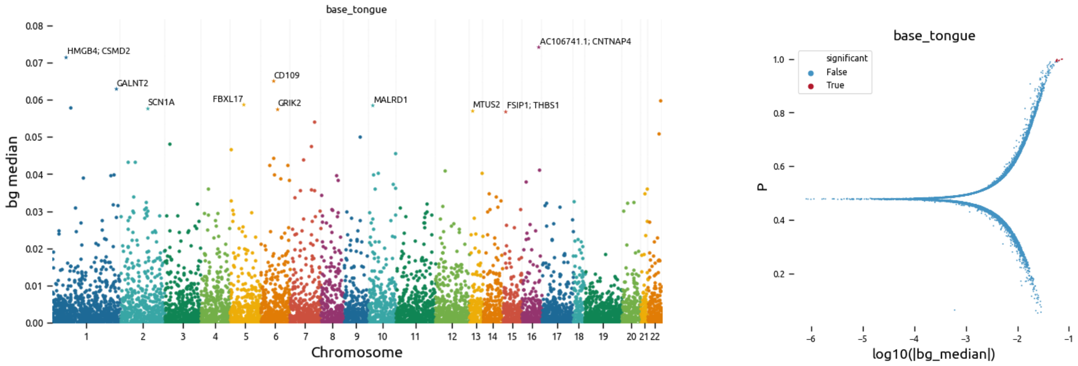

Supplementary Figure 34: **BAGHERA results for C01: Malignant neoplasm of base of tongue**. On the left we show a Manhattan plot of the local heritability $h_k^2$ (y-axis bg median in the implementation) for each gene in the analsis (x-axis, genes are sorted by position). Star markers denote significant genes, we report the names of the top scoring genes. On the right, we show the relationship between the local heritability estimates and the testing results. For each gene, on the x-axis we report the $log_{10}|h_k^2|$ and on the y-axis we report the indicator function value $\eta$, respectively bg median and P in the implementation. Significant genes are represented in red. The discontinuity at P$\sim$ $0.5$ denotes the change between positive and negative values of local heritability.

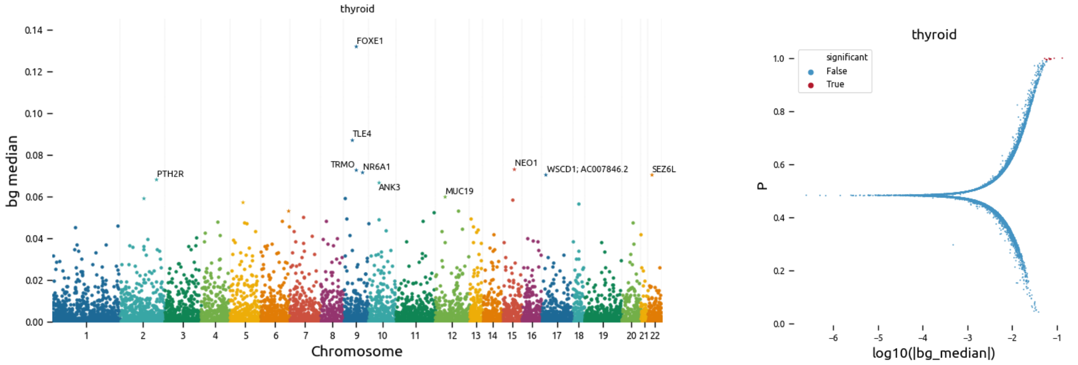

Supplementary Figure 35: **BAGHERA results for C73: Malignant neoplasm of thyroid gland**. On the left we show a Manhattan plot of the local heritability $h_k^2$ (y-axis bg median in the implementation) for each gene in the analsis (x-axis, genes are sorted by position). Star markers denote significant genes, we report the names of the top scoring genes. On the right, we show the relationship between the local heritability estimates and the testing results. For each gene, on the x-axis we report the $log_{10}|h_k^2|$ and on the y-axis we report the indicator function value $\eta$, respectively bg median and P in the implementation. Significant genes are represented in red. The discontinuity at P$\sim$ $0.5$ denotes the change between positive and negative values of local heritability.

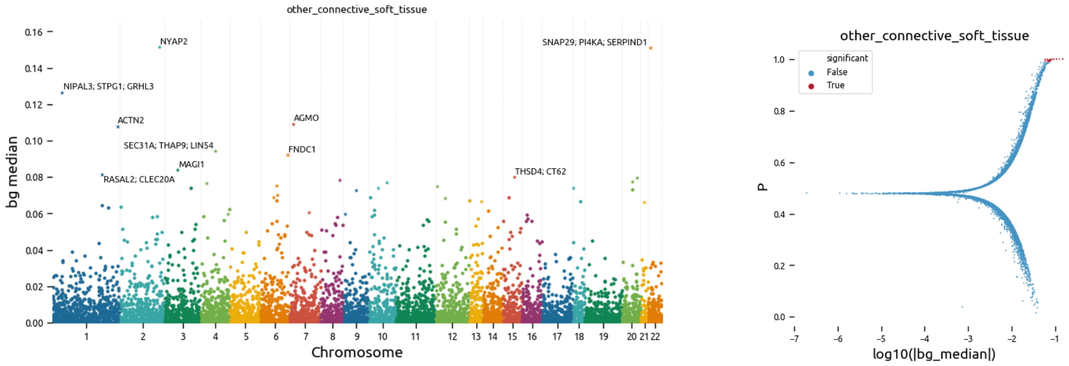

Supplementary Figure 36: **BAGHERA results for C49: Malignant neoplasm of other connective and soft tissue.** On the left we show a Manhattan plot of the local heritability $h_k^2$ (y-axis bg median in the implementation) for each gene in the analsis (x-axis, genes are sorted by position). Star markers denote significant genes, we report the names of the top scoring genes. On the right, we show the relationship between the local heritability estimates and the testing results. For each gene, on the x-axis we report the $log_{10}|h_k^2|$ and on the y-axis we report the indicator function value $\eta$, respectively bg median and P in the implementation. Significant genes are represented in red. The discontinuity at P$\sim 0.5$ denotes the change between positive and negative values of local heritability.

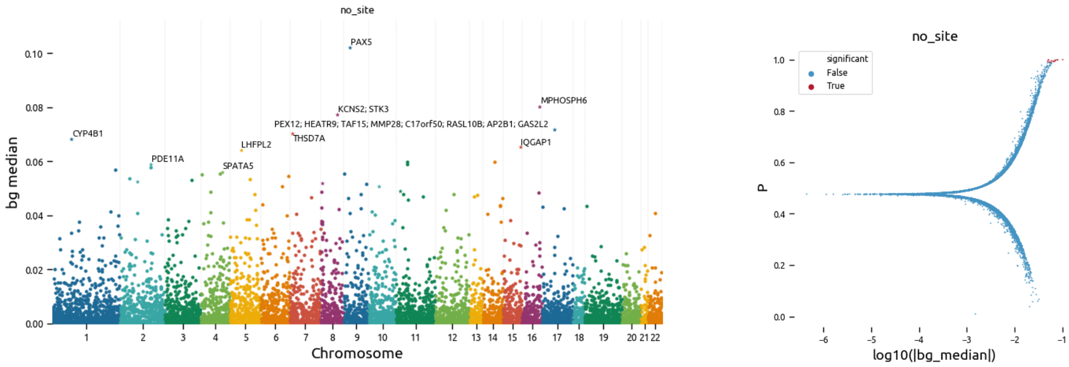

Supplementary Figure 37: **BAGHERA results for C80: Malignant neoplasm without specification of site.** On the left we show a Manhattan plot of the local heritability $h_k^2$ (y-axis bg median in the implementation) for each gene in the analsis (x-axis, genes are sorted by position). Star markers denote significant genes, we report the names of the top scoring genes. On the right, we show the relationship between the local heritability estimates and the testing results. For each gene, on the x-axis we report the $log_{10}|h_k^2|$ and on the y-axis we report the indicator function value $\eta$, respectively bg median and P in the implementation. Significant genes are represented in red. The discontinuity at P$\sim 0.5$ denotes the change between positive and negative values of local heritability.

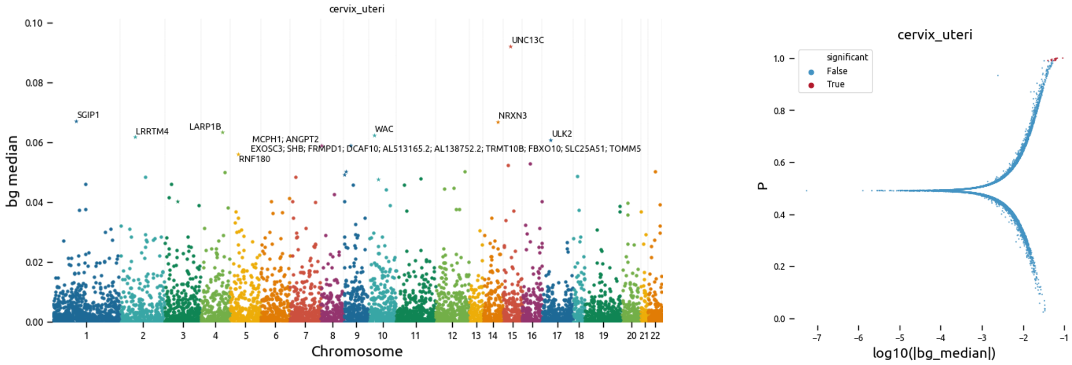

Supplementary Figure 38: **BAGHERA results for C53: Malignant neoplasm of cervix utery**. On the left we show a Manhattan plot of the local heritability $h_k^2$ (y-axis bg median in the implementation) for each gene in the analsis (x-axis, genes are sorted by position). Star markers denote significant genes, we report the names of the top scoring genes. On the right, we show the relationship between the local heritability estimates and the testing results. For each gene, on the x-axis we report the $log_{10}|h_k^2|$ and on the y-axis we report the indicator function value $\eta$, respectively bg median and P in the implementation. Significant genes are represented in red. The discontinuity at P$\sim 0.5$ denotes the change between positive and negative values of local heritability.

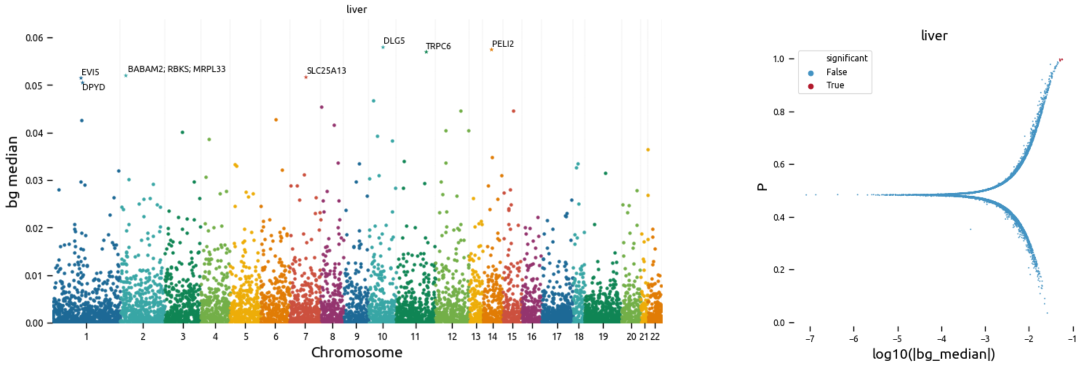

Supplementary Figure 39: **BAGHERA results for C22: Malignant neoplasm of liver and intrahepatic bile ducts.** On the left we show a Manhattan plot of the local heritability $h_k^2$ (y-axis bg median in the implementation) for each gene in the analsis (x-axis, genes are sorted by position). Star markers denote significant genes, we report the names of the top scoring genes. On the right, we show the relationship between the local heritability estimates and the testing results. For each gene, on the x-axis we report the $log_{10}|h_k^2|$ and on the y-axis we report the indicator function value $\eta$, respectively bg median and P in the implementation. Significant genes are represented in red. The discontinuity at P$\sim 0.5$ denotes the change between positive and negative values of local heritability.

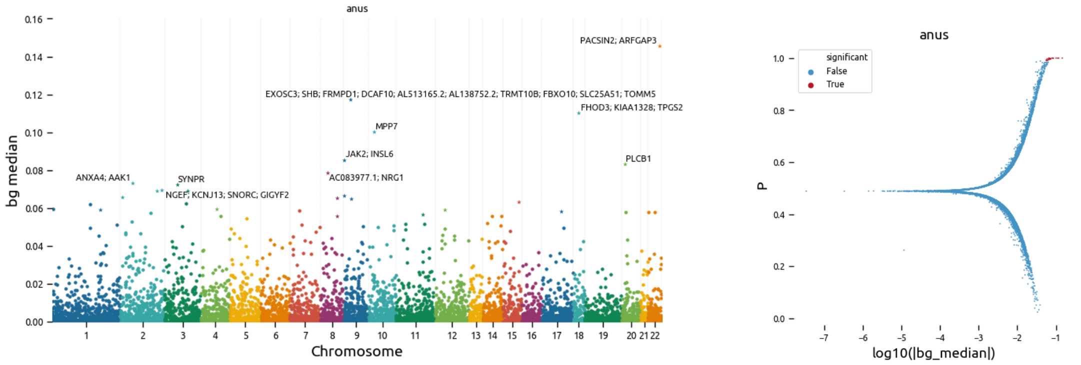

Supplementary Figure 40: **BAGHERA results for C21: Malignant neoplasm of anus and anal canal.** On the left we show a Manhattan plot of the local heritability $h_k^2$ (y-axis bg median in the implementation) for each gene in the analsis (x-axis, genes are sorted by position). Star markers denote significant genes, we report the names of the top scoring genes. On the right, we show the relationship between the local heritability estimates and the testing results. For each gene, on the x-axis we report the $log_{10}|h_k^2|$ and on the y-axis we report the indicator function value $\eta$, respectively bg median and P in the implementation. Significant genes are represented in red. The discontinuity at P$\sim$ $0.5$ denotes the change between positive and negative values of local heritability.

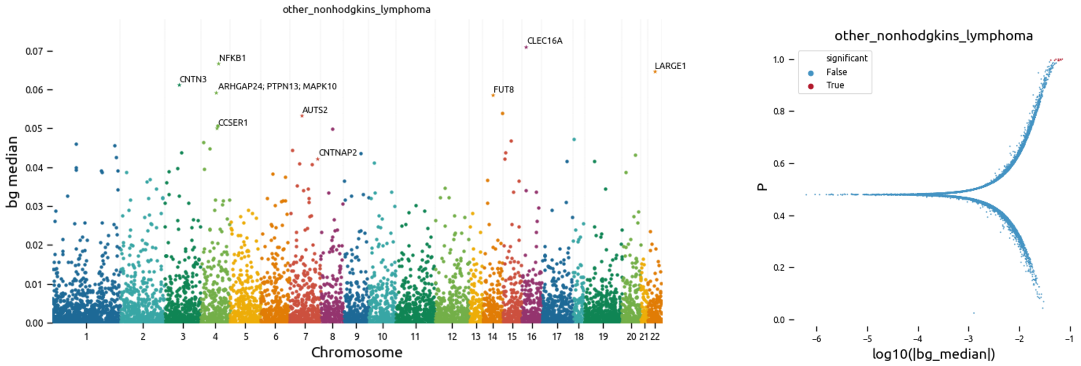

Supplementary Figure 41: **BAGHERA results for C85: Other and unspecified types of non-Hodgkins's lymphoma**. On the left we show a Manhattan plot of the local heritability $h_k^2$ (y-axis bg median in the implementation) for each gene in the analsis (x-axis, genes are sorted by position). Star markers denote significant genes, we report the names of the top scoring genes. On the right, we show the relationship between the local heritability estimates and the testing results. For each gene, on the x-axis we report the $log_{10}|h_k^2|$ and on the y-axis we report the indicator function value $\eta$, respectively bg median and P in the implementation. Significant genes are represented in red. The discontinuity at P$\sim 0.5$ denotes the change between positive and negative values of local heritability.

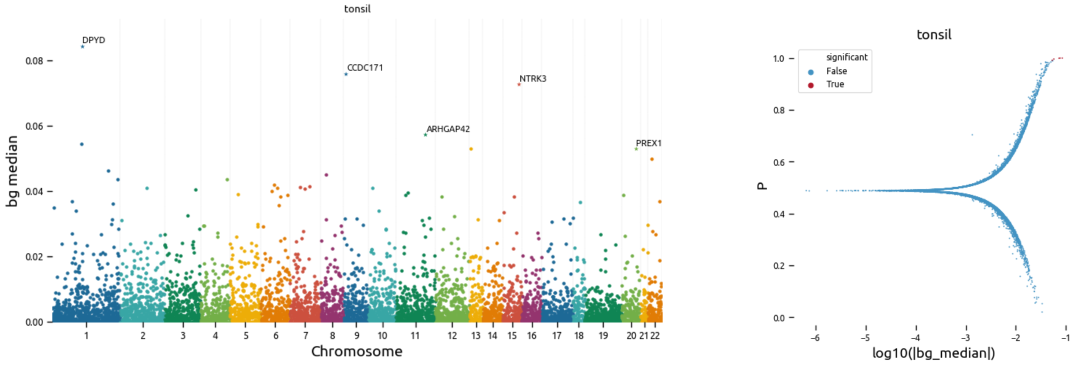

Supplementary Figure 42: **BAGHERA results for C09: Malignant neoplasm of tonsil**. On the left we show a Manhattan plot of the local heritability $h_k^2$ (y-axis bg median in the implementation) for each gene in the analsis (x-axis, genes are sorted by position). Star markers denote significant genes, we report the names of the top scoring genes. On the right, we show the relationship between the local heritability estimates and the testing results. For each gene, on the x-axis we report the $log_{10}|h_k^2|$ and on the y-axis we report the indicator function value $\eta$, respectively bg median and P in the implementation. Significant genes are represented in red. The discontinuity at P$\sim 0.5$ denotes the change between positive and negative values of local heritability.

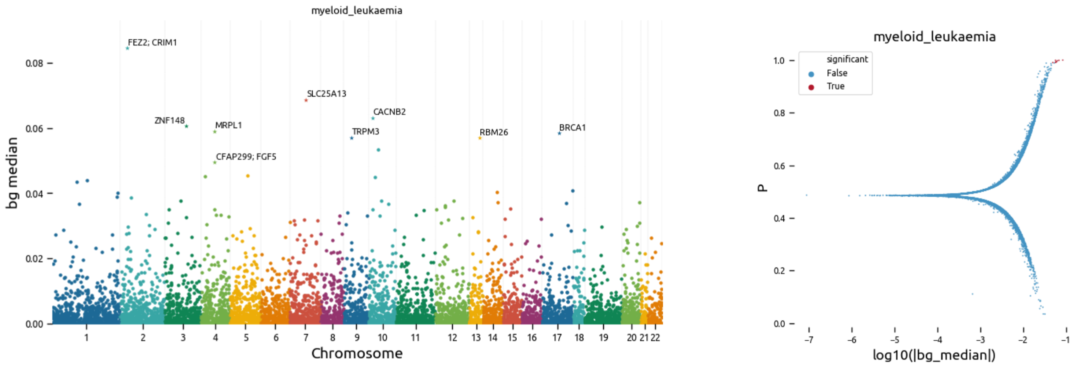

Supplementary Figure 43: **BAGHERA results for C92: Myeloid Leukaemia**. On the left we show a Manhattan plot of the local heritability $h^2_k$ (y-axis bg median in the implementation) for each gene in the analisis (x-axis, genes are sorted by position). Star markers denote significant genes, we report the names of the top scoring genes. On the right, we show the relationship between the local heritability estimates and the testing results. For each gene, on the x-axis we report the $log_{10}|h^2_k|$ and on the y-axis we report the indicator function value $\eta$, respectively bg median and P in the implementation. Significant genes are represented in red. The discontinuity at P$\sim 0.5$ denotes the change between positive and negative values of local heritability.

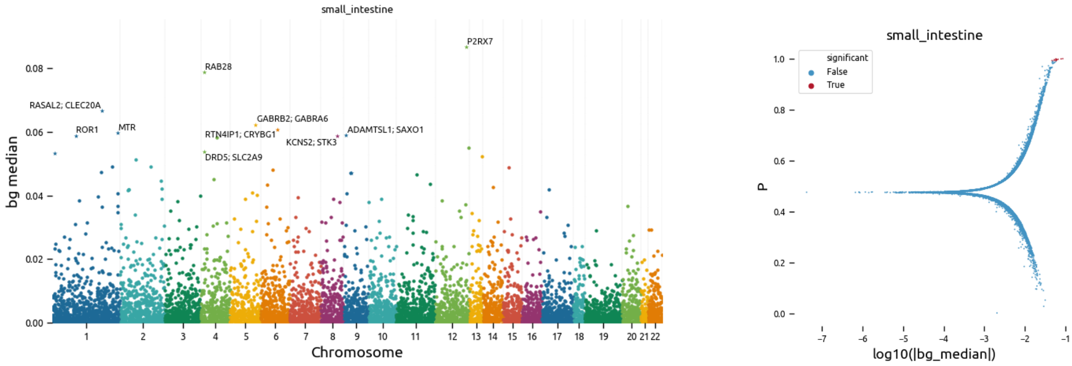

Supplementary Figure 44: **BAGHERA results for C17: Malignant neoplasm of small intestine.** On the left we show a Manhattan plot of the local heritability $h^2_k$ (y-axis bg median in the implementation) for each gene in the analisis (x-axis, genes are sorted by position). Star markers denote significant genes, we report the names of the top scoring genes. On the right, we show the relationship between the local heritability estimates and the testing results. For each gene, on the x-axis we report the $log_{10}|h^2_k|$ and on the y-axis we report the indicator function value $\eta$, respectively bg median and P in the implementation. Significant genes are represented in red. The discontinuity at P$\sim 0.5$ denotes the change between positive and negative values of local heritability.

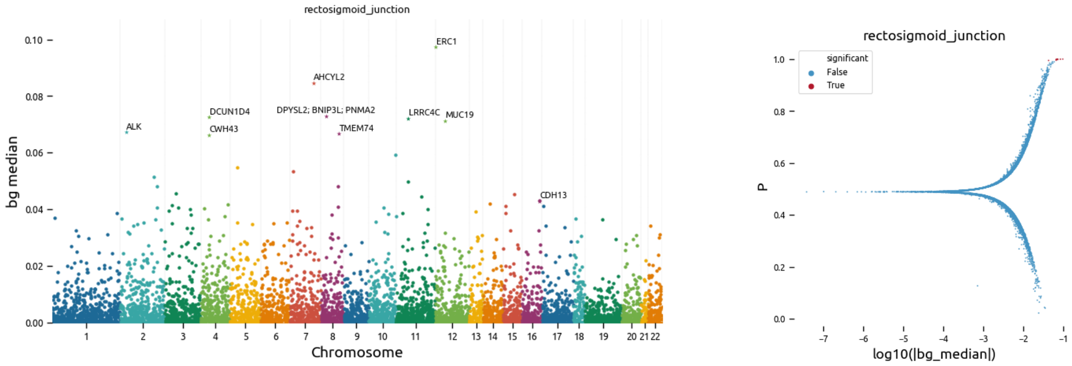

Supplementary Figure 45: **BAGHERA results for C19: Malignant neoplasm of rectosigmoid junction**. On the left we show a Manhattan plot of the local heritability $h_k^2$ (y-axis bg median in the implementation) for each gene in the analsis (x-axis, genes are sorted by position). Star markers denote significant genes, we report the names of the top scoring genes. On the right, we show the relationship between the local heritability estimates and the testing results. For each gene, on the x-axis we report the $log_{10}|h_k^2|$ and on the y-axis we report the indicator function value $\eta$, respectively bg median and P in the implementation. Significant genes are represented in red. The discontinuity at P$\sim 0.5$ denotes the change between positive and negative values of local heritability.

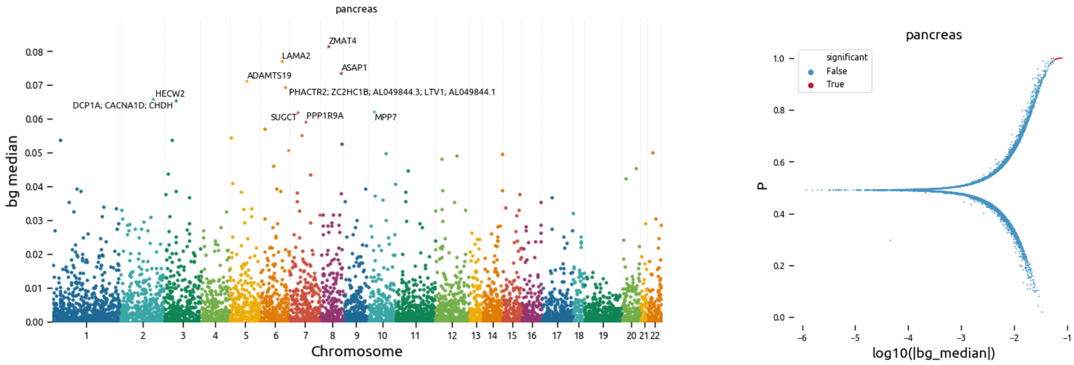

Supplementary Figure 46: **BAGHERA results for C25: Malignant neoplasm of pancreas**. On the left we show a Manhattan plot of the local heritability $h_k^2$ (y-axis bg median in the implementation) for each gene in the analsis (x-axis, genes are sorted by position). Star markers denote significant genes, we report the names of the top scoring genes. On the right, we show the relationship between the local heritability estimates and the testing results. For each gene, on the x-axis we report the $log_{10}|h_k^2|$ and on the y-axis we report the indicator function value $\eta$, respectively bg median and P in the implementation. Significant genes are represented in red. The discontinuity at P$\sim 0.5$ denotes the change between positive and negative values of local heritability.

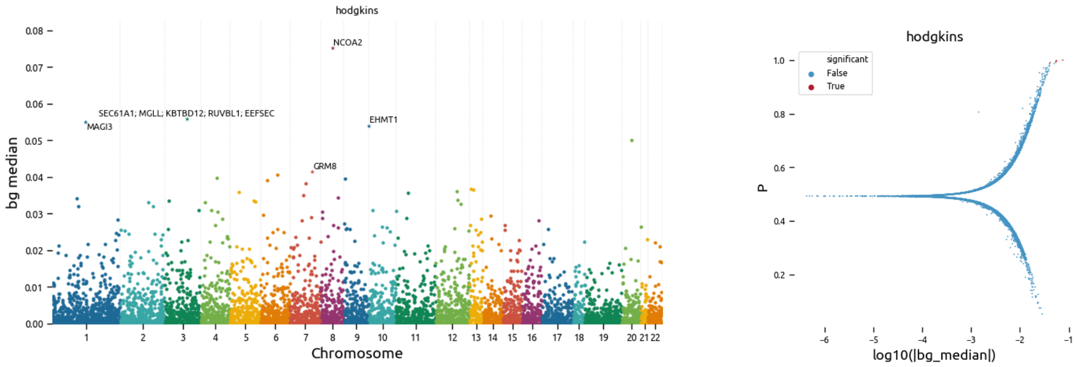

Supplementary Figure 47: **BAGHERA results for C81: Hodgkin's disease**. On the left we show a Manhattan plot of the local heritability $h_k^2$ (y-axis bg median in the implementation) for each gene in the analsis (x-axis, genes are sorted by position). Star markers denote significant genes, we report the names of the top scoring genes. On the right, we show the relationship between the local heritability estimates and the testing results. For each gene, on the x-axis we report the $log_{10}|h_k^2|$ and on the y-axis we report the indicator function value $\eta$, respectively bg median and P in the implementation. Significant genes are represented in red. The discontinuity at P$\sim 0.5$ denotes the change between positive and negative values of local heritability.

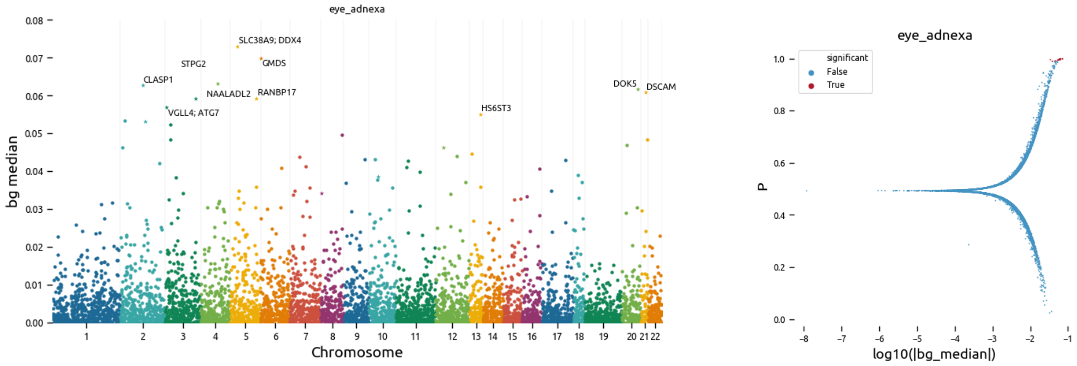

Supplementary Figure 48: **BAGHERA results for C69: Malignant neoplasm of eye and adnexa**. On the left we show a Manhattan plot of the local heritability $h_k^2$ (y-axis bg median in the implementation) for each gene in the analsis (x-axis, genes are sorted by position). Star markers denote significant genes, we report the names of the top scoring genes. On the right, we show the relationship between the local heritability estimates and the testing results. For each gene, on the x-axis we report the $log_{10}|h_k^2|$ and on the y-axis we report the indicator function value $\eta$, respectively bg median and P in the implementation. Significant genes are represented in red. The discontinuity at P$\sim 0.5$ denotes the change between positive and negative values of local heritability.

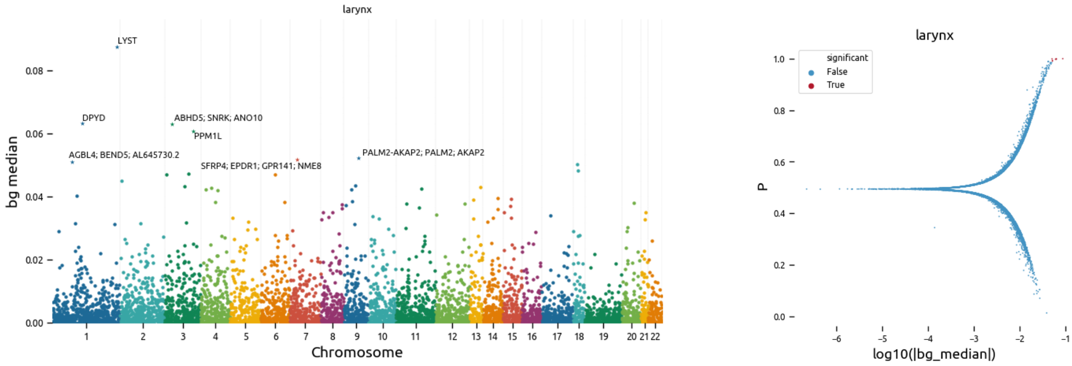

Supplementary Figure 49: **BAGHERA results for C32: Malignant neoplasm of larynx**. On the left we show a Manhattan plot of the local heritability $h_k^2$ (y-axis bg median in the implementation) for each gene in the analsis (x-axis, genes are sorted by position). Star markers denote significant genes, we report the names of the top scoring genes. On the right, we show the relationship between the local heritability estimates and the testing results. For each gene, on the x-axis we report the $log_{10}|h_k^2|$ and on the y-axis we report the indicator function value $\eta$, respectively bg median and P in the implementation. Significant genes are represented in red. The discontinuity at P$\sim 0.5$ denotes the change between positive and negative values of local heritability.

#### 4.7 Relationship between GWAS and BAGHERA results

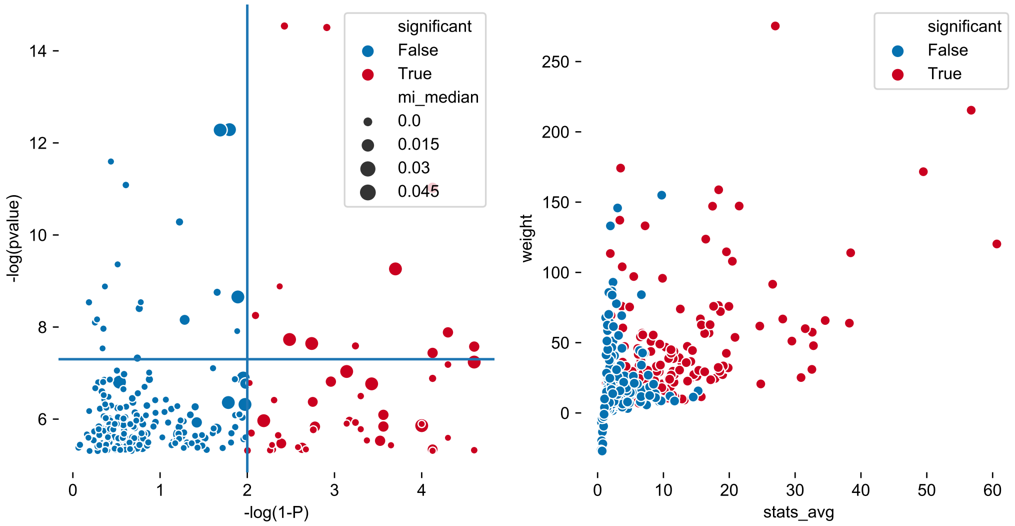

Supplementary Figure 50: **Relationship between GWAS and BAGHERA results.** On the left, we show the correlation between GWAS pvalues (x-axis, we consider only p-values$< 10^{-5}$) and BAGHERA's $\eta$ (x-axis, in the implementation $\eta$ is named P, and we transform the value as $1-\eta$ to be comparable to pvalues). For each gene analysed by BAGHERA, we selected the smallest p-value of its SNPs. Horizontal line is the GWAS significance threshold, pvalue $= 5 \times 10^{-8}$, vertical line is for $\eta = 0.99$. Size of the marker is proportional to the genomewide $h^2$ estimate (mi median). It's worth noting that there is no linear dependence of BAGHERA's significance and GWAS pvalues. In some cases, see top left quadrant, there are genes harboring SNPs with very small p-values, that are not significant for the heritability analysis. On the right, instead, we show the correlation between each gene average $\chi^2$ and local heritability (y-axis, to make results from different cancer types comparable we show the gene weight as $w_k = (h_k^2 - h^2)/h^2$). Significant genes are color coded in red. We can notice here that, as expected, there is correlation between the average statistics of a gene and local heritability.

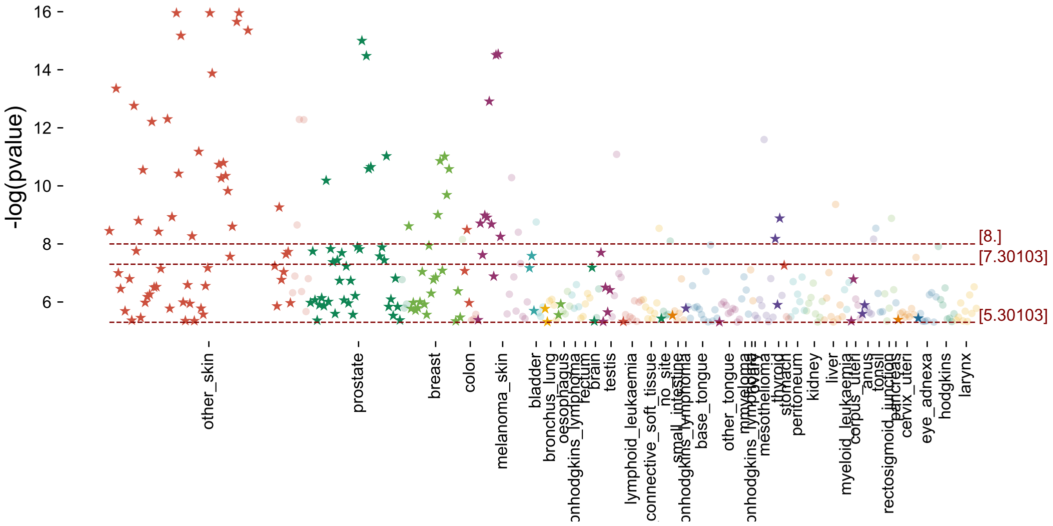

Supplementary Figure 51: **Single malignancy GWAS significant pvalues.** For each cancer type, color coded, we selected the genes harbouring a SNP with p-value $> 10^{-5}$ and we plotted their logarithm. In the x-axis, for each single malignancy, we sorted the genes by their $\eta$, from biggest to smallest. Genes that are significant for BAGHERA are dark stars, while those that are not significant are represented with dots. Horizontal lines are different p-value significance levels. This figure details the results above (Supp. Fig. 50)

#### 4.8 Complete Results on 38 cancers

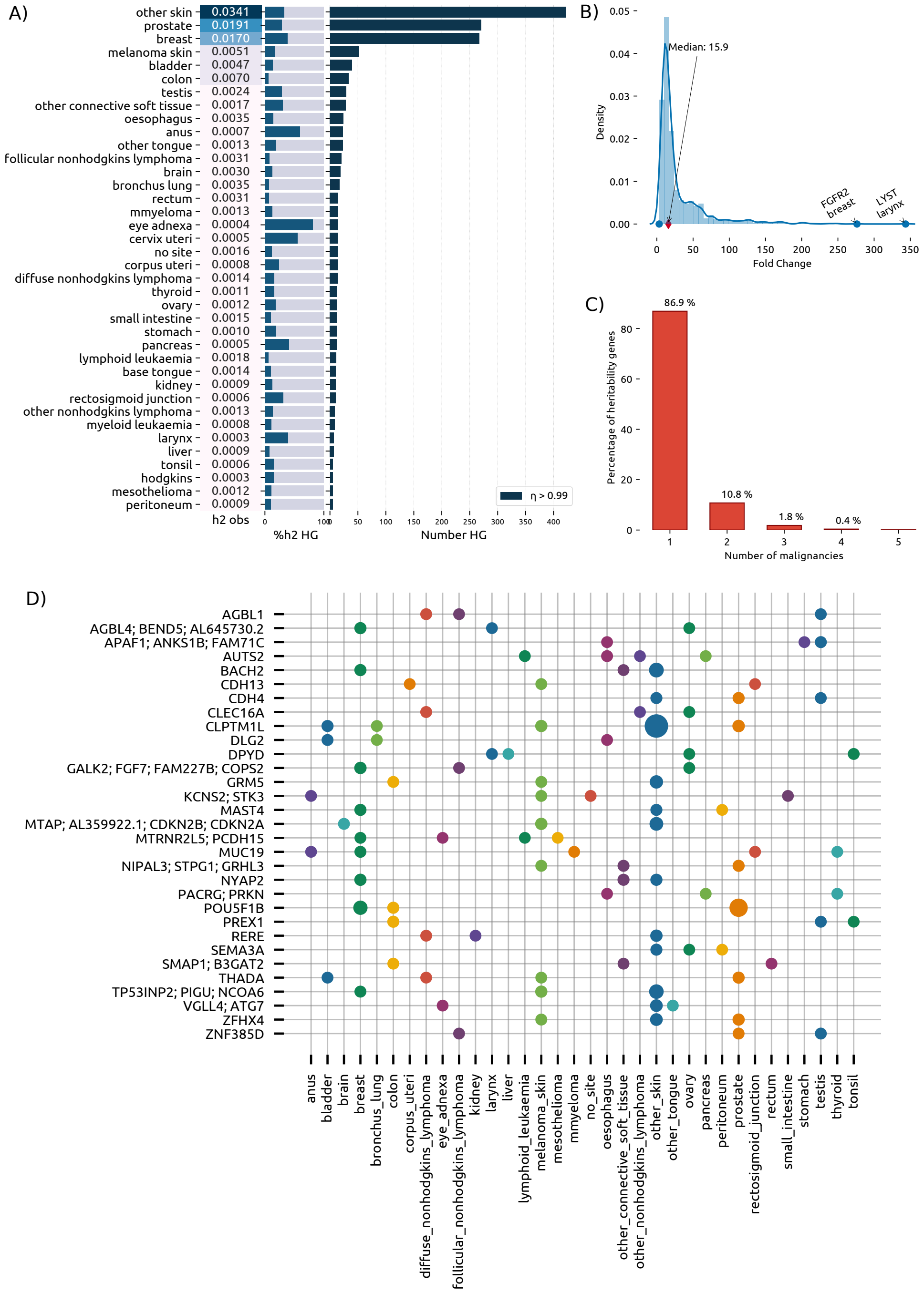

Supplementary Figure 52: **Heritability genes across $38$ cancers in the UK Biobank** A) For each malignancy we report the observed heritability ($h^2_{SNP}$, left box), the percentage of $h^2_{SNP}$ explained by heritability genes (central barplot, dark blue is the percentage explained by HGs) and the number of heritability genes (right barplot). B) Gene-level heritability density distribution across heritability genes, expressed as fold-change with respect to the genome-wide estimate. Highlighted are the top genes and the median fold-change across all cancers. C) Percentage of cancer heritability genes associated with multiple cancers. More than $10\%$ of CHGs are common to multiple malignancies.

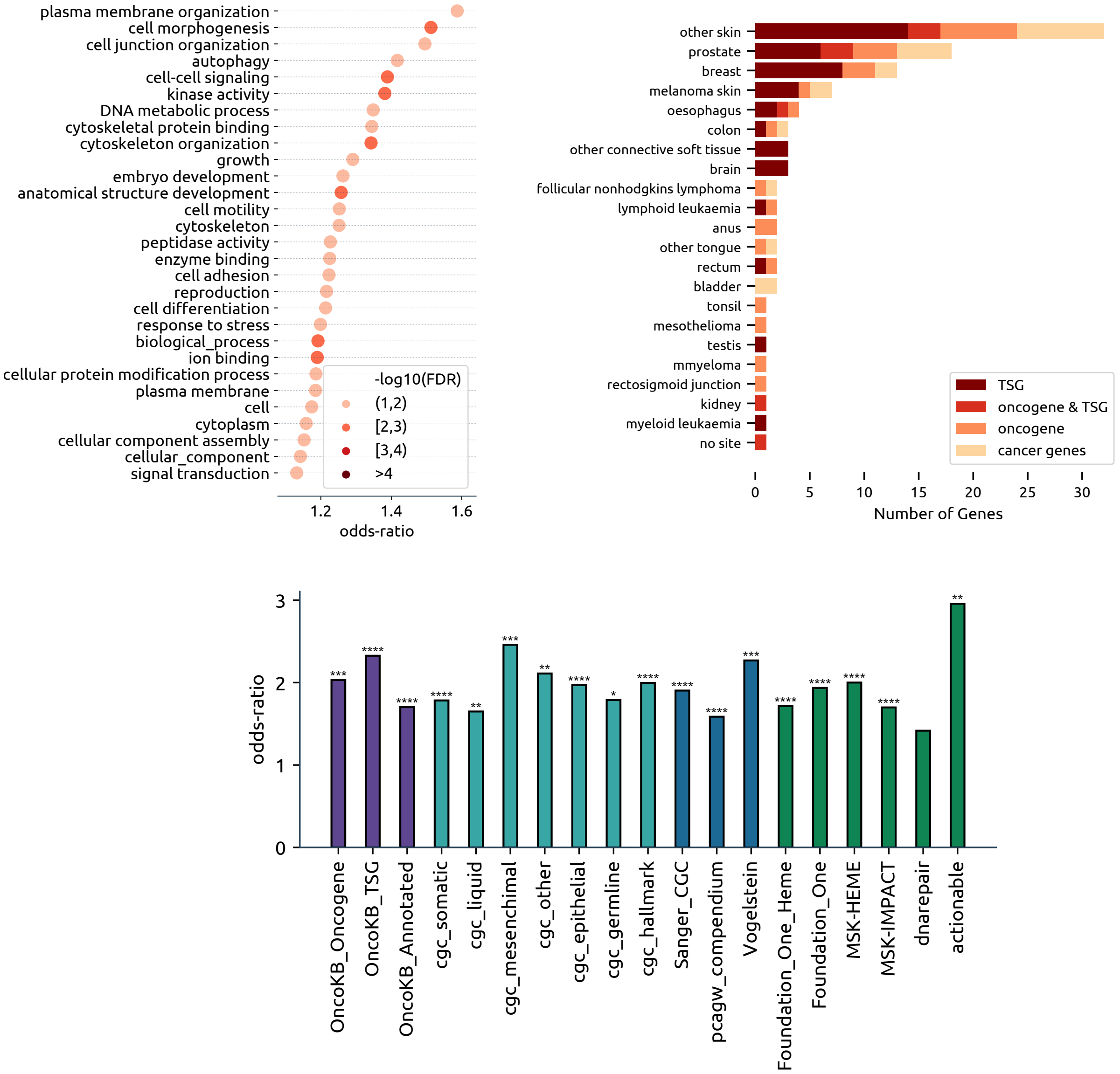

Supplementary Figure 53: **Functional Characterisation of the cancer heritability genes**. A) Gene Ontology terms that are significant in the pathway analysis. For each significant term the point plot represents the odds-ratio while with colours we denote statistical significance, after FDR correction. B) OncoKB annotation for each cancer. Most genes are tumor suppressors, while some of them (denoted as cancer genes in light shade) are known to be drivers but their specific role is not reported. C) CHGs have been tested for association with cancer-related genesets obtained by OncoKB (purple), Cosmic database (light blue), different cancer driver sets (dark blue) and other sets (green) like DNA repair genes and known actionable targets. Stars indicate statistical significance, with multiple terms having $P < 10^{-4}$.

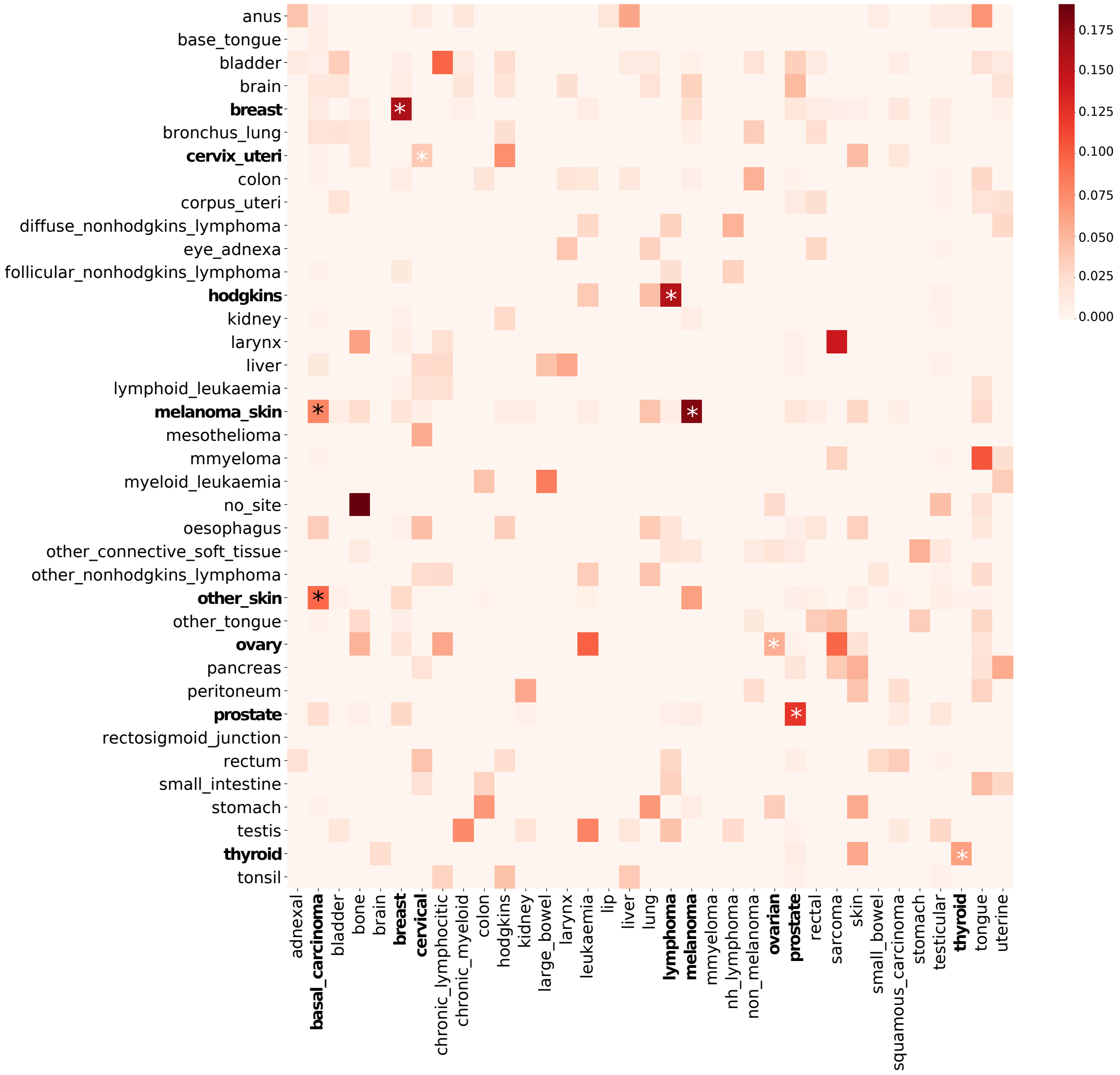

Supplementary Figure 54: **Jaccard Similarity of the significant genes between the ICD10 datasets and the self-reported.** We show a heatmap of the Jaccard Similarity between the significant genes of the histologically characterised dataset, y-axis, and the self-reported one, x-axis. Dark colours correspond to larger similarity. In bold and with white stars we have highlighted high similarities for the same tumour type, while with the dark stars we have highlighted the similarity between different skin-cancer types.

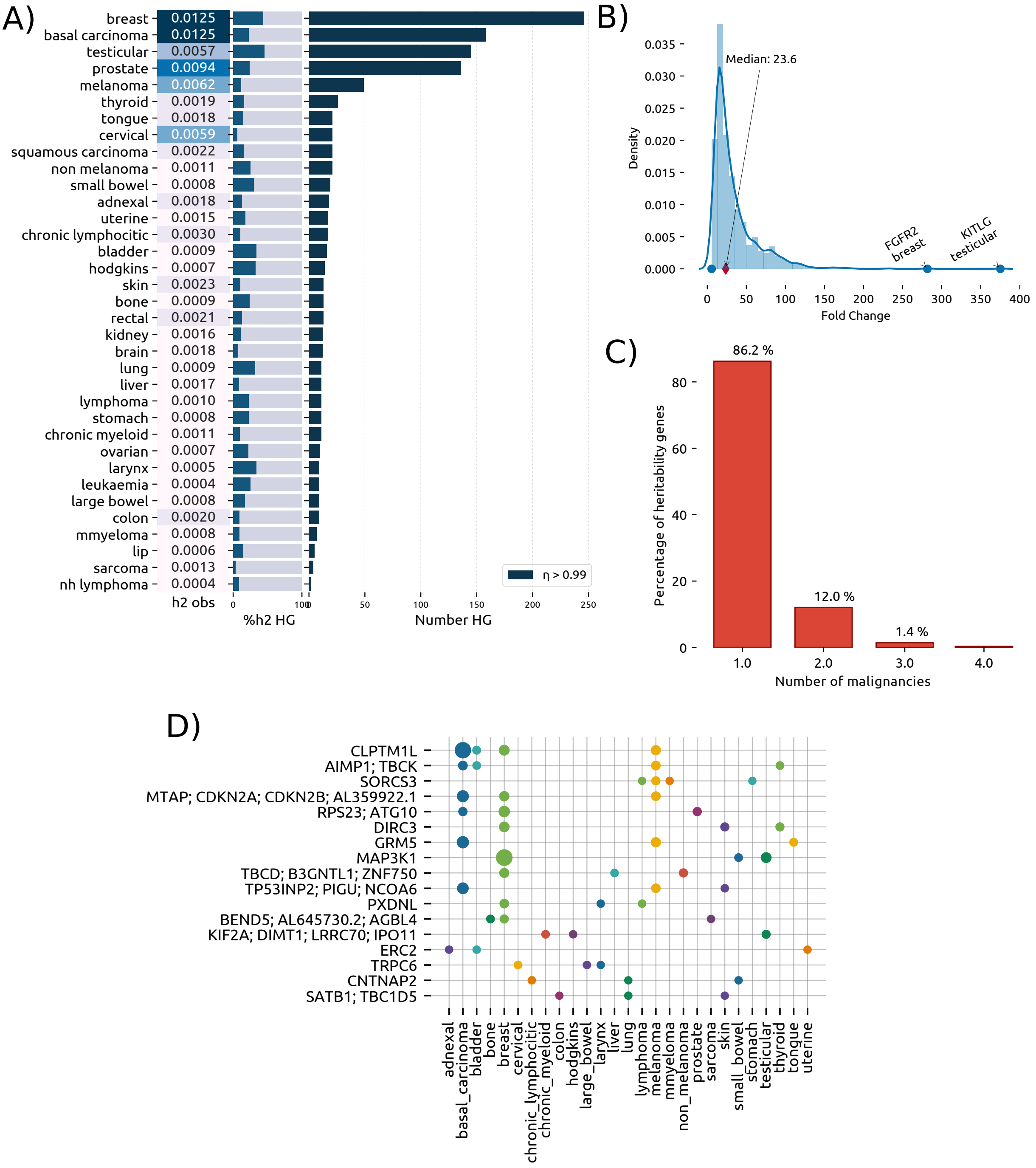

Supplementary Figure 55: **Heritability genes across** $35$ **self-reported cancers in the UK Biobank** A) For each malignancy, we report the observed heritability ($h^2_{SNP}$, left box), the percentage of $h^2_{SNP}$ explained by heritability genes (central barplot, dark blue is the percentage explained by HGs) and the number of heritability genes (right barplot). B) Gene-level heritability density distribution across heritability genes, expressed as fold-change with respect to the genome-wide estimate. Highlighted are the top genes and the median fold-change across all cancers. C) Percentage of cancer heritability genes associated with multiple cancers. More than $10\%$ of CHGs are common to multiple malignancies. D) Cancer heritability genes associated with multiple cancers. We report the $32$ CHGs common to at least 2 cancers; here the size of the dot is proportional to the heritability enrichment of the gene in the specific cancer.

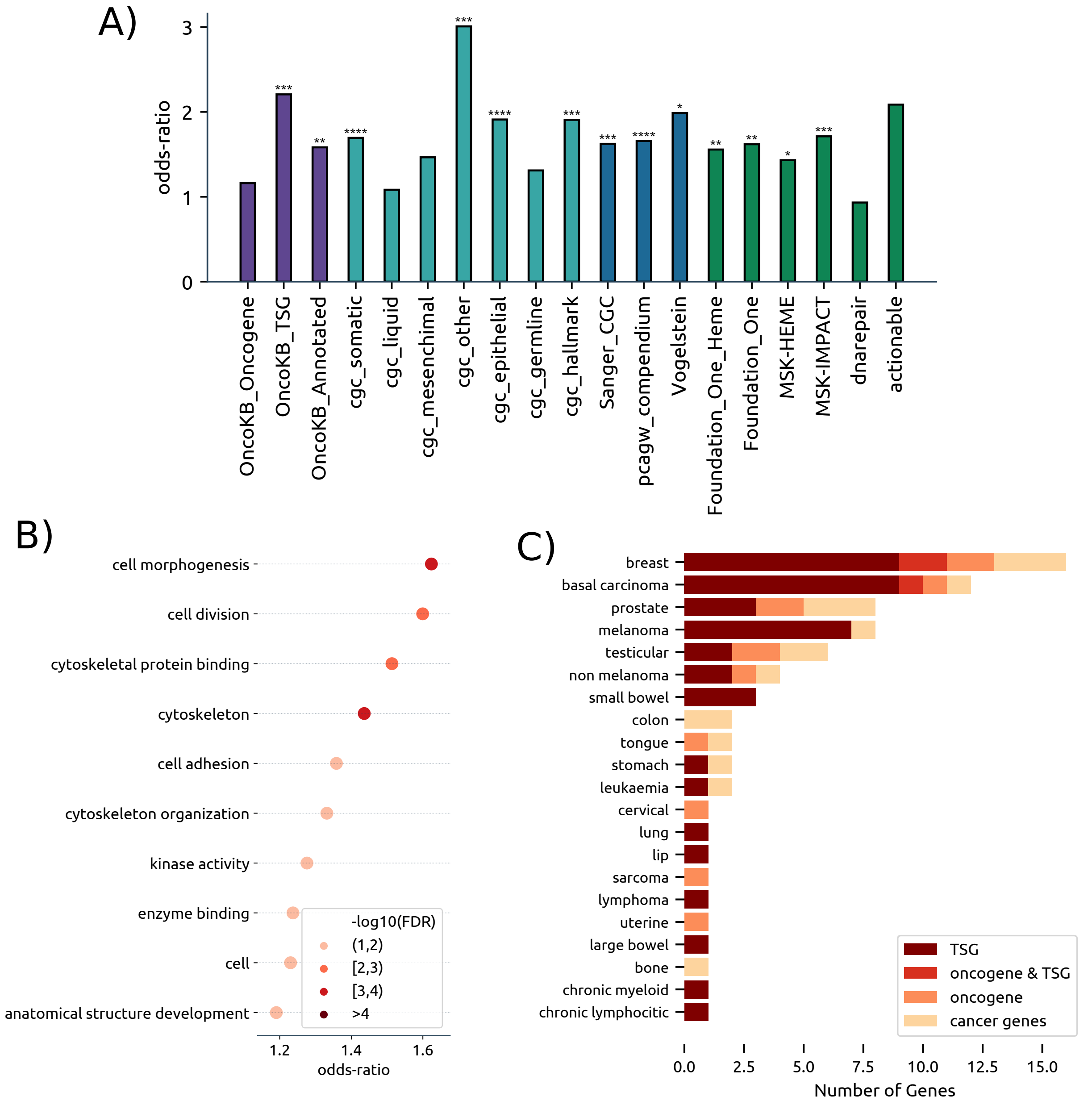

Supplementary Figure 56: **Functional Characterisation of the cancer heritability genes for self reported cancers.** A) CHGs have been tested for association with cancer-related genesets obtained by OncoKB (purple), Cosmic database (light blue), different cancer driver sets (dark blue) and other sets (green) like dna repair genes and known actionable targets. Stars indicate statistical significance, with multiple terms having pvalues $< 10^{-4}$. B) Gene Ontology terms that are significant in the pathway analysis. For each significant term the point plot represents the odds-ratio while with colours we denote statistical significance, after FDR correction. C) OncoKB annotation for each cancer. Most genes are tumor suppressors, while some of them (denoted as cancer genes in light shade) are known to be drivers but their specific role is not reported.

### 5 Supplementary Tables

| Pathway | descriptor | Path. Length | TP | OR | pvalue | FDR |
|---|---|---|---|---|---|---|
| anatomical structure development | GO:0048856 | 4094 | 352 | 1.313969 | 0.000044 | 0.006133 |
| kinase activity | GO:0016301 | 1291 | 126 | 1.442725 | 0.000237 | 0.012169 |
| growth | GO:0040007 | 797 | 84 | 1.558321 | 0.000263 | 0.012169 |
| DNA metabolic process | GO:0006259 | 789 | 82 | 1.531672 | 0.000481 | 0.016723 |
| cytoskeleton organization | GO:0007010 | 1260 | 120 | 1.397913 | 0.000861 | 0.023924 |
| ion binding | GO:0043167 | 5328 | 431 | 1.222055 | 0.001248 | 0.028903 |
| biosynthetic process | GO:0009058 | 4434 | 361 | 1.211568 | 0.002711 | 0.041872 |
| biological_process | GO:0008150 | 6375 | 505 | 1.203162 | 0.002224 | 0.041872 |
| cell morphogenesis | GO:0000902 | 822 | 81 | 1.438235 | 0.002419 | 0.041872 |
| cell proliferation | GO:0008283 | 1636 | 146 | 1.300766 | 0.003404 | 0.047312 |
| cytoskeleton | GO:0005856 | 1597 | 141 | 1.282098 | 0.005851 | 0.054216 |
| cellular protein modification process | GO:0006464 | 3321 | 275 | 1.216768 | 0.004476 | 0.054216 |
| cell-cell signaling | GO:0007267 | 1364 | 123 | 1.309693 | 0.005097 | 0.054216 |
| peptidase activity | GO:0008233 | 1118 | 103 | 1.336823 | 0.005513 | 0.054216 |
| DNA binding transcription factor activity | GO:0003700 | 1832 | 160 | 1.270744 | 0.005068 | 0.054216 |
| enzyme binding | GO:0019899 | 2076 | 178 | 1.246774 | 0.006568 | 0.057059 |
| cell differentiation | GO:0030154 | 3263 | 268 | 1.201015 | 0.007776 | 0.063577 |
| embryo development | GO:0009790 | 818 | 77 | 1.361579 | 0.009437 | 0.069042 |
| cytoskeletal protein binding | GO:0008092 | 817 | 77 | 1.363526 | 0.009173 | 0.069042 |
| nucleus | GO:0005634 | 4363 | 347 | 1.164549 | 0.014507 | 0.097916 |
| DNA binding | GO:0003677 | 2069 | 174 | 1.215415 | 0.014793 | 0.097916 |

Supplementary Table 1: **Significant Gene Ontology** Results of the pathway analysis with the Gene Ontology terms with the heritability genes of the curated 16 datasets. This table corresponds to the results in Figure 3 in the main text

| Dataset | OR | CHG in dataset | pvalue |
|---|---|---|---|
| actionable | 2.63453493776791 | 7 | 0.026951610993734 |
| OncoKB_TSG | 2.4758427927671 | 27 | 7.90E-05 |
| cgc_mesenchimal | 2.24609098939929 | 14 | 0.007835265509946 |
| MSK-HEME | 2.19714313105167 | 48 | 3.93E-06 |
| cgc_other | 2.07244104690334 | 11 | 0.027306118314416 |
| cgc_hallmark | 2.06286703907705 | 33 | 0.00030018959537 |
| Foundation_One | 1.93993932601498 | 37 | 0.000393887476418 |
| Foundation_One_Heme | 1.83497871569604 | 60 | 3.91E-05 |
| OncoKB_Oncogene | 1.83348095659876 | 17 | 0.019840274395826 |
| Vogelstein | 1.83053839364519 | 13 | 0.038349307393117 |
| OncoKB_Annotated | 1.78464447477968 | 52 | 0.000213156056084 |
| MSK-IMPACT | 1.7840487630967 | 47 | 0.000407877043307 |
| cgc_epithelial | 1.75509927797834 | 38 | 0.001711381100092 |
| Sanger_CGC | 1.74637430939227 | 60 | 0.000130077696757 |
| cgc_somatic | 1.70276736998878 | 67 | 0.000110483300951 |
| pcagw_compendium | 1.55039109506619 | 66 | 0.001117467439307 |
| dnarepair | 1.54926413964234 | 15 | 0.080471017287826 |
| cgc_germline | 1.50414250207125 | 10 | 0.151946556078459 |
| cgc_liquid | 1.49944841979726 | 28 | 0.033703719324945 |

Supplementary Table 2: **Cancer Datasets enrichments** Results of the enrichments between the Curated cancer dataset terms and the heritability genes of the curated 16 datasets. This table corresponds to the results in Figure 3 in the main text.

| gene | PS | SG | EIR | CRI | TPI | IM | A | GIM | EPCD | CCE | tsg | oncogene | fusion |
|---|---|---|---|---|---|---|---|---|---|---|---|---|---|
| XPO1 | P | | | | | | | | P | | 0 | 1 | 0 |
| TP63 | P | | | | | P | | | P | | 1 | 1 | 0 |
| SMAD2 | | P | | P | | S | | | S | | 1 | 0 | 0 |
| ROS1 | P | | | | | | | | | | 0 | 1 | 1 |
| RAP1GDS1 | P | | | | | P | | | | | 0 | 1 | 1 |
| RABEP1 | | P | | | | | | | | | 0 | 0 | 1 |
| PPARG | P | P | | | | | | | | P | 1 | 0 | 0 |
| POT1 | | | | | | | | S | | | | 0 | 0 |
| PIK3R1 | | P | | S | | S | | | | | 1 | 0 | 0 |
| PBX1 | | | | | | | P | | P | P | 0 | 1 | 1 |
| PBRM1 | | P | P | | S | S | | S | S | P | 1 | 0 | 0 |
| NT5C2 | P | | | | | | | | P | | 0 | 1 | 0 |
| NCOR2 | | P | | | | | | S | P,S | | 1 | 0 | 0 |
| NAB2 | | | | | | | S | | | | 1 | 0 | 1 |
| MTOR | P | | | | | P | P | | P | P | 0 | 1 | 0 |
| MLLT10 | | | | P | | | | | | | 0 | 1 | 1 |
| LRP1B | | P | | | | S | | | | | 1 | 0 | 0 |
| JAK2 | P | | | | P,S | | | | P | P | 0 | 0 | 0 |
| FOXA1 | P | | | | | S | | | | | 0 | 1 | 0 |
| FGFR2 | P | | | | | | | | P | | 1 | 1 | 0 |
| FAT4 | | P | | | | S | | | | | 1 | 0 | 0 |
| ESR1 | P | P | P | | | P,S | | | | | 1 | 1 | 1 |
| ERBB4 | P | P | | | | | | | P,S | | 1 | 1 | 0 |
| EBF1 | | P | | | | | | | | | 1 | 0 | 1 |
| CTNNB1 | P | P | P | P | | P | P | S | P | P | 0 | 1 | 1 |
| CLIP1 | | | | | | | | | | | 0 | 0 | 1 |
| CIITA | | | S | | | | | | | | 1 | 0 | 1 |
| CDKN2A | | P | | | | S | S | | S | | 1 | 0 | 0 |
| CDH11 | | | | | | S | | | S | | 1 | 0 | 0 |
| CCDC6 | | P | | | | | | S | S | P | 1 | 0 | 1 |
| CBFA2T3 | | P | | | | | | | | P | 1 | 0 | 1 |
| ALK | P | | | | | P | | | P,S | | 0 | 1 | 1 |
| LATS2 | | P | | | | P,S | | S | S | | 1 | 0 | 0 |

Supplementary Table 3: **Hallmarks of Cancer overlaps** Overlap between the hallmark of cancer dataset terms and the heritability genes of the curated 16 datasets. Each column corresponds to one of the hallmarks. P stands for promotes, S stands for suppresses. We also report whether the gene has known, TSG, oncogene or fusion function. This table corresponds to the results in Figure 3 in the main text. **PS**:proliferative signalling, **SG**:suppression of growth, **EIR**:escaping immunic response to cancer, **CRI**:cell replicative immortality, **TPI**:tumour promoting inflammation, **IM**:invasion and metastasis, **A**:angiogenesis, **GIM**:genome instability and mutations, **EPCD**:escaping programmed cell death, **CCE**:change of cellular energetics.

| Phenotype | Complete name | Significant SNPs | minSNPs | minSNP ∩ HG | n_sig |
|---|---|---|---|---|---|
| C44 | Other malignant neoplasms of skin | 580 | 58 | 55 | 422 |
| C50 | Malignant neoplasm of breast | 178 | 10 | 9 | 267 |
| C61 | Malignant neoplasm of prostate | 203 | 20 | 20 | 271 |
| C18 | Malignant neoplasm of colon | 4 | 1 | 1 | 33 |
| C43 | Malignant melanoma of skin | 42 | 14 | 9 | 52 |
| C15 | Malignant neoplasm of oesophagus | 0 | 0 | 0 | 24 |
| C67 | Malignant neoplasm of bladder | 11 | 2 | 1 | 39 |
| C34 | Malignant neoplasm of bronchus and lung | 0 | 0 | 0 | 17 |
| C20 | Malignant neoplasm of rectum | 0 | 0 | 0 | 15 |
| C62 | Malignant neoplasm of testis | 19 | 2 | 1 | 29 |
| C71 | Malignant neoplasm of brain | 0 | 0 | 0 | 19 |
| C45 | Mesothelioma | 1 | 1 | 0 | 5 |
| C91 | Lymphoid leukaemia | 0 | 0 | 0 | 11 |
| C02 | Malignant neoplasm of other and unspecified parts of tongue | 0 | 0 | 0 | 23 |
| C16 | Malignant neoplasm of stomach | 0 | 0 | 0 | 12 |
| C83 | Diffuse non-Hodgkin's lymphoma | 1 | 0 | 0 | 14 |
| C82 | Follicular non-Hodgkin's lymphoma | 0 | 0 | 0 | 21 |
| C90 | Multiple myeloma and malignant plasma cell neoplasms | 0 | 0 | 0 | 15 |
| C56 | Malignant neoplasm of ovary | 0 | 0 | 0 | 13 |
| C54 | Malignant neoplasm of corpus uteri | 0 | 0 | 0 | 14 |
| C48 | Malignant neoplasm of retroperitoneum and peritoneum | 0 | 0 | 0 | 5 |
| C64 | Malignant neoplasm of kidney except renal pelvis | 0 | 0 | 0 | 10 |
| C01 | Malignant neoplasm of base of tongue | 1 | 1 | 0 | 10 |
| C73 | Malignant neoplasm of thyroid gland | 23 | 2 | 2 | 13 |
| C49 | Malignant neoplasm of other connective and soft tissue | 1 | 1 | 0 | 28 |
| C80 | Malignant neoplasm without specification of site | 1 | 1 | 0 | 14 |
| C53 | Malignant neoplasm of cervix uteri | 1 | 1 | 0 | 14 |
| C22 | Malignant neoplasm of liver and intrahepatic bile ducts | 5 | 1 | 0 | 7 |
| C21 | Malignant neoplasm of anus and anal canal | 1 | 1 | 0 | 23 |
| C85 | Other and unspecified types of non-Hodgkin's lymphoma | 0 | 0 | 0 | 9 |
| C09 | Malignant neoplasm of tonsil | 1 | 1 | 0 | 5 |
| C92 | Myeloid leukaemia | 0 | 0 | 0 | 9 |
| C17 | Malignant neoplasm of small intestine | 0 | 0 | 0 | 12 |
| C19 | Malignant neoplasm of rectosigmoid junction | 1 | 1 | 0 | 10 |
| C25 | Malignant neoplasm of pancreas | 0 | 0 | 0 | 12 |
| C81 | Hodgkin's disease | 6 | 1 | 0 | 5 |
| C69 | Malignant neoplasm of eye and adnexa | 0 | 0 | 0 | 14 |
| C32 | Malignant neoplasm of larynx | 1 | 0 | 0 | 7 |

Supplementary Table 4: **Comparison between GWAS raw results and BAGHERA.** For each cancer type we report the number of significant SNPs originally in the dataset, the number of genes that harbor at least a significant SNP (minSNSP), the number of heritability genes, and the overlap between minSNP and HG.

| **UKBB code** | **Malignancy** | **cases** | **prevalence** | $\hat{\chi}^2$ | $h^2_{SNP}$ | $h^2_{SNP_L}$ | **HG** |
|---|---|---|---|---|---|---|---|
| 20001_1002 | **breast cancer** | 7480 | 0.02219 | 1.08192 | 0.01245 | 0.09668 | 246 |
| 20001_1061 | **basal cell carcinoma** | 3156 | 0.00936 | 1.06533 | 0.01250 | 0.18314 | 158 |
| 20001_1044 | **prostate cancer** | 2495 | 0.00740 | 1.05405 | 0.00939 | 0.16460 | 136 |
| 20001_1045 | **testicular cancer** | 614 | 0.00182 | 1.03105 | 0.00567 | 0.30420 | 145 |
| 20001_1059 | **malignant melanoma** | 2677 | 0.00794 | 1.02615 | 0.00622 | 0.10342 | 49 |
| 20001_1041 | **cervical cancer** | 1347 | 0.00400 | 1.02078 | 0.00590 | 0.16776 | 21 |
| 20001_1022 | **colon cancer/sigmoid cancer** | 1134 | 0.00336 | 1.01659 | 0.00196 | 0.06403 | 9 |
| 20001_1040 | **uterine/endometrial cancer** | 843 | 0.00250 | 1.01499 | 0.00148 | 0.06127 | 17 |
| 20001_1062 | **squamous cell carcinoma** | 404 | 0.00120 | 1.01276 | 0.00225 | 0.17012 | 21 |
| 20001_1065 | **thyroid cancer** | 317 | 0.00094 | 1.01245 | 0.00195 | 0.18077 | 26 |
| 20001_1023 | **rectal cancer** | 253 | 0.00075 | 1.01187 | 0.00213 | 0.23923 | 13 |
| 20001_1034 | kidney/renal cell cancer | 436 | 0.00129 | 1.00968 | 0.00156 | 0.11121 | 12 |
| 20001_1035 | bladder cancer | 799 | 0.00237 | 1.00685 | 0.00091 | 0.03954 | 16 |
| 20001_1003 | skin cancer | 1046 | 0.00310 | 1.00679 | 0.00226 | 0.07854 | 13 |
| 20001_1019 | small intestine/small bowel cancer | 156 | 0.00046 | 1.00618 | 0.00076 | 0.12919 | 19 |
| 20001_1030 | eye and/or adnexal cancer | 102 | 0.00030 | 1.00408 | 0.00184 | 0.44827 | 18 |
| 20001_1052 | hodgkins lymphoma / hodgkins disease | 331 | 0.00098 | 1.00324 | 0.00067 | 0.06010 | 14 |
| 20001_1047 | lymphoma | 92 | 0.00027 | 1.00229 | 0.00101 | 0.26830 | 11 |
| 20001_1063 | primary bone cancer | 105 | 0.00031 | 1.00193 | 0.00090 | 0.21425 | 13 |
| 20001_1053 | non-hodgkins lymphoma | 631 | 0.00187 | 1.00082 | 0.00043 | 0.02267 | 2 |
| 20001_1060 | non-melanoma skin cancer | 507 | 0.00150 | 1.00076 | 0.00109 | 0.06863 | 21 |
| 20001_1018 | stomach cancer | 121 | 0.00036 | 0.99947 | 0.00079 | 0.16616 | 11 |
| 20001_1068 | sarcoma/fibrosarcoma | 181 | 0.00054 | 0.99930 | 0.00126 | 0.18758 | 4 |
| 20001_1011 | tongue cancer | 115 | 0.00034 | 0.99905 | 0.00181 | 0.39809 | 21 |
| 20001_1006 | larynx/throat cancer | 250 | 0.00074 | 0.99786 | 0.00052 | 0.05865 | 9 |
| 20001_1004 | cancer of lip/mouth/pharynx/oral cavity | 78 | 0.00023 | 0.99756 | 0.00060 | 0.18505 | 5 |
| 20001_1039 | ovarian cancer | 579 | 0.00172 | 0.99745 | 0.00069 | 0.03903 | 10 |
| 20001_1056 | chronic myeloid | 85 | 0.00025 | 0.99734 | 0.00112 | 0.32044 | 11 |
| 20001_1032 | brain cancer / primary malignant brain tumour | 155 | 0.00046 | 0.99648 | 0.00177 | 0.30057 | 12 |
| 20001_1048 | leukaemia | 158 | 0.00047 | 0.99611 | 0.00045 | 0.07506 | 9 |
| 20001_1024 | liver/hepatocellular cancer | 125 | 0.00037 | 0.99530 | 0.00168 | 0.34389 | 11 |
| 20001_1020 | large bowel cancer/colorectal cancer | 475 | 0.00141 | 0.99524 | 0.00077 | 0.05125 | 9 |
| 20001_1001 | lung cancer | 190 | 0.00056 | 0.99519 | 0.00091 | 0.13020 | 11 |
| 20001_1050 | multiple myeloma | 115 | 0.00034 | 0.99491 | 0.00083 | 0.18195 | 7 |

Supplementary Table 5: **Self reported data.** Here we report a summary of the data and results on the self reported cancer types that were in the first round of GWAS analysis on the UK Bio Bank. Out of $337,159$ total samples, we report the number of cases, the replicative prevalence, and average $\chi^2$ for each cancer type. We also report estimates of heritability, both on the observed $h^2_{SNP}$ and the liability scale $h^2_{SNP_L}$ and the number of heritability genes (HG). Both prevalence and $\hat{\chi}^2$ are lower than the data we used in the paper, here there are only $11$ datasets with $\hat{\chi}^2 > 1.01$.

| Genes | chrom | No SNPs | No Cancers | Cancers |
|---|---|---|---|---|
| CLPTM1L | 5 | 27 | 4 | prostate, melanoma skin, bladder,bronchus lung |
| THADA | 2 | 165 | 4 | prostate, melanoma skin,bladder, diffuse nonhodgkins lymphoma |
| APAF1; ANKS1B; FAM71C | 12 | 582 | 3 | oesophagus, testis, stomach |
| MTRNR2L5; PCDH15 | 10 | 978 | 3 | breast, mesothelioma, lymphoid_leukaemia |
| AGBL1 | 15 | 698 | 3 | testis, diffuse nonhodgkins lymphoma, follicular nonhodgkins lymphoma |
| POU5F1B | 8 | 137 | 3 | breast, prostate, colon |
| ZNF385D | 3 | 862 | 3 | prostate, testis, follicular nonhodgkins lymphoma |
| DLG2 | 11 | 1014 | 3 | oesophagus, bladder, bronchus lung |

Supplementary Table 6: **Cancer Heritability Genes.** Genes common to more than 2 malignancies. These results are obtained from the curated 16 datasets in the main paper. This table corresponds to the top hits of Figure 3D

| Genes | chrom | No SNPs | No cancers | Cancer name |
|---|---|---|---|---|
| CLPTM1L | 5 | 27 | 5 | melanoma skin, prostate, other skin, bronchus lung, bladder |
| MUC19 | 12 | 183 | 5 | thyroid, myeloma, breast, anus, rectosigmoid junction |
| MTRNR2L5; PCDH15 | 10 | 978 | 4 | lymphoid leukaemia, mesothelioma, eye adnexa, breast |
| AUTS2 | 7 | 489 | 4 | oesophagus, lymphoid leukaemia, other nonhodgkins lymphoma, pancreas |
| DPYD | 1 | 574 | 4 | liver, ovary, tonsil, larynx |
| THADA | 2 | 165 | 4 | melanoma skin, prostate, diffuse nonhodgkins lymphoma, bladder |
| KCNS2; STK3 | 8 | 188 | 4 | melanoma skin, small intestine, no site, anus |
| CDH13 | 16 | 1502 | 3 | corpus uteri, melanoma skin, rectosigmoid junction |
| PACRG; PRKN | 6 | 1353 | 3 | thyroid, oesophagus, pancreas |
| NIPAL3; STPG1; GRHL3 | 1 | 136 | 3 | melanoma skin, prostate, other connective soft tissue |
| CLEC16A | 16 | 170 | 3 | other nonhodgkins lymphoma, ovary, diffuse nonhodgkins lymphoma |
| MAST4 | 5 | 383 | 3 | peritoneum, other skin, breast |
| DLG2 | 11 | 1014 | 3 | oesophagus, bronchus lung, bladder |
| APAF1; ANKS1B; FAM71C | 12 | 582 | 3 | testis, oesophagus, stomach |
| SMAP1; B3GAT2 | 6 | 162 | 3 | rectum, other connective soft tissue, colon |
| AGBL1 | 15 | 698 | 3 | testis, diffuse nonhodgkins lymphoma, follicular nonhodgkins lymphoma |
| AGBL4; BEND5; AL645730.2 | 1 | 475 | 3 | ovary, larynx, breast |
| TP53INP2; PIGU; NCOA6 | 20 | 106 | 3 | melanoma skin, other skin, breast |
| GRM5 | 11 | 313 | 3 | melanoma skin, other skin, colon |
| ZFHX4 | 8 | 116 | 3 | melanoma skin, prostate, other skin |
| RERE | 1 | 141 | 3 | kidney, other skin, diffuse nonhodgkins lymphoma |
| CDH4 | 20 | 540 | 3 | testis, prostate, other skin |
| VGLL4; ATG7 | 3 | 245 | 3 | other skin, eye adnexa, other tongue |
| NYAP2 | 2 | 154 | 3 | other skin, other connective soft tissue, breast |
| MTAP; AL359922.1; CDKN2B; CDKN2A | 9 | 162 | 3 | melanoma skin, other skin, brain |
| BACH2 | 6 | 215 | 3 | other skin, other connective soft tissue, breast |
| PREX1 | 20 | 190 | 3 | testis, tonsil, colon |
| GALK2; FGF7; FAM227B; COPS2 | 15 | 157 | 3 | ovary, follicular nonhodgkins lymphoma, breast |
| SEMA3A | 7 | 287 | 3 | peritoneum, other skin, ovary |
| ZNF385D | 3 | 862 | 3 | testis, prostate, follicular nonhodgkins lymphoma |
| POU5F1B | 8 | 137 | 3 | prostate, breast, colon |

Supplementary Table 7: **Cancer Heritability Genes for all 38 malignancies.** Genes common to more than 2 malignancies. These results are obtained from all datasets

| Pathway | descriptor | Path. length | TP | OR | pvalue | FDR |
|---|---|---|---|---|---|---|
| cell morphogenesis | GO:0000902 | 822 | 140 | 1.51249 | 0.00002 | 0.00145 |
| cell-cell signaling | GO:0007267 | 1364 | 215 | 1.38895 | 0.00003 | 0.00145 |
| anatomical structure development | GO:0048856 | 4094 | 576 | 1.25771 | 0.00002 | 0.00145 |
| kinase activity | GO:0016301 | 1291 | 203 | 1.38162 | 0.00006 | 0.00214 |
| cytoskeleton organization | GO:0007010 | 1260 | 194 | 1.34248 | 0.00029 | 0.00703 |
| biological_process | GO:0008150 | 6375 | 848 | 1.19188 | 0.00030 | 0.00703 |
| ion binding | GO:0043167 | 5328 | 716 | 1.18984 | 0.00045 | 0.00900 |
| cell differentiation | GO:0030154 | 3263 | 454 | 1.21364 | 0.00058 | 0.01009 |
| plasma membrane | GO:0005886 | 4994 | 672 | 1.18500 | 0.00069 | 0.01068 |
| response to stress | GO:0006950 | 2975 | 412 | 1.19890 | 0.00164 | 0.02240 |
| cytoskeleton | GO:0005856 | 1597 | 232 | 1.25172 | 0.00210 | 0.02240 |
| cellular protein modification process | GO:0006464 | 3321 | 455 | 1.18618 | 0.00205 | 0.02240 |
| enzyme binding | GO:0019899 | 2076 | 295 | 1.22522 | 0.00199 | 0.02240 |
| DNA metabolic process | GO:0006259 | 789 | 123 | 1.34854 | 0.00244 | 0.02257 |
| cytoskeletal protein binding | GO:0008092 | 817 | 127 | 1.34455 | 0.00231 | 0.02257 |
| cytoplasm | GO:0005737 | 4713 | 628 | 1.15849 | 0.00318 | 0.02763 |
| cell motility | GO:0048870 | 1274 | 186 | 1.25239 | 0.00470 | 0.03845 |
| cellular_component | GO:0005575 | 5314 | 699 | 1.14209 | 0.00581 | 0.04488 |
| growth | GO:0040007 | 797 | 120 | 1.29075 | 0.00852 | 0.06231 |
| signal transduction | GO:0007165 | 5214 | 683 | 1.13158 | 0.00974 | 0.06681 |
| autophagy | GO:0006914 | 379 | 62 | 1.41713 | 0.01009 | 0.06681 |
| cell | GO:0005623 | 2157 | 297 | 1.17421 | 0.01101 | 0.06958 |
| cell adhesion | GO:0007155 | 1149 | 165 | 1.22322 | 0.01378 | 0.07801 |
| peptidase activity | GO:0008233 | 1118 | 161 | 1.22701 | 0.01351 | 0.07801 |
| embryo development | GO:0009790 | 818 | 121 | 1.26283 | 0.01409 | 0.07801 |
| cell junction organization | GO:0034330 | 245 | 42 | 1.49541 | 0.01459 | 0.07801 |
| cellular component assembly | GO:0022607 | 2556 | 346 | 1.15242 | 0.01559 | 0.08027 |
| plasma membrane organization | GO:0007009 | 172 | 31 | 1.58688 | 0.01700 | 0.08150 |
| reproduction | GO:0000003 | 1133 | 162 | 1.21614 | 0.01689 | 0.08150 |

Supplementary Table 8: **Significant Gene Ontology for all malignancies** Results of the pathway analysis with the Gene Ontology terms with the heritability genes of all datasets. This table corresponds to the results in Supp. Figure 56

| Dataset | CHGs in dataset | OR | pvalue |
|---|---|---|---|
| actionable | 12 | 2.95704402853006 | 0.003010513617533 |
| OncoKB_Annotated | 82 | 1.70182693656355 | 3.45E-05 |
| OncoKB_Oncogene | 30 | 2.03015313527443 | 0.000989619358728 |
| OncoKB_TSG | 41 | 2.32559883961873 | 1.10E-05 |
| MSK-IMPACT | 74 | 1.69855042892001 | 8.20E-05 |
| MSK-HEME | 72 | 2.00040589657017 | 1.04E-06 |
| Foundation_One | 60 | 1.93523581681476 | 1.70E-05 |
| Foundation_One_Heme | 93 | 1.71410442349529 | 8.99E-06 |
| Vogelstein | 25 | 2.26853809360218 | 0.000688378926034 |
| Sanger_CGC | 105 | 1.90314876984706 | 4.42E-08 |
| cgc_hallmark | 52 | 1.99517925729025 | 2.98E-05 |
| cgc_somatic | 114 | 1.78333561882259 | 2.14E-07 |
| cgc_germline | 19 | 1.7869406867846 | 0.021626151797484 |
| cgc_epithelial | 68 | 1.96978537106247 | 3.10E-06 |
| cgc_other | 18 | 2.11038080867497 | 0.006672381597831 |
| cgc_mesenchimal | 24 | 2.45812653699978 | 0.000340751626412 |
| cgc_liquid | 50 | 1.64904739495146 | 0.001689100231574 |
| dnarepair | 23 | 1.41604940491173 | 0.085540295201593 |
| pcagw_compendium | 111 | 1.58551000032207 | 2.59E-05 |

Supplementary Table 9: **Cancer Datasets enrichments of all malignancies** Results of the enrichments between the Curated cancer dataset terms and the heritability genes of all datasets. This table corresponds to the results in Supplementary Figure 56
